# An enhancer:involucrin regulatory module impacts human skin barrier adaptation out-of-Africa and modifies atopic dermatitis risk

**DOI:** 10.1101/816520

**Authors:** Mary Elizabeth Mathyer, Erin A. Brettmann, Alina D. Schmidt, Zane A. Goodwin, Ashley M. Quiggle, Inez Y. Oh, Eric Tycksen, Lisa Zhou, Yeriel D. Estrada, X.F. Colin C. Wong, Simon L.I.J. Denil, Scot A. Matkovich, Avner Shemer, John E. A. Common, Emma Guttman-Yassky, Cristina de Guzman Strong

**Affiliations:** Division of Dermatology, Department of Medicine, Washington University School of Medicine, St. Louis, MO 63110, USA; Center for Pharmacogenomics, Department of Medicine, Washington University School of Medicine, St. Louis, MO 63110, USA; Center for the Study of Itch, Washington University School of Medicine, St. Louis, MO 63110 USA; McDonnell Genome Institute, Washington University School of Medicine, St. Louis, MO 63110, USA; Department of Dermatology, Icahn School of Medicine at Mt. Sinai, New York, NY 10029 USA; Skin Research Institute of Singapore, A*STAR, Singapore 138648, Singapore; Tel-Aviv University, Tel-Aviv, Israel

## Abstract

The genetic modules that contribute to human evolution are poorly understood. We identified positive selection for two independent involucrin (*IVL*) haplotypes in European (CEU) and Asian (JPT/CHB) populations for skin epidermis. CEU *IVL* associated with increased *IVL* and a known epidermal-specific enhancer underwent a recent selective sweep out-of-Africa correlating with increased northern latitude. CRISPR/Cas9 deletion of the mouse enhancer revealed enhancer-mediated *cis* regulation for *Ivl* expression with human population-specific enhancer reporter assays confirming the additive effect. Furthermore, *IVL* enhancer eQTLs associated with decreased *IVL* together with filaggrin loss-of-function variants are enriched in atopic dermatitis cases vs. controls. Together, our enhancer-*IVL* cis regulatory module findings reveal an emerging paradigm for recently evolved traits to impact skin disease risk in contemporary populations.

## INTRODUCTION

Modern humans (*Homo sapiens)* have evolved by adapting to local environments and niches, under constant pressure to provide a fitness advantage for survival^1^. Our current understanding of human evolution has been founded on observable phenotypic changes and the underlying genetic variation. However, whole genome sequencing and downstream methodology have facilitated a more reverse genomics approach, allowing researchers to more accurately pinpoint significant genetic differences and thus reveal additional adaptive traits^2,3^. In particular, the availability of multiple divergent human genomes has revolutionized the field, enabling the identification of loci undergoing selection in different populations and their impact on human health^3,4^. Here we investigate human skin barrier adaptation by examining evolution of the Epidermal Differentiation Complex (EDC) locus. The EDC (located at human Chr1q21, Chr3q in mice) exhibited the highest rate of non-synonymous substitutions in the human genome and was ranked as the most rapidly diverging gene cluster in the human-chimp genome comparison^5^. The EDC spans approximately 1.6 Mb and contains 64 genes representing four gene families, including the Filaggrin (*FLG*)-like or SFTP (S100 fused type protein), Late Cornified Envelope (LCE), Small Proline Repeat-Rich (SPRR), and S100-domain (S100) family members^6,7^. The expression of many of these EDC genes is a hallmark feature of the terminally differentiated epidermal cells (keratinocytes) that comprise the interfollicular epidermis at the skin surface^8^. Many EDC proteins, including involucrin (IVL) and many SPRRs and LCEs, are covalently cross-linked to form the cornified envelope that surrounds the keratinocyte^8^. Identification of EDC orthologs across many mammalian genomes^9–12^ led to the discovery of the observed linearity and synteny of the EDC in both mammals and vertebrates^6,7,13–17^. We previously identified positive selection in the EDC for specific genes across the mammalian phylogeny, and even specifically in primates and humans^18^. Motivated by these studies, we hypothesized ongoing evolution within the EDC for skin barrier adaptation among the geographically diverse group of modern-day human populations.

## RESULTS

### Population-specific signals of positive selection in the Epidermal Differentiation Complex reveal recent human evolution in the skin barrier with a convergence at involucrin (*IVL*)

We identified positive selection in the EDC using 2 independent algorithms, Composite of Multiple Signals (CMS)^19,20^ and the integrated Selection of Allele Favored by Evolution (iSAFE)^21^ (Fig. 1). CMS scores were previously calculated for HapMap II single nucleotide polymorphisms (SNPs) from Utah Individuals of European descent [CEU], Yoruba from Ibadan, Nigeria [YRI], and pooled Han Chinese from Beijing/Japanese from Tokyo, Japan [JPT/CHB]) populations. We extracted CMS scores for each EDC SNP from each population. Positive selection was only found near *LCE3E* in YRI, as evidenced by a cluster of three SNPs with CMS scores > 0 (Fig. 1b, Table S1). In contrast, we identified three clusters of positive CMS-scoring SNPs in JPT/CHB in the regions of *HRNR-FLG, CRNN-LCE5A*, and *IVL-LINC01527*. The *HRNR-FLG* genomic region exhibited the strongest signal of positive selection for this combined population (0.63≤CMS≤5.23) and is in linkage disequilibrium (LD) (r^2^≥0.8) with a previously reported positively selected *HRNR*-*FLG* “Huxian haplogroup” in the Han Chinese population^22^. CEU harbored four genomic regions with evidence of positive selection: *LCE2D-LCE2C, SMCP*-*IVL, SPRR2E-SPRR2F*, and *S100A6*. The signal of selection at *SMCP*(3.18≤CMS≤8.17) was the strongest signal in CEU and of any population. As *SMCP* encodes a sperm mitochondrial cysteine-rich protein^23^, the finding suggests a role for the positively selected SNPs in sperm function. Importantly, *IVL* was the only gene with evidence of positive selection observed in both JPT/CHB and CEU populations. However, the positively selected SNPs in the *IVL* region were distinct between the two populations (Table S1). Together, our results identify multiple population specific signals of positive selection, with a notable convergence of distinct positive selection at *IVL* in two geographically distinct human populations.

**Figure 1.**
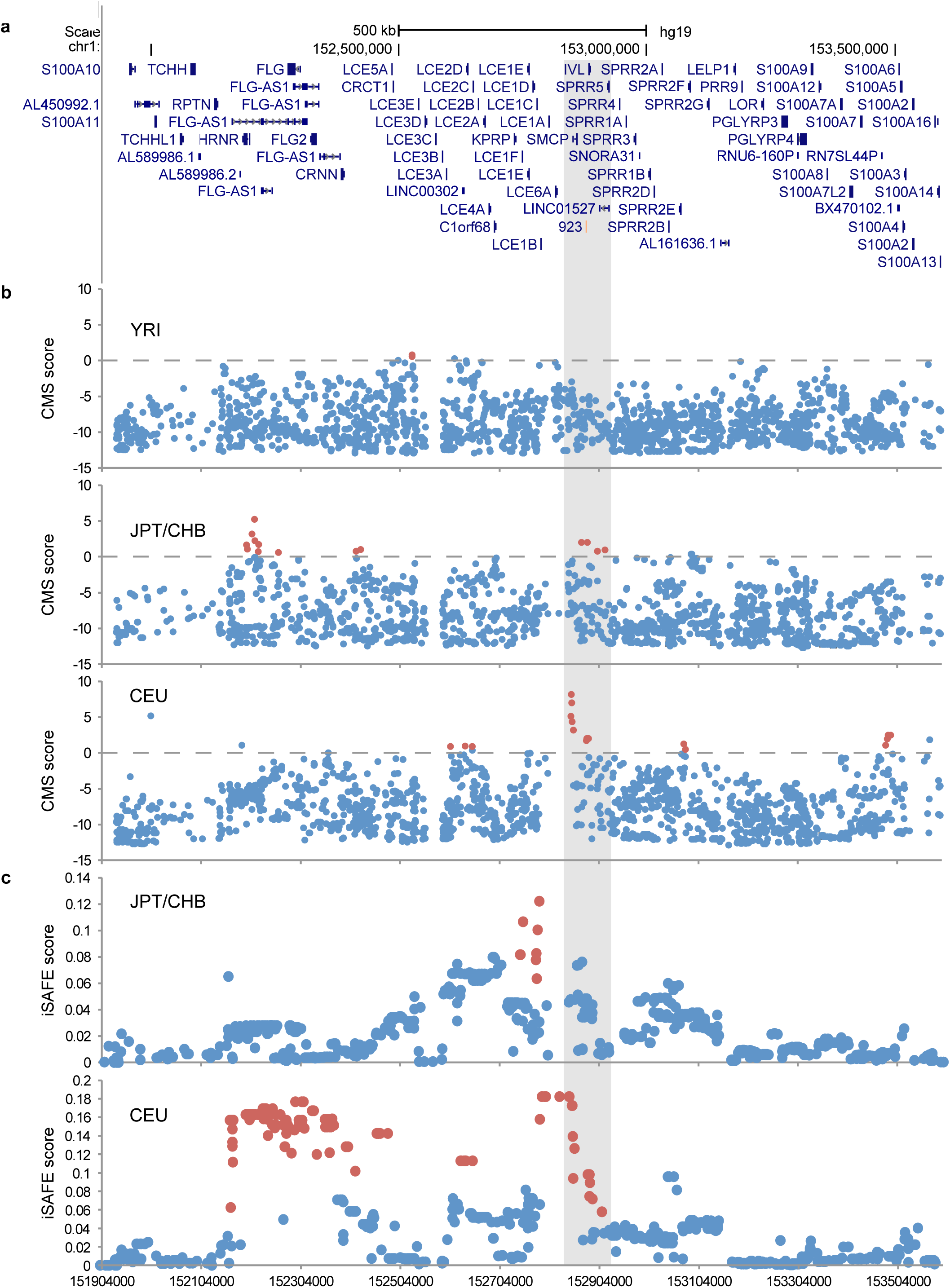
Recent human evolution in the skin barrier as evidenced by multiple population-specific signals for positive selection in the Epidermal Differentiation Complex. Positive selection within the **a**) EDC (hg19; chr1:151,904,000-153,593,700) was determined by **b**) clusters of SNPs with CMS score > 0 (above dashed lines) and **c**) SNPs in iSAFE peaks with maximum peak height > 0.1. High-CMS SNPs and SNPs within iSAFE peaks are shown as red dots. The gray shaded region indicates the region of shared evidence of positive selection near *SMCP-SPRR5.*CMS scores were previously calculated for 1KGP Pilot Phase SNPs in YRI, JPT/CHB, and CEU populations and iSAFE scores were calculated for 1KGP Phase 3 SNPs in JPT/CHB and CEU.

We sought to validate and refine the positively selected regions identified by CMS using iSAFE. iSAFE scores were calculated for 1000 Genomes Project (1KGP) Phase 3 SNPs within the EDC in YRI, JPT/CHB, and CEU populations^21^, with evidence of positive selection defined as a peak height greater than 0.1. We found no evidence of positively selected SNPs using iSAFE in YRI. iSAFE revealed positive selection at *LCE1F-LCE1B* in JPT/CHB that was not identified by CMS (Fig. 1c, Table S2), and did not replicate the CMS findings for *HRNR*-*FLG* and *CRNN-LCE5A* (Fig. 1b). A weak iSAFE peak (peak height .076) between *SMCP* and *IVL* was observed in JPT/CHB but was just below the 0.1 threshold for positive selection. In CEU, iSAFE identified three genomic regions under positive selection that included *HRNR-CRNN, LCE2D*, and *LCE1B-SPRR5* (Fig. 1c, Table S3). The *HRNR-CRNN* signal includes multiple genes in the S100-fused family (*HRNR, FLG, FLG2, CRNN* genes), and the noncoding RNA *FLG-AS1.* Interestingly, neither the *HRNR-CRNN* peak nor the peak at *LCE2D* was identified by CMS. The *LCE1B*-*SPPR5* iSAFE peak region includes the genes *LCE1B, LCE1A, LCE6A, SMCP, IVL, LINC01527*, and *SPRR5*. Strikingly, the *LCE1B-SPRR5* iSAFE peak includes 12 of the 13 SNPs identified by CMS in CEU, further validating positive selection associated with these SNPs. Together, our results identify positive selection at the *SMCP-IVL* loci for both CEU and JPT/CHB using CMS with independent validation for CEU by iSAFE, solidifying the significance of this region for human skin barrier evolution.

We next determined the haplotypes for the positively selected SNPs in the *SMCP-IVL* regions in JPT/CHB and CEU. We identified that positively selected JPT/CHB *SMCP*-*IVL* and CEU-*IVL* haplotypes overlap at *IVL* and represent independent yet shared emergences of positive selection at this locus. Specifically, linkage analysis for the high-CMS SNPs rs6668295 and rs11205130 (r^2^=1.0) revealed a *SMCP-IVL* haplotype (46.5 kb) in JPT/CHB that spans from the *SMCP* promoter to 3’ of *IVL* (Fig. 2a, Table S1). A second, neighboring JPT/CHB-*LINC01527* haplotype (23.0 kb; rs78730108 and rs4240864, r^2^=1.0) is named for and contains a skin-specific long intergenic non-coding RNA^24^. In CEU, eight high-CMS SNPs comprise a distinct CEU-*IVL* haplotype (41.2 kb) that spans from 3’ of *SMCP* to 3’ of *IVL* (0.8146<r^2^<1.0). A neighboring CEU *LCE1A-SMCP* haplotype (65.2 kb) contains four of the five high-CMS SNPs in the region (r^2^=1.0). Notably, the SNPs that comprise the CEU high-iSAFE *LCE1B-SPRR5* peak belong to the two haplotypes defined by the high-CMS SNPs in the region. Together, this analysis identified a convergence of positive selection around *IVL*.

**Figure 2.**
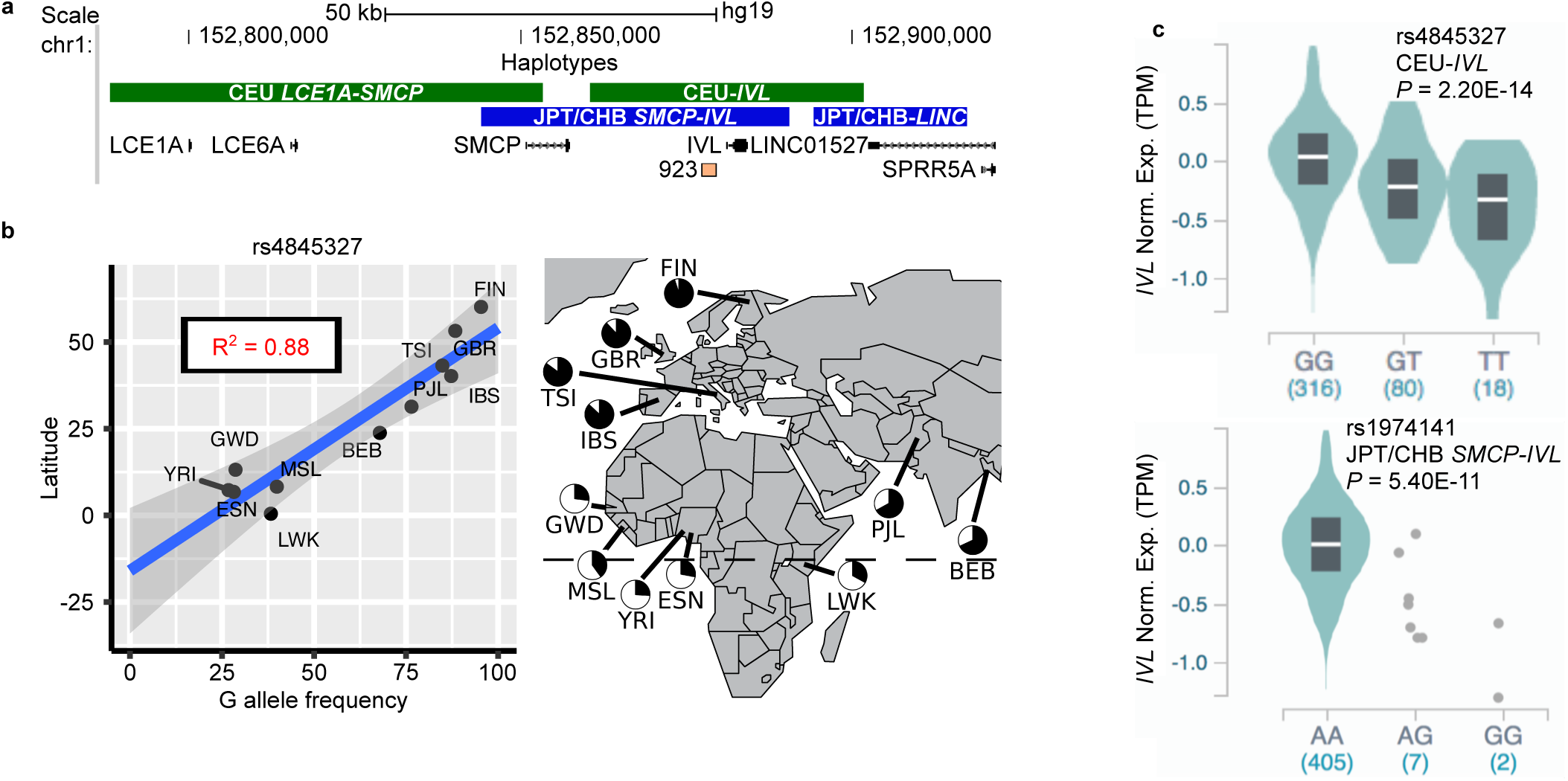
Distinct yet convergent emergences of positively selected *IVL* haplotypes in CEU and JPT/CHB with a recent selective sweep of CEU-*IVL* in Northern latitudes associated with increased *IVL* expression. **a**) Phased SNPs in linkage disequilibrium (r^2^>0.8) with the high CMS SNPs revealed 2 positively selected haplotypes for each of the CEU and JPT/CHB populations, depicted as colored bars (green, CEU *LCE1A*-*SMCP* and CEU-*IVL*; blue, JPT/CHB *SMCP*-*IVL* and JPT/CHB-*LINC01527*). CEU-*IVL* and JPT/CHB *SMCP-IVL* haplotypes share commonality for both *IVL* and the epidermal-specific 923 enhancer (pink). **b**) A direct and positive correlation between the frequency of rs4845327-G, a tagging SNP for the CEU-*IVL* haplotype, and latitude reveals a selective advantage for CEU-*IVL* with increasingly Northern latitudes. The blue line indicates a linear relationship between allele frequency and geographic latitude and the gray area surrounding the regression line represents the 95% confidence intervals of the latitude values for each allele frequency value. Pie charts of allele frequency for each population are indicated on the map. **c**) Violin plots from GTEx of *IVL* expression level by genotype for tagging SNPs for the CEU-*IVL* and JPT/CHB *SMCP-IVL* haplotypes (rs4845327, top, and rs1974141, bottom, respectively). The CEU allele of rs4845327 (G) is associated with significantly increased *IVL* expression whereas the JPT/CHB allele of rs1974141 (G) is associated with significantly reduced *IVL* expression. Numbers in parenthesis indicate sample size for each genotype. TPM, transcripts per million.

Notably, the positively selected CEU-*IVL* haplotype is globally distributed, with the haplotype found in European (AF=0.86), African (AF=0.27), East Asian (AF=0.42), South Asian (AF=0.68), and admixed American (AF=0.67) populations. This suggests that the origin of the CEU-*IVL* haplotype minimally dates prior to the migration out of Africa. We next examined the correlation between the allele frequency for the CEU-*IVL* haplotype proxy SNP, rs4845327-G, for a given 1KGP population with respect to the population’s latitude. We find a direct and positive correlation between the G allele of rs4845327 and northern latitude (*R*^*2*^= 0.95, ρ= 0.88, Fig. 2b). Together, these data reveal a recent selective sweep for the CEU-*IVL* haplotype in European populations at increasingly Northern latitudes.

### CEU-*IVL* haplotype is associated with increased *IVL* gene expression and JPT/CHB *SMCP*-*IVL* haplotype is associated with decreased *IVL* gene expression

We next determined the functional significances of the positively selected CEU-*IVL* and JPT/CHB *SMCP*-*IVL* haplotypes using Genotype-Tissue Expression Project (GTEx)^25^. The CEU-*IVL* haplotype contains expression quantitative trait loci (eQTLs) associated with increased *IVL* expression (Fig. 2c, top; representative SNP rs4845327 shown), while the JPT/CHB *SMCP-IVL* haplotype harbors eQTLs associated with decreased *IVL* expression (Fig. 2c, representative SNP rs1974141 shown). We hypothesize that this differential expression is driven by an enhancer with population-specific variation. We previously identified and characterized a strong epidermal-specific enhancer located upstream of *IVL*^7^ (Fig. 2a, Fig. 3a). This enhancer, termed ‘923,’ is located 923 kb from the most 5’ EDC gene and is associated with the dynamic chromatin remodeling of the EDC and cJun/AP1 binding concomitant with EDC gene expression^26^. Both the CEU-*IVL* and JPT/CHB *SMCP-IVL* haplotypes contain the 923 enhancer and *IVL*. This led us to hypothesize that the 923 enhancer modulates *IVL* expression associated with positive selection.

**Figure 3.**
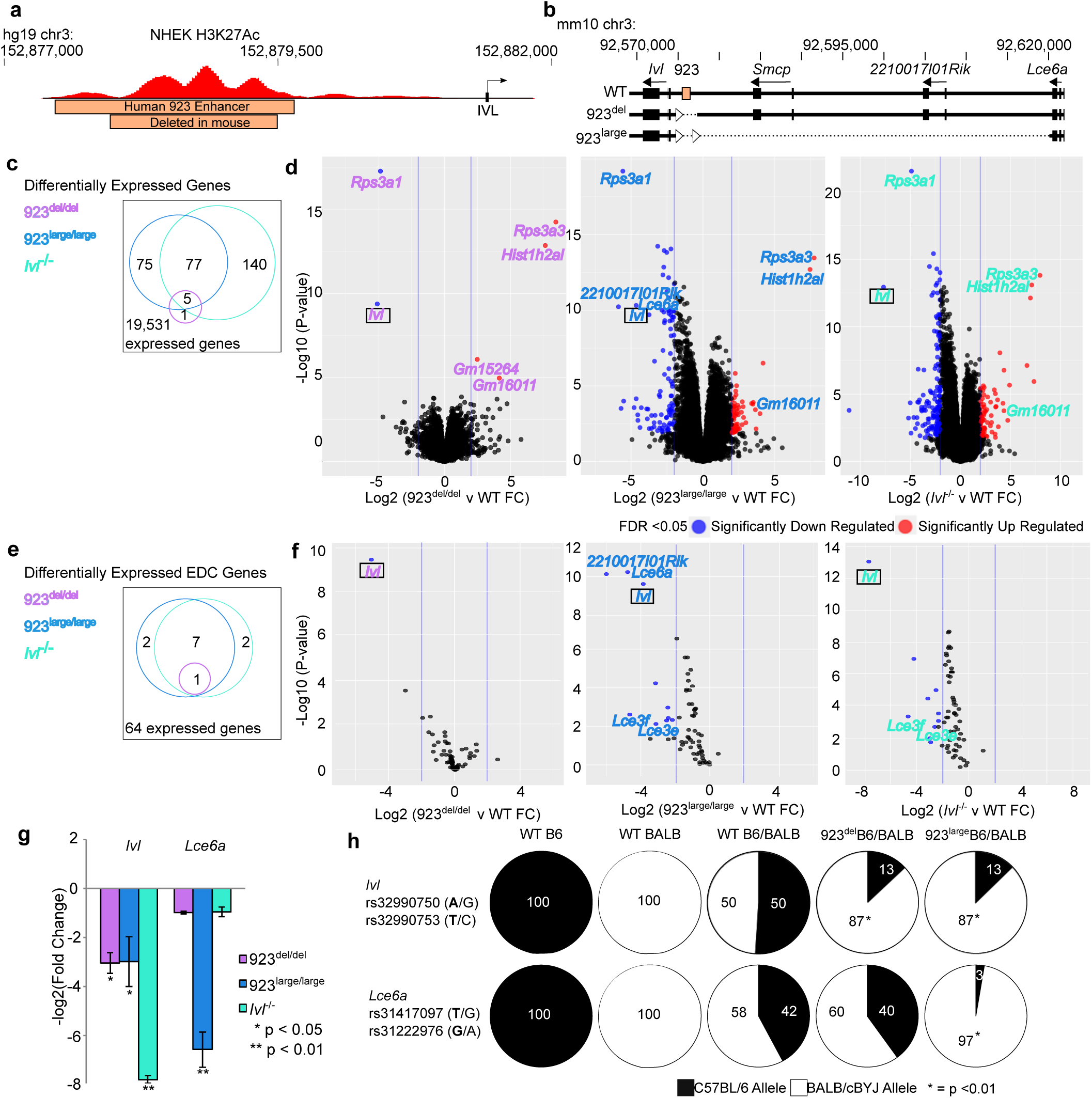
Deletion of the 923 enhancer in vivo identifies involucrin as a target gene via cis-regulation. **a)** Schematic of the human 923 enhancer as defined by H3K27Ac marks in NHEK; hg19; chr1:152,877,349-152,879,610 and the orthologous deleted enhancer sequence in the mouse (orange). **b)** Generation of two 923 enhancer deletion alleles by CRISPR/Cas9 genome editing in mice, 923^del^: Deletion of mouse orthologous 923 enhancer; 923^large^: ∼40kb deletion (mm10: chr3: 90,575,863 – 92,577,075) that includes 923 enhancer, *Smcp*, and *2210017I01Rik* annotations. Intact flanking *loxP* sites (triangles) were also successfully introduced via homologous recombination. **c)** Venn diagram (FDR<0.05, log2(FC)>|2|) and **d**) volcano plots of ribosomal-zero RNA-seq differential gene expression analyses of 923^del/del^, 923^large/large^, and *Ivl*^-/-^ newborn mouse skins each compared to WT reveal significant decreased expressions for *Ivl. n*=3/genotype. **e)** Venn diagram of differentially expressed EDC genes (FDR <0.05, log2(FC)>|2|) and **f)** volcano plots of EDC subset from differential gene expression analyses of 923^del/del^, 923^large/large^, and *Ivl*^-/-^ newborn mouse skins each compared to WT (*n*=3/genotype). **g)** Confirmatory qPCR validations for decreased *Ivl* expression for *Ivl*^-/-^, 923^large/large^ and 923^del/del^ and additional decreased *Lce6a* expression specific to 923^large/large^ newborn mouse skins; *n*=3/genotype; Error bars, ±SEM. **h)** Proportions of allele-specific (SNP) *Ivl* and *Lce6a* expression shown in pie charts for each mouse strain. Two SNPs per allele were measured and showed identical allele frequencies. C57BL/6 SNP bolded, \**P*<0.01, ANOVA compared to WT C57BL/6/BALB/cBYJ mice.

### Deletion of the 923 enhancer in mice results in decreased expression of the proximal gene involucrin in the skin

To determine the function of the 923 enhancer, we generated knockout mice for the orthologous 923 enhancer using CRISPR/Cas9 genome editing (Fig. 3a). Two independent deletions were successfully generated, resulting in a specific deletion of the 923 enhancer (923^del^) and a large 40kb deletion that included the 923 enhancer and proximal genes *Smcp* and *2210017I01Rik* (923^large^) (Fig. 3b, Table S4, Fig. S1). 923^del^ and 923^large^ knockout mice were viable at birth and did not deviate from the expected genotype ratios for both heterozygous parental crosses (^2^ test, α=0.05) (Table S5). We next examined the molecular impact of the 923 enhancer deletion on the skin transcriptome. RNA-seq on 923^del^ and 923^large^ homozygous, heterozygous, and wild-type (WT) newborn skins was performed (Table S6-9). Given the close proximity of the 923 enhancer to involucrin *(Ivl)*, we included the *Ivl* knockout mouse^4^ in our analyses to distinguish the direct effect of the loss of the enhancer from the hypothesized loss of involucrin (Table S10). Analysis of the skin transcriptomes identified 6 significantly differentially expressed genes (DEGs) in 923^del/del^ skins, in contrast to 157 and 222 genes in 923^large/large^ and *Ivl*^-/-^ skins, respectively (FDR < 0.05, log2(FC)≥|2|) (Fig. 3cd). Strikingly, 5 of the 6 DEGs in 923^del/del^ (*Rps3a3, Hist1h2aI, Gm16011, Rsp3a1*, and *Ivl)* were also differentially expressed in the same directions in 923^large/large^ skins (Fig. 3d, Fig. S2). *Gm15264* was only differentially expressed in 923^del/del^ mice. As *Rps3a1, Rps3a3, Hist1h2aI*, and *Gm16011* were also differentially expressed in the same directions in *Ivl*^-/-^ skins, their observed differential expression in the 923 deletion lines are likely secondary effects associated with decreased *Ivl* expression in the two enhancer deletion skins. Together, our *in vivo* results identify a functional role for the 923 enhancer in the regulation of *Ivl* target gene expression.

The effect of the enhancer deletion on the expression of its most proximal gene, *Ivl*, motivated us to further examine local transcriptional effects with respect to only the EDC. RNA-seq analyses of the EDC loci identified one DEG (*Ivl*) in 923^del/del^,10 in 923^large/large^, and 10 in *Ivl*^-/-^ skin (Fig. 3ef) (FDR < 0.05, log2(FC)≥|2|). *Smcp*, the most proximal gene 3’ of 923 (Fig. 3b), was below our detection limits for this assay, consistent with the low numbers of transcripts detected in whole skin scRNA-seq^27^. We validated the significance of decreased *Ivl* expression in all 3 mouse lines via qPCR (ANOVA; Tukey post-hoc *P*<0.05) (Fig. 3g). By contrast, 923^large/large^ mouse skins also exhibited significantly decreased expression of *Lce6a*, the most proximal gene 3’ of the deletion, that was not observed in the other 2 mouse lines. This suggests the presence of an additional as yet uncharacterized enhancer region for *Lce6a* that was also deleted in the 923^large^ allele. Together, both enhancer deletion lines demonstrate a requirement for the enhancer to positively regulate *Ivl* expression and for the 923^large^ allele for *Lce6a* expression.

### 923 enhancer regulates *Ivl* target gene expression in an allele-specific (*cis*) manner

To determine if the 923 enhancer regulates gene expression in *cis*, we assessed allele-specific gene expression for *Ivl* and *Lce6a* in heterozygous 923^del^[C57BL/6]/[BALB/cBYJ] and 923^large^[C57BL/6]/[BALB/cBYJ] mouse skins. The BALB/cBYJ (hereafter, BALB) genetic strain was selected given the presence of single nucleotide polymorphisms (SNPs) in the *Ivl* and *Lce6a* coding sequences that can be distinguished from the C57BL/6 (hereafter, B6) background (923^del^, 923^large^). Targeted sequencing of *Ivl* and *Lce6a* cDNAs obtained from 923^del^B6/BALB and 923^large^B6/BALB heterozygous mouse skins revealed a significantly lower proportion of B6 SNPs from the 923^del^ and 923^large^ alleles for *Ivl* (13%) compared to 50% of B6 SNPs from the control WT B6/BALB mice (Fig. 3h) (ANOVA, Tukey post-hoc, *P*<0.01). However, a significantly lower proportion of B6 SNPs (3%) for *Lce6a* was observed only in the 923^large^B6/BALB heterozygous mice compared to 42% observed in the control B6/BALB WT skins (ANOVA, Tukey post-hoc, *P*<0.01). This further supports the hypothesis of the loss of a regulatory enhancer in the 923^large^ allele that regulates allele-specific *Lce6a* expression. Together, our genetic findings identify *cis* regulation by the 923 enhancer for *Ivl* and additionally for *Lce6a* by the 923^large^ allele.

### Deletion of 923 enhancer affects the chromatin landscape in the EDC

We next sought to determine the effect of the 923 enhancer deletion on the EDC chromatin landscape. Open (or accessible) chromatin in the epidermis of newborn 923^del/del^, 923^large/large^, and WT mice was determined using ATAC-seq (assay for transposase accessible chromatin using sequencing)^28^. A total of 383,971 accessible chromatin regions were identified genome-wide. We next determined differential chromatin accessibility between WT and 923^del/del^ and also 923^large/large^ epidermis. Ten differentially accessible regions (DARs) were identified between 923^del/del^ and WT mice in contrast to 23 DARs found in the 923^large/large^ comparison (Fig. 4a, Table S11-12) (FDR<0.05, log2(FC)≥|2|). Six of these DARs were shared between the 923^del/del^ and 923^large/large^ comparisons (Fig. 4a), including two peaks on chromosome X and four peaks on chromosome 3, three of which are within or near (<250kb) the EDC (Fig. 4b, Fig. S3). Two DARs, intergenic to *Lce3e*/*Lce3f* and corresponding with pseudogene *Gm29950* (intergenic to *S100a11*/*S100a10)*, were less accessible whereas an intergenic DAR at *Tdpoz4*/*Tdpoz3* was more accessible in both 923^del/del^ and 923^large/large^ skins compared to WT. Although none of the DAR proximal genes were significantly differentially expressed in 923^del/del^ skin, 3 of the 23 923^large/large^ DARs correlated with significant changes in 4 genes in 923^large/large^ mice, all located within the EDC (*Ivl, 2210017l01Rik, Lce3e, Lce3f) (*FDR <0.05). (Fig. S4). Thus, we determine that deletion of the 923 enhancer results in less accessible chromatin in the EDC and highlights a functional role to facilitate proximal chromatin accessibility.

**Figure 4.**
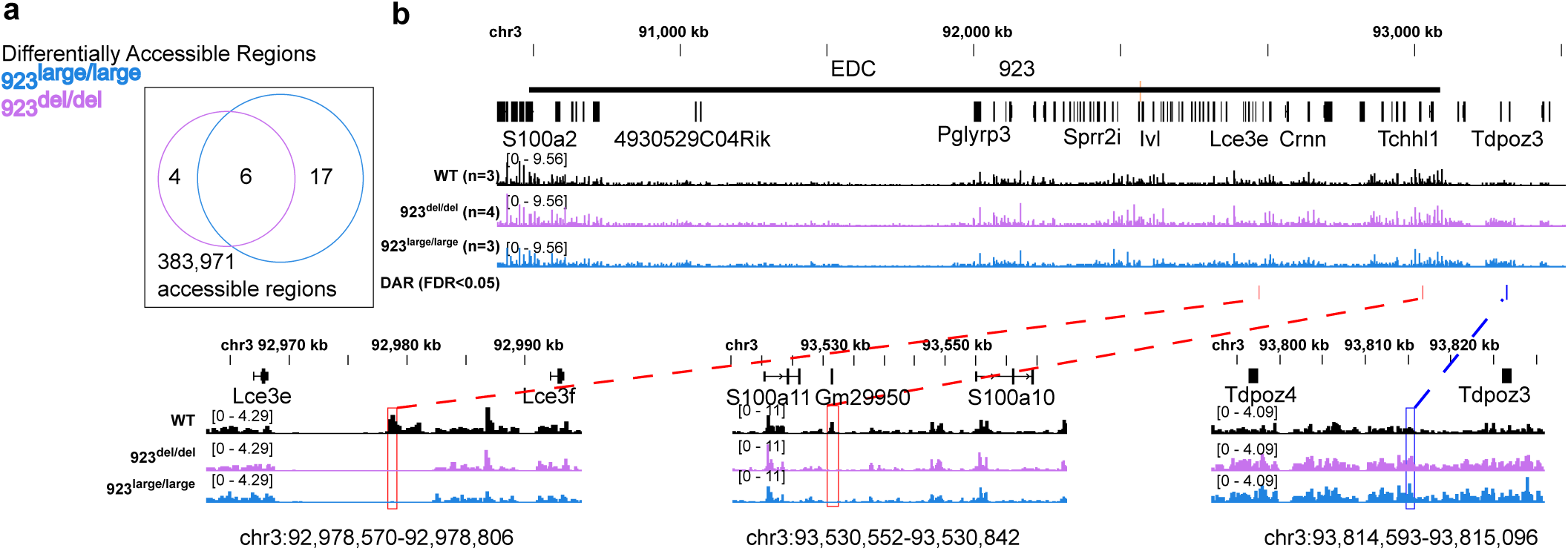
Chromatin accessibility is altered in and around the EDC in 923^del/del^ and 923^large/large^ newborn epidermis. Venn diagram (FDR <0.05) of all differentially accessible regions (DARs) using comparative ATAC-seq in 923^del/del^ and 923^large/large^ newborn epidermis that were each compared to WT with an enrichment of shared DARs in the EDC. Peaks shown as fold change signal per genotype (WT, *n*=3; 923^del/del^, *n*=4; 923^large/large^, *n*=3) with shared less accessible (red) and more accessible (blue) DARs indicated in the 923 deletion lines compared to WT.

### The human 923 sequence enhances expression from the *IVL* promoter and both elements exhibit population-specific regulatory activities

Our discoveries for the positively selected *IVL* human haplotypes associated with differential *IVL* expressions and the 923 enhancer:*Ivl* module in mice led us to investigate the functionality of human population-specific alleles for the 923 enhancer:*IVL* module. We tested population-specific alleles containing the *IVL* promoter, first (noncoding) exon, and intron, which are known to exhibit collective regulatory activity^29^ (Fig. 5a). The CEU *IVL* promoter/intron allele exhibited significantly higher luciferase activity than the JPT/CHB and YRI alleles in both proliferating and differentiated keratinocyte cell culturing conditions (*P*<0.05), consistent with the GTEx annotation for increased *IVL* expression (Fig. 5b, Fig. 2c). We next added the respective population-specific enhancer to its *IVL* promoter/intron allele and found further increased luciferase reporter activities for all population-specific alleles compared to the promoter/intron only alleles, and specifically more so in differentiated cells, where *IVL* is endogenously expressed in skin tissue. However, a comparison between population-specific alleles revealed relatively decreased luciferase activity for the JPT/CHB enhancer/promoter/intron allele compared to CEU and YRI alleles, again consistent with the relatively decreased *IVL* expression associated with this haplotype in GTEx (Fig. 5b, Fig. 2c). Thus, our findings further support the *cis*-regulation of human *IVL* by the enhancer, with the CEU enhancer/*IVL* promoter/intron allele driving increased *IVL* expression and conversely decreased expression in the JPT/CHB allele, both of which we show are under positive selection (Fig. 1).

**Figure 5.**
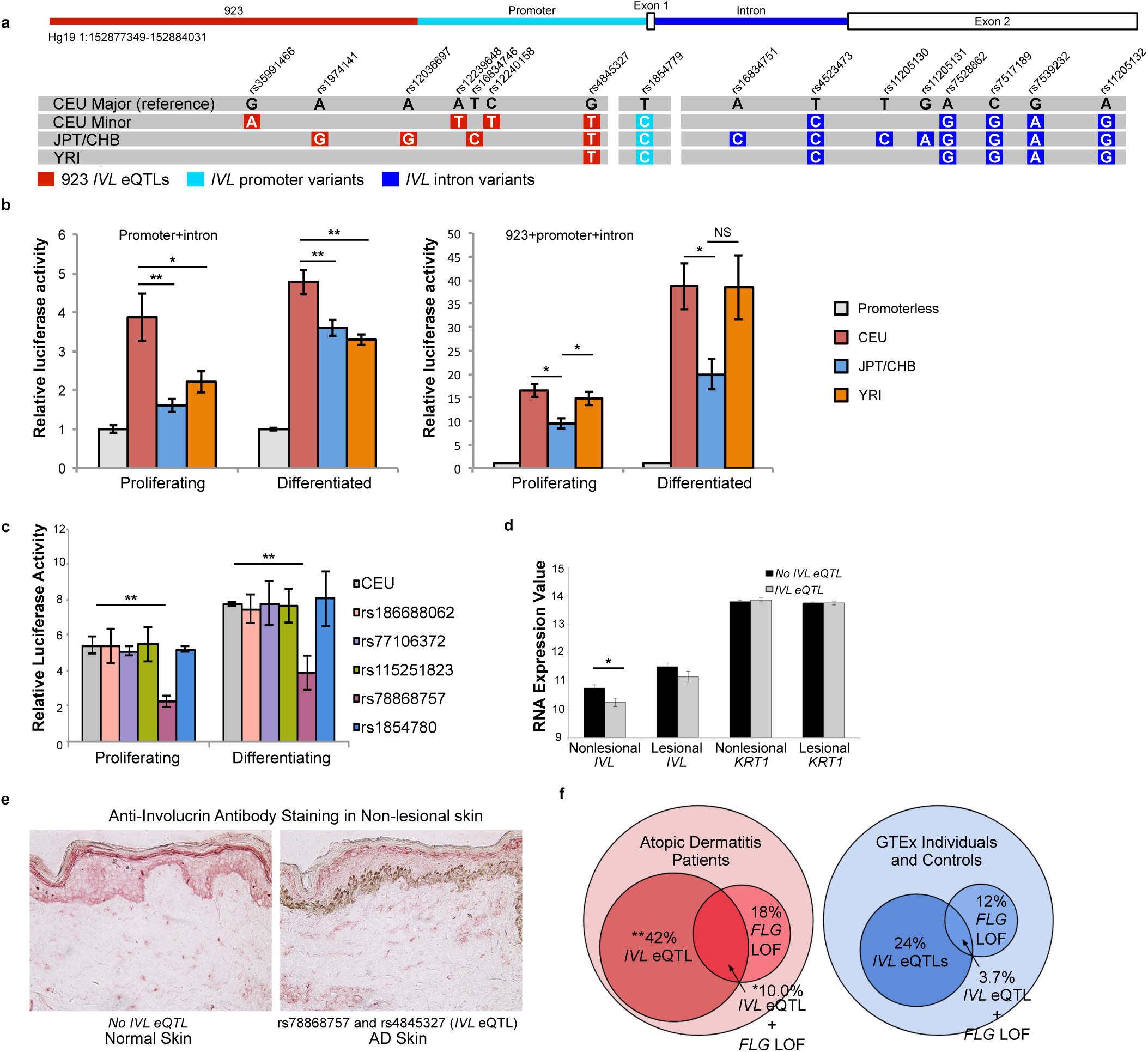
Population-specific 923 enhancer:*IVL*promoter/intron alleles and eQTLs impact regulatory activity and gene expression and are enriched in AD subjects. **a**) Schematic of population-specific human 923 enhancer (red), *IVL* promoter (light blue), 1^st^ exon, and intron (dark blue) alleles with phased common haplotypes for the 7 *IVL* eQTLs in 923. Positions of variants not to scale. **b)** Luciferase assays of population-specific alleles for the *IVL*promoter/intron + respective enhancer reveals that the CEU promoter/intron allele has the highest regulatory activity. The addition of the enhancer confers higher reporter expression, especially in differentiated cells, with more activity in CEU and YRI than JPT/CHB. Mean ± SEM of 3-4 independent experiments. *, *P*<0.05; **, *P*<0.01, ANOVA followed by Tukey’s HSD. **c**) Luciferase assays of 923 singleton alleles in SP-1 keratinocytes, Mean ± SEM, n = 3, \**P*<0.05 vs. 923 reference. **d**) Decreased *IVL*, but not *KRT1*, mean gene expressions in the lesional and nonlesional skins of *IVL* eQTL (n=13) vs. *IVL* non-eQTL AD (n=23) patients, microarray, Mean + SEM, \**P*<0.05, Student’s t-test, two-tailed. **e**) Decreased IVL protein expression in the AD lesional skins of *IVL* eQTL and 923 enhancer rs78868757 in AD patient compared to non-*IVL* eQTL genotyped normal skin. **f**) Venn diagrams highlighting frequencies of *IVL* eQTLs and *FLG* LOF variants and was enriched in atopic dermatitis subjects vs. controls, \**P*<0.05.

### *IVL* eQTLs in the 923 enhancer are a modifier for *FLG* LOF-associated AD

The impact of the 923 enhancer variation on *IVL* expression in human evolution led us to examine a potential role for the 923 enhancer variation in skin disease. Atopic dermatitis (AD) is a common inflammatory skin disease with skin barrier deficiency as a major etiological factor, as evidenced by the discoveries for more than 100 pathogenic loss-of-function (LOF) variants in the filaggrin (*FLG*) gene^30–32^. The observation that not all *FLG* LOF heterozygotes are diagnosed with AD supports a hypothesis of incomplete penetrance and suggests additional modifiers for the disease. We hypothesized that 923 enhancer variation, and resulting decreased *IVL* expression, impact *FLG* LOF-associated AD. Targeted sequencing of *FLG* and the 923 enhancer were independently performed in subjects with moderate to severe AD (n=110) and healthy controls (n=30)^33–37^. Individuals from GTEx with whole genome sequencing data (n=642)^25^ were included as additional controls and variants were identified through filtering of vcf files.

Targeted sequencing of the 923 enhancer region in AD cases identified 30 variants (Table S13). Three were unique to cases (rs1344252044, rs139775257, and rs76513010) and strikingly, seven of the 30 variants are *IVL* eQTLs associated with decreased *IVL* expression in skin and correspond with two different haplotypes, CEU minor and JPT/CHB (Fig. 5a-b, Table S14). In fact, a ranked list of effect sizes for genes with eQTLs within the EDC revealed *IVL* as the gene exhibiting the largest range of effect sizes for decreased expression (Table S15). *IVL* was the only gene to harbor eQTLs that all had negative effect sizes which correlates with the CEU haplotype being associated with increased *IVL* expression (171 eQTLs; -0.16 to -1.00) (adjusted *P* < 0.05) (GTEx V7, Tables S14). Through sample allele phasing and imputation, we established that 77% of the 30 alleles present in the AD cases harbored at least one *IVL* eQTL (n=22) (Fig. S6). Luciferase assays for single variant alleles revealed no differences in four discovered variants (rs186688062-A, rs77106372-A, rs115251823-T, and rs1854780-A minor alleles); however, the rs78868757-A variant exhibited significantly decreased enhancer activity compared to the major allele in both proliferating and differentiated keratinocytes (∼2-fold; proliferating [*P* < 0.01]; differentiated [*P* < 6.4 × 10^−5^]) (Fig. 5c). We next examined the impact of the *IVL* eQTLs in AD subjects in both nonlesional and lesional skin areas. AD cases with *IVL* eQTLs exhibited a trend toward decreased *IVL* expression in lesional skin and significantly decreased expression in nonlesional skin (*P* < 0.14 and 0.04, respectively) (Fig. 5d). As a result, IVL immunohistochemical staining was markedly decreased in the patients’ nonlesional skin in comparison to non-*IVL* eQTL skin (Fig. 5e). Our results for *IVL* eQTLs and the uncommon rs78868757 923 enhancer variant associated with decreased *IVL* gene and protein expression highlight the functional impact of these enhancer variants on diseased skin.

### Significant enrichment of *IVL* eQTLs and *FLG* LOF variants in atopic dermatitis patients

We evaluated the impact of having at least one *IVL* eQTL and at least one *FLG* LOF variant on AD outcome. *FLG* LOF variants were enriched in AD cases compared to controls, but did not reach significance (*P* < 0.06). The proportion of AD patients with at least one *IVL* eQTL was 42% (n=47) and significantly enriched compared to healthy and GTEx controls (24%; *P*< 0.0001) (Fig. 5f). Among the controls, only 3.7% (n=25) genotyped for both an *IVL* eQTL and a *FLG* LOF variant (Fig. 5f). By contrast, this genetic combination was significantly enriched in AD patients at 10.0% (n=11; *P* < 0.0109). An analysis of the effect of this genetic combination on disease severity was inconclusive given the small number of patients with SCORAD data in our cohort (data not shown). Nevertheless, our data identifies a significant enrichment of *IVL* eQTLs in the presence of *FLG* LOF variants, thus identifying *IVL* eQTLs and *FLG* LOF variants as potential modifiers for the skin barrier in AD pathogenesis.

## DISCUSSION

Our work thus far has identified recent skin barrier evolution in modern human populations, having discovered multiple population-specific signals of positive selection within the EDC and selection in two independent yet convergent haplotypes for *IVL*. The associations of CEU- and JPT/CHB-specific haplotypes with relatively increased and decreased *IVL* gene expression, respectively, and their overlap to the known epidermal-specific enhancer 923 led us to examine a functional role for the 923 enhancer to regulate *Ivl* expression *in vivo.* Deletion of the orthologous mouse 923 enhancer via CRISPR/Cas9 editing identified *Ivl* as the primary target gene of 923 with regulation in *cis*, thus establishing a 923 enhancer-*Ivl* gene regulatory module. The functional significance of this genetic module in humans was further revealed with our discoveries of population-specific 923 enhancers that “boosted” regulatory activity and with the observation that *IVL* eQTLs are enriched in atopic dermatitis patients with *FLG*LOF variants. Together, we establish the significance of a paradigmatic regulatory module for a cis-regulatory enhancer/target gene in both human evolution and disease.

Our understanding of the molecular mechanisms that drive enhancer activity^38,39^ and the comprehensive genomic elucidation of putative enhancers for a vast array of tissue types, as well as single cells, has exponentially grown since the inception of the “enhancer” concept in 1981^40^. Yet we are challenged to embark on more rigorous investigations to functionally validate enhancers and correctly assign their predicted target genes^39^. Even more challenging is the growing body of recent reports demonstrating a lack of phenotypes for enhancer deletions, even for putative developmental enhancers, suggesting enhancer redundancy consistent with the ENCODE findings of 2-3 enhancers per target gene^41–43^. As *Ivl* expression was not completely lost in our enhancer knockout mice, there is likely redundancy in regulation of *Ivl* yet we have firmly established the 923 enhancer as a prominent proximal enhancer. 5C studies in keratinocytes identified physical interactions between the *Ivl* promoter and enhancers located in the first topologically associated domain (TAD) of the EDC, presenting additional putative regulators^44^. Our discovery for the 923 enhancer regulation of *Ivl* expression was premised on extensive previous analyses^7,26^ and offers a framework to prioritize such ongoing and future enhancer *in vivo* deletion studies. Furthermore, our CRISPR/Cas9 methodology led to the serendipitous generation of a larger deletion (923^large^), which displayed decreased *Lce6a* expression that generates the hypothesis for additional enhancers that were deleted in the 923^large^ allele.

Another interesting finding is the effect on DARs due to loss of 923 enhancer. DARs that were enriched within the EDC were less accessible upon deletion of 923, minimally suggesting a direct role for the enhancer in governing physical chromatin accessibility within the EDC. The shared EDC DARs are also located in different TADs than 923, supporting inter-TAD interactions as previously described^44^. The mechanism of chromatin accessibility regulation by 923 remains elusive, however, no physical interactions were identified between 923 and these DARs in previous 3C and 5C analysis of the EDC^26,44^. Also of note, one of the shared DARs almost completely overlaps the pseudogene *Gm29950* that was not detected in our whole skin RNA-seq data set for 923^del/del^, 923^large/large^, *Ivl*^-/-^ and WT keratinocytes but warrants further higher resolution investigation in this region. It is also interesting that we did not identify the 923 enhancer as an accessible region in wild-type cells. We hypothesize the dense occupation of transcription factors at this active enhancer biases against transposase insertion needed for ATAC-seq, resulting in the interpretation of ‘closed’ chromatin at the focused region yet enrichment at the flanking sites, a hypothesis that should be considered when using chromatin accessibility as a genome-wide predictor of enhancers. Overall, our ATAC-seq illustrates a role for the 923 enhancer in governing local chromatin accessibility.

Interestingly, the backbone of the CEU-*IVL* haplotype appears to have emerged in Africa and rose to high frequency in CEU. This recent selective sweep for CEU-*IVL* is associated with increased *IVL* expression, which we tied to the promoter/intron variants. The promoter SNP rs1854779 is a strong candidate for the causal variant, as it is predicted to generate a gain-of-function NFI transcription factor binding site that could increase transcription. By contrast, the positively selected JPT/CHB enhancer/promoter/intron allele exhibited relatively decreased reporter activity and contains unique polymorphisms in the enhancer compared to the other population-specific alleles, suggesting enhancer variation as causative for decreased expression. We hypothesize that the selective forces present in East Asia favor reduced *IVL* expression; however, the precise nature of these forces is unknown. The *IVL* eQTLs that define the JPT/CHB haplotype and the other 923 *IVL* eQTLs fall within the boundaries of the three largest peaks for H3K27ac marks, suggesting that the actual enhancer boundary may not extend to the smaller, most 5’ H3K27ac peak (Fig 3a). Our data linking *IVL* eQTLs with AD further supports the emerging paradigm of previously-selected alleles that are now at a “mismatch” in the context of our current lifestyles and environment in the industrialized world^45^. These findings motivate future studies to more closely examine *IVL* eQTLs (and other eQTLs in the EDC) as potential modifiers of penetrance and severity for *FLG*-LOF AD (Table S14).

Our discovery of enhancer variation for human skin barrier function further contributes to a growing body of research that highlights selective events in human history targeted at enhancers, such as lactase persistence^46–50^ and immune function^51^. Furthermore, our finding for enhancer evolution further supports genetic innovation for human skin evolution that until now has been reported for only a few genes in the EDC, including *IVL*^52^. *IVL* has undergone extensive evolution in primates, with recent and continuing expansion of the number of repeats in human *IVL*^53–61^. However, targeted deletion of the *Ivl* gene in mice revealed no overt phenotype in barrier-housed conditions^62^, similar to our 923^del/del^ and 923^large/large^ homozygous mice (Fig. S1). Together this supports a tolerance for decreased, and even lack of, Ivl in barrier-housed conditions, consistent with human population-specific selection for differential *IVL* expression that warrants further investigation. The findings for a recent selective sweep for increased IVL in Northern latitudes suggests a possible link between cutaneous vitamin D production and *IVL* expression as a mechanism of environmental adaptation for the skin barrier. Vitamin D is known to stimulate the differentiation of keratinocytes and promote the expression of CE genes, including *IVL*; it is possible that high rates of vitamin D deficiency in modern populations^63^ could contribute to reduced skin barrier integrity and, subsequently, disease states.

In summary, our results highlight the significance of genetic variation attributed to differential gene expression in a population-specific manner and heighten our awareness for human population-specific evolution of the epidermis. Furthermore, our work provides a framework with which we can examine genetic variation in enhancer:target gene modules for recent adaptive traits. It also suggests a need for future studies to more closely examine variants associated with recently evolved traits as newly emerging disease risk variants during human transition to modernity.

## Acknowledgements

We thank Renate Lewis at the Hope Center for the design and targeting of the gRNAs, Mia Wallace at the Mouse Core facility for the mice, Toni Sinnwell at the Washington University Genome Technology Access Center for the preparations of the RNA-seq libraries and Illumina sequencing, and Justin Fay, Don Conrad, Gary Stormo, and John Edwards for helpful discussions. The work cited in this publication was performed in a facility partially supported by NCI Cancer Center Support Grant P30CA91842 to the Siteman Cancer Center and by ICTS/CTSA UL1TR002345 and C06RR015502 from National Center for Research Resources (NCRR), a component of the National Institutes of Health (NIH), NIH Roadmap for Medical Research, Society of Investigative Dermatology/Sun Pharma Research Innovation Fellowship (EAB), NHGRI T32HG000045 (ZAG), T32GM007067 (MEM), R25GM103757 (AS), A*STAR-BMRC-EDBH17/01/a0/004 (XFCCW, SLIJD and JEAC) and by NIAMS R01AR065523 (CdGS). This content is solely the responsibility of the authors and does not necessarily represent the official views of NCRR or NIH.

