## Supplemental Materials and Methods for "An enhancer:involucrin regulatory module impacts human skin barrier adaptation out-of-Africa and modifies atopic dermatitis risk"

**CMS and iSAFE Analyses**

CMS scores for all SNPs in the EDC (hg18; chr1:150,198,268-151,892,013) in CEU, YRI and JPT/CHB populations were downloaded (https://www.broadinstitute.org/cms/results; download date August 17, 2015). iSAFE (<https://github.com/alek0991/iSAFE>) was used to identify positive selection in 1KGP Phase 3 phased data (<http://ftp.1000genomes.ebi.ac.uk/vol1/ftp/release/20130502>) (hg19; chr1:151,896,000-153,612,000). Sample cases were defined as all subjects belonging to the population of interest (CEU, YRI, or JPT and CHB) and sample controls defined as subjects belonging to all other superpopulations. All other arguments were run using default settings.

**Haplotype Block Construction and LD analysis**

Haplotypes were identified using LDLink (<https://ldlink.nci.nih.gov>) by querying the LDproxy module for the SNP with the highest CMS score in the relevant population(s). Haplotypes were defined as the set of proxy variants with R^2^≥0.8. Pairwise and small-scale analyses of LD were performed using the LDpair module, again defining LD as R^2^≥0.8.

**Global Allele Frequency and eQTL Analysis**

Allele frequencies of EDC SNPs from the twenty-six 1KGP populations were obtained from Ensembl v. 81^1,2^. Geographical latitude for each population was determined by the latitude of the city from which the 1KGP population data was collected. The relationship between latitudes and allele frequencies was analyzed using linear regression using the lm function in R. The direction and strength of the correlation between allele frequency and latitude was determined using Spearman’s correlation coefficient (*ρ*) using the cor function in R. Geographic plots for the global distribution of allele frequency was generated using ggplot2 in R. SNPs were queried as eQTLs in sun-exposed or not sun-exposed skins using GTEx portal ([http://www.gtexportal.org](http://www.gtexportal.org/); V7 release).

**Generation of 923 enhancer knockout alleles in mice by CRISPR/Cas9 genome editing**

Deletion of the 923 enhancer was targeted using 2 small guide RNAs to facilitate Cas9-mediated double stranded breaks on either side of the orthologous mouse 923 enhancer sequence. *LoxP* insertions at these flanking sgRNA targeted sites were generated using two single stranded oligodeoxynucleotides ssODNs (containing *loxP* sites with specific 80bp homology arms on either side of *loxP* and restriction enzyme sites (SphI for 5’ end, HindIII at 3’ end) which were simultaneously introduced via by site-directed homologous recombination (Integrated DNA Technologies, Coralville, IA). Sequences of sgRNAs and ssODNs are listed in Supplementary Material (Table S4). Three rounds of injection in 779 zygotes were performed having confirmed target specificity using *in vitro* pilot studies prior to zygote injection. Founders were initially screened for large deletions via PCR using flanking primers designed outside the homology arms and 5’ and 3’ *loxP*-specific primers that were resolved on 1% agarose gel electrophoresis. Of the 779 C57BL6/6XCBA hybrid zygotes injected, 80 F_0_newborns were recovered, of which 75 survived to weaning age. Seven out of 80 mice whose 923 allele size deviated from the wild-type allele were identified via PCR (8.75% targeting efficiency) with further analysis by Sanger sequencing. Two 923 enhancer knockout alleles were confirmed. 923^del^ allele contains a ~1250bp deletion of the 923 enhancer and the 5’ *loxP* site. The 923^large^ allele includes the 923 deletion flanked by both 5’ and 3’ *loxP* sites and a 40kb deletion between proximal gene *Lce6a* and 923, including the genes *Smcp* and *2210017I01Rik*. Genotyping primers are listed in Supplementary Material (Table S4). All mice were housed in pathogen-free, barrier facilities at Washington University School of Medicine (St. Louis, MO). All animal procedures were approved by the Division of Comparative Medicine Animal Studies Committee at Washington University. All animal work was conducted in accordance with the Guide for the Care and Use of Laboratory Animals of the National Institutes of Health. Morning observation of a vaginal plug was designated as embryonic day (E) 0.5. Both 923^del^ and 923^large^ mouse lines were backcrossed at least 7 times to the C57BL/6 background to generate isogenic strains and to exclude potential off-target effects arising from CRISPR/Cas9 editing.

**RNA-seq**

Total RNA from whole skin was isolated by TriZol extraction (Life Technologies, Frederick, MD). Ribo-zero (ribosome-depleted) RNA sequencing libraries were prepped according to the manufacturer’s library kit protocol, indexed, pooled, and sequenced on Illumina HiSeq 3000 (1 X 50bp) by the Washington University Genome Technology Access Center.

Basecalls and demultiplexing were performed with Illumina’s bcl2fastq software and a custom python demultiplexing program with a maximum of one mismatch in the indexing read. RNA-seq reads were then aligned to the Ensembl release 96 top-level assembly with STAR version 2.0.4b. Gene counts were derived from the number of uniquely aligned unambiguous reads by Subread:featureCount version 1.4.5. Sequencing performance was assessed for the total number of aligned reads, total number of uniquely aligned reads, and features detected. The ribosomal fraction, known junction saturation, and read distribution over known gene models were quantified with RSeQC version 2.3. All gene counts were then imported into the R/Bioconductor package EdgeR and TMM normalization size factors were calculated to adjust for samples for differences in library size. Ribosomal genes and genes not expressed in the smallest group size minus one samples greater than one count-per-million were excluded from further analysis. The TMM size factors and the matrix of counts were then imported into the R/Bioconductor package Limma. Performance of the samples was assessed with Spearman correlations, and a Multi-Dimensional Scaling plot, and hierarchical clustering. Weighted likelihoods based on the observed mean-variance relationship of every gene and sample were then calculated for all samples with the voomWithQualityWeights. The performance of all genes was assessed with plots of the residual standard deviation of every gene to their average log-count with a robustly fitted trend line of the residuals. Differential expression analysis was then performed to analyze for differences between conditions and the results were filtered for only those genes with Benjamini-Hochberg false-discovery rate adjusted p-values less than or equal to 0.05, and a log2(fold change) ≥ |2|.

The R/Bioconductor package heatmap3 and Pathview was used to display heatmaps or annotated KEGG graphs across groups of samples for each GO term or KEGG pathway (respectively) with a Benjamini-Hochberg false-discovery rate adjusted p-value less than or equal to 0.05.

Real-time qPCR on cDNA (generated using SuperScript II reverse transcriptase (Thermo Fisher Scientific) using TaqMan Gene Expression Assay was performed in triplicate (Applied Biosystems, Life Technologies, Carlsbad, CA) and normalized to β2-microglobulin. Only CT values with single peaks on melt-curve analyses were included.

**Allele-specific gene expression**

RNA from newborn whole skins was isolated as described above from C57Bl6 and BALB/cBYJ wild type mice and from [C57Bl6]/[BALB/cBYJ], [923^del^/[BALB/cBYJ], and [923^large^/[BALB/cBYJ] heterozygous mice. RNA was DNaseI treated and reverse transcribed into cDNA using Invitrogen SuperscriptII Reverse Transcriptase (Invitrogen, Carlsbad, CA). 260bp amplicons targeting the *Ivl, 2210017l01Rik*, and *Lce6a* genes were amplified from cDNA using NEBPhusion High Fidelity PCR 2X master mix (New England BioLabs, Ipswich, MA) (Amplicon Sequence and primers in supplement). PCR products were purified on Qiagen QIAquick PCR columns (Hilden, Germany), A-tailed with 1mM dATP and NEB Taq polymerase for 20min at 72°C (New England BioLabs, Ipswich, MA) followed by an additional column purification on Qiagen MinElute columns. Next-generation sequencing compatible adapters were ligated to A-tailed PCR products in molar excess using the LigaFast DNA ligase kit (Promega, Madison, WI) at room temperature for 20 minutes. Products were size selected using Agencourt AMPure XP beads (Beckman Coulter, Brea, CA) at 1.2X product volume to remove excess adapter. Adapter ligated products were quantitated using Qubit dsDNA High Sensitivity Assay kit (Life Technologies, Carlsbad, CA). To determine appropriate PCR amplification cycle number to avoid over amplification, quantitative PCR was preformed using 2X NEB Phusion High Fidelity PCR master mix, 0.5uM PCR primer1, 0.01uM PCR primer2, 0.5uM unique Index Primer(sequences in supplement), 100X SYBR Green and 50X ROX Dye and 2ng DNA in 10ul. PCR cycle number for each template was determined by identifying the cycle where ¼ max fluorescence was reached. 10ng of each library was amplified using the same reaction, without SYBR and ROX, scaled up to 50ul. Amplified libraries were size-selected again to remove any remaining adapter dimer using 1X concentration of AmPure beads. All libraries were pooled at equal molar ratio and sequenced as a spike-in to a 2x150bp MiSeq sequencing run, averaging 154,000 reads per sample. De-multiplexed reads were mapped using Bowtie2 and visualized using the IGV viewer. The proportion of nucleotides at each informative snp in the amplicon was calculated by IGV. Primers are listed in supplementary material (Table S16).

**ATAC-Seq**

ATAC-seq was performed on 75,000 epidermal cells from each mouse as previously described with variations^3^. Cells were lysed for 5 mins in 37.5ul ice cold buffer (10mM TrisHCl, 10mM NaCl, 2mM MgCl, 0.5% IGEPAL CA-630). Cells were then pelleted at 500g (4°C) for 15mins. Lysis buffer was replaced with transposition reaction mix (12.5ul TD (2x reaction buffer from Nextera Kit, Illumina, San Diego California, USA), 2.5ul TDE1 (Nextera Tn5 Transposase from Nextera kit, Illumina), and 10ul H_2_O) and samples were incubated for 1 hr at 37°C. Samples were purified using Qiagen MinElute PCR Purification Kit (Qiagen, Valencia, CA) and PCR-amplified. Adapter dimer bands were removed using AMPure XP bead treatment as previously described (Beckman Coulter, Brea, CA). All Samples exhibited expected nucleosome periodicity as assayed by High Sensitivity ScreenTape (Agilent Technologies, Santa Clara, CA). Samples were sequenced via Illumina HiSeq2500 (2 X 50bp). An average of 93.4% of the reads were mapped with 14.3-39.8 million qualified reads per sample with only an average of 6.8% mitochondrial reads (Table S17). Prior to sequencing, all samples exhibited the expected periodicity of insert length and were enriched for reads at transcription start sites. ATAC-seq data was processed using the ENCODE ATAC-seq processing pipeline using Caper with Conda (<https://github.com/ENCODE-DCC/atac-seq-pipeline>). Reads were mapped using Bowtie2, and filtered to remove unmapped reads, duplicates, and reads mapping to chrM. Peaks were called on each replicate using MACS2^4^. Biological replicates were included if both the rescue and self-consistency IDR values per genotype were below (or very near) 2 (Table S17). Differential accessibility was assessed using EdgeR^5^ within the DiffBind R package ([http://bioconductor.org/packages/DiffBind/](https://bioconductor.org/packages/DiffBind/))(FDR <0.5, log2(FC) ≥|2|).

**Variant Analyses of GTEx WGS samples**

GTEx vcf files for whole genome sequencing were downloaded resulting in genotypes for 642 unique individuals. A list of individuals with *FLG* LOF variants was obtained by filtering for 1) variants in *FLG* and subsequent annotation for LOF with SnpEff (v4.3)^6^ and 2) known *FLG* LOF from ExAC^7^, gnomAD (V2.1.1)^8^ and in-house lab lists^9^. A list of individuals with *IVL* eQTLs was obtained by filtering for the known *IVL* eQTLs in the 923 enhancer (rs35991466, rs1974141, rs12036697, rs12239648, rs16834746, rs12240158, rs4845327).

**Luciferase Assays**

Population-specific alleles for the *IVL* promoter, noncoding first exon, and intron were cloned from human gDNA by PCR using primers 5’-GGATCCGATAGGTTCTAGGGGTATAGTGG/5’-AAGCTTCTTAGAAGCTACTGTCAACCTG (restriction sites underlined). PCR products were digested with BamHI and HindIII and cloned into the BglII/HindIII site in pGL3 (Promega) to yield pGL3-IVLpromoter and confirmed by Sanger sequencing. The Gateway cassette B was then cloned into the SmaI site to yield pGL3-IVLpromoter-GW. The 923 enhancer region was amplified from human gDNA using primers 5’- GGGG**ACCACTTTGTACAAGAAAGCTGGGT**GAAGAACAGTGAATTTTACGACC/5’- GGGG**ACAAGTTTGTACAAAAAAGCAGGCT**AGACATTCTGCTGCTGGACA (attB sites in bold) and introduced into pDONR221 by BP recombination using BP Clonase II (Invitrogen, Thermo Fisher Scientific). Haplotype-specific variants were confirmed by Sanger sequencing. Enhancer alleles were then introduced into pGL3-*IVL*promoter-GW by LR recombination using LR Clonase II (Invitrogen, Thermo Fisher Scientific) to yield pGL3-*IVL*promoter-923. For testing singleton variants rs186688062, rs77106372 and rs1854780 within 923 from AD patients, the enhancer was amplified from patient gDNA and introduced into pDONR221 as above. Enhancer alleles were then introduced to pGL3i1Rfb12^10^ by LR recombination. Luciferase reporter assays were performed as previously described^10,11^, with measurements performed at 48 hr and 72 hr post-differentiation for proliferating and differentiated cells, respectively using a Glomax luminometer (Promega). Significance was determined using ANOVA followed by Tukey’s HSD. For testing singleton variants rs115251823 and rs78868757 within 923 from AD patients, synthetic gBlocks (IDT) of the 923 enhancer (Hg19 1:152878418-152879076) containing each variant were cloned into a pGL3 firefly luciferase vector and confirmed by Sanger sequencing. SP-1 mouse keratinocytes were transfected for each of the pGL3 variants and Renilla luciferase vectors using Lipofectamine 2000. Dual-luciferase assays were performed on the transfected proliferating and differentiated cells as previously described^10,11^. Significance was determined via two-tailed Student’s t-test.

**DNA Isolation**

Human DNA was isolated from blood samples using PAXgene Blood DNA kit. The quality of DNA was confirmed by visualization agarose gel electrophoresis and spectrophotometry to ascertain the high molecular weight of DNA and 260/280 nm values, respectively. DNA quantification was accomplished by PicoGreen fluorescence prior to sequencing.

**Targeted Amplicon Sequencing**

Individual DNA (375ng per individual sample) were microfluidically arrayed (Fluidigm), indexed, and amplified using 23 primer sets that tile across the 923 enhancer (Methods Table S5) (Genome Technology Access Center [GTAC], Department of Genetics, Washington University School of Medicine). Paired end sequencing (Illlumina MiSeq, 2 X 150) was performed and raw reads were subsequently demultiplexed and aligned (hg19 human reference). Reported variants were sequenced with a minimum of 100X and 200X coverages for *FLG* and the 923 enhancer, respectively. Targeted sequencing for *FLG* was performed as previously described^12^. Variants called were subsequently validated by either Sanger sequencing or TaqMan allele-specific probes.

**Immunohistochemistry**

Filaggrin protein was detected using mouse anti-filaggrin monoclonal antibody (Abnova, MAB3538). Involucrin protein was detected using mouse anti-involucrin monoclonal antibody (Sigma, sc-21748).

**AD patient gene expression**

The expression of *FLG* and *IVL* were determined by microarray as previously described^13,14^.

**Histology**

Dorsal skin was excised from 8-week old mice and preserved in 4% paraformaldehyde (Electron Microscopy Sciences, Hatfield, PA) prior to paraffin sectioning. Sections were stained with hematoxylin and eosin by the Washington University Developmental Biology Histology Core. Slides were imaged on a Nikon Eclipse 80i brightfield microscope (Nikon, Tokyo, Japan).

**Dye penetration barrier assays**

Barrier assays were performed as previously described^15^ with X-gal solution incubations for at least 4 hours at 37°C. Images were captured on a CanoScan 5600F scanner (Canon, Melville, NY).

**Cornified envelope preparations**

Epidermis (cut into 1cm^2^ pieces) was incubated at 95°C in a solution of 2% SDS to obtain cornified envelopes in a single-cell suspension as previously described. The suspension was placed on a slide and inspected using phase contrast light microscopy.

**Trans-epidermal water loss assay**

TEWL was measured on the abdominal skin surface of newborn mice using the nail attachment of a VapoMeter (Delfin Technologies, Kuopio, Finland, courtesy of Jeff Miner).

**Immunohistofluorescence**

Fresh sections were fixed in 4% paraformaldehyde (Electron Microscopy Sciences, Hatfield, PA) prior to permeabilization with 0.1% Triton X-100 and subsequent antibody incubation. The following antibodies were used for immunofluorescence: rabbit IVL (4b-KSCN, 1:200), chicken K14 (5560, 1:500; courtesy of J Segre), goat anti-rabbit (Alexa Fluor 488, 1:500), and goat anti-chicken (Alexa Fluor 594, 1:500) IgG antibodies (Life Technologies, Frederick, MD) and DAPI counterstained (SlowFade Gold antifade reagent) (Life Technologies). Fluorescent microscopy was performed on Zeiss AxioImager Z1 and captured with AxioCam MRc and Axiovision software (Carl Zeiss, Stockholm, Sweden), or imaged on a DMI3000 B (Leica, Wetzlar, Germany) and captured with the QIClick camera (QImaging, Surrey, BC, Canada).
