## Supplemental Figure Legends for "An enhancer:involucrin regulatory module impacts human skin barrier adaptation out-of-Africa and modifies atopic dermatitis risk"

**Supplemental Table Legends**

**Table S1. SNPs in clusters with high CMS scores (CMS>0) for each population.** The genomic region defined by nearby genes is also indicated.

**Table S2. iSAFE scores for EDC SNPs in JPT/CHB.** iSAFE scores are reported and the SNPs located in iSAFE peaks are marked by bold and highlighting.

**Table S3. iSAFE scores for EDC SNPs in CEU.** iSAFE scores are reported and the SNPs located in iSAFE peaks are marked by bold and highlighting.

**Table S4. CRISPR/Cas9 editing strategy reagents and mouse allele sequences.** a) Sequences of small guide RNAs used to target the Cas9 protein on a C57BL/6 genetic background. b) Single-stranded oligonucleotide (ssODN) sequences utilized with loxP site and introduced informative restriction sites marked. c) Sanger sequencing of wildtype 923, d) 923^del^ allele, e) 923^large^ allele, and f) primers used to genotype each allele.

**Table S5. Offspring genotype distribution from heterozygous 923^del^ as well as 923^large^ parental crosses.** Chi-squared test calculations included.

**Table S6. Ranked list of differentially expressed genes between 923^del/del^ and WT mice whole skin from RNA-seq.** List ranked by log2FC. FDR (adj.P.val) <0.05 and logFC ≤|2| cutoffs used.

**Table S7. Ranked list of differentially expressed genes between 923^del/+^ and WT mice whole skin from RNA-seq.** List ranked by log2FC. FDR (adj.P.val) <0.05 and logFC ≤|2| cutoffs used.

**Table S8. Ranked list of differentially expressed genes between 923^large/large^ and WT mice whole skin from RNA-seq.** List ranked by log2FC. FDR (adj.P.val) <0.05 and logFC ≤|2| cutoffs used.

**Table S9. Ranked list of differentially expressed genes between 923^large/+^ and WT mice whole skin from RNA-seq.** List ranked by log2FC. FDR (adj.P.val) <0.05 and logFC ≤|2| cutoffs used.

**Table S10. Ranked list of differentially expressed genes between *Ivl*^-/-^ and WT mice whole skin from RNA-seq.** List ranked by log2FC. FDR (adj.P.val) <0.05 and logFC ≤|2| cutoffs used.

**Table S11. Ranked list of differentially accessible regions between 923^del/del^ and WT mice epidermis from ATAC-seq.** List ranked by FC. FDR<0.05 and FC ≤|2| cutoffs used.

**Table S12. Ranked list of differentially accessible regions between 923^large/large^ and WT mice epidermis from ATAC-seq.** List ranked by FC. FDR <0.05 and FC ≤|2| cutoffs used.

**Table S13. Variants in the 923 enhancer region found in Atopic Dermatitis Patients.**

**Table S14. Ranked list of genes in the EDC with eQTLs and their gene expression effect size range.** List ranked by location.

**Table S15. Ranked list of *IVL* eQTLs.** List ranked by location.

**Table S16. Allele-Specific Amplicons.** Amplicon primers length in base pairs. Amplicon sequences with SNPs bolded and primers underlined.

**Table S17. ATAC-Seq Library Statistics.** Included statistics reported as recommended by ENCODE guidelines.

**Supplemental Figure Legends**

**Figure S1. Both 923 deletion mouse lines exhibit normal morphology under barrier-housed homeostatic conditions. a**) Homozygous 923^del/del^ and 923^large/large^ mice are viable and appear normal in barrier-housed conditions at newborn or 8 weeks when compared to aged matched control WT (+/+) mice. **b**) H&E staining of WT, 923^del/del^, and 923^large/large^ epidermal sections from newborn and 8-week-old (adult) mice appear normal. **c**) Normal cornified envelope morphology observed in homozygous 923^del/del^ and 923^large/large^ keratinocytes compared to WT littermates**.** Similar quantities of angular and balloon shaped cornified envelopes with smooth edges were isolated from newborn skin of homozygous deletion and wildtype littermates of 923^del^ and 923^large^. **d**) Normal inside-out skin barrier function in 923 deletion mice. Barrier function was measured by transepidermal water loss (TEWL). e) Normal patterning of skin barrier development in 923 deletion mice. The extent of skin barrier formation was assessed by an outside-in X-gal dye penetration assay. Blue stain indicates X-gal reactivity with endogenous β-galactosidase where the X-gal solution has penetrated the epidermis where the skin barrier has not formed.

**Figure S2. Heatmap of 5 differentially expressed genes in 923^del/del^, 923^large/large^ and *Ivl*^-/-^ mice skin RNA-seq compared to WT.**

**Figure S3. Chromatin accessibility in and around the EDC in 923^del/del^ and 923^large/large^ newborn epidermis with all replicates displayed.** ATAC-seq signal tracks for each biological replicate shown as fold change signal per genotype (WT, *n*=3; 923^del/del^, *n*=4; 923^large/large^, *n*=3) with shared less accessible (red) and more accessible (blue) DARs indicated in the 923 deletion lines compared to WT.

**Figure S4. Differentially Accessible Regions in 923^large/large^ epidermis corresponds with Differentially Expressed genes in the EDC. a**) Venn diagram of the overlap between DAR-associated genes and DE genes. **b**) ATAC-seq signal track of each genotype for the *Ivl*-*Lce6a* region showing the 923 deletion alleles. Displayed DARs are unique to 923^large/large^ allele.

**Figure S5. Predicted NF1 binding sites are gained in the CEU allele of rs1854779.** Logos for NFIX and NFIC, two isoforms of NF1, are shown above the nucleotide sequence surrounding rs1854779 in the reference (CEU) and alternate (JPT/CHB and YRI) alleles. The nucleotide at rs1854779 is boxed.

**Figure S6. Phased and imputed 923 enhancer alleles in AD patients.** Schematic of 923 enhancer alleles marked by variants (red) found in AD patients (n=110). Phased alleles were validated through sanger sequencing. Positions of variants not to scale.
