## SupplementalFigures for "An enhancer:involucrin regulatory module impacts human skin barrier adaptation out-of-Africa and modifies atopic dermatitis risk"

### Supplementary Figure 1

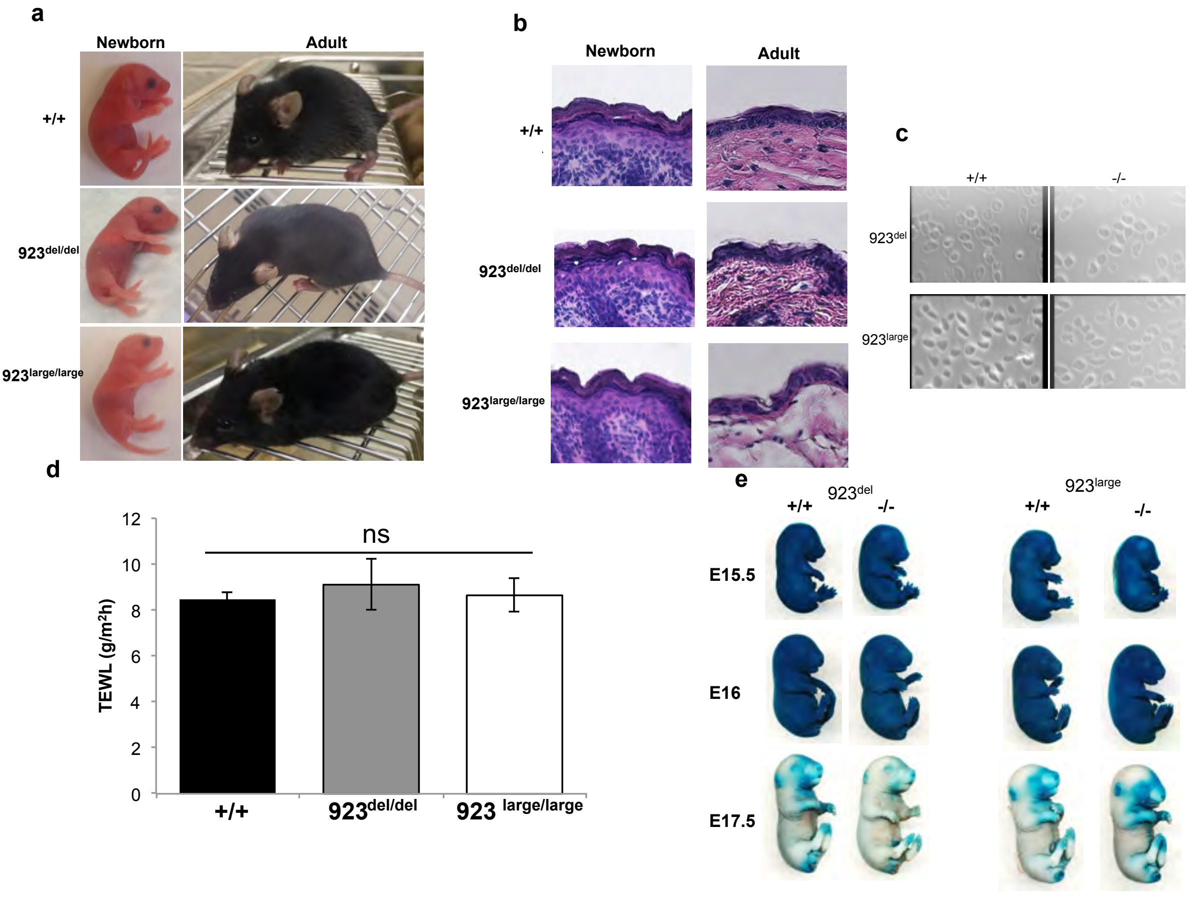

**Supplementary Figure 2**

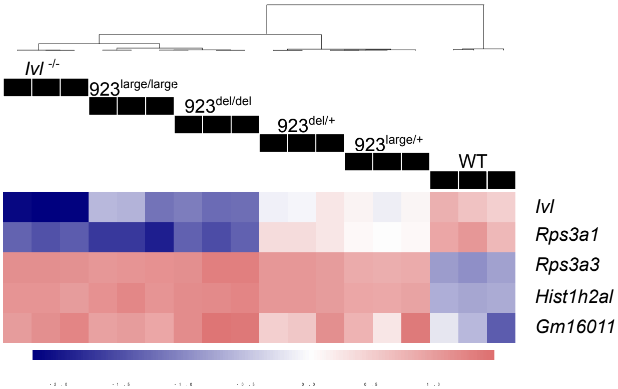

Supplementary Figure 3

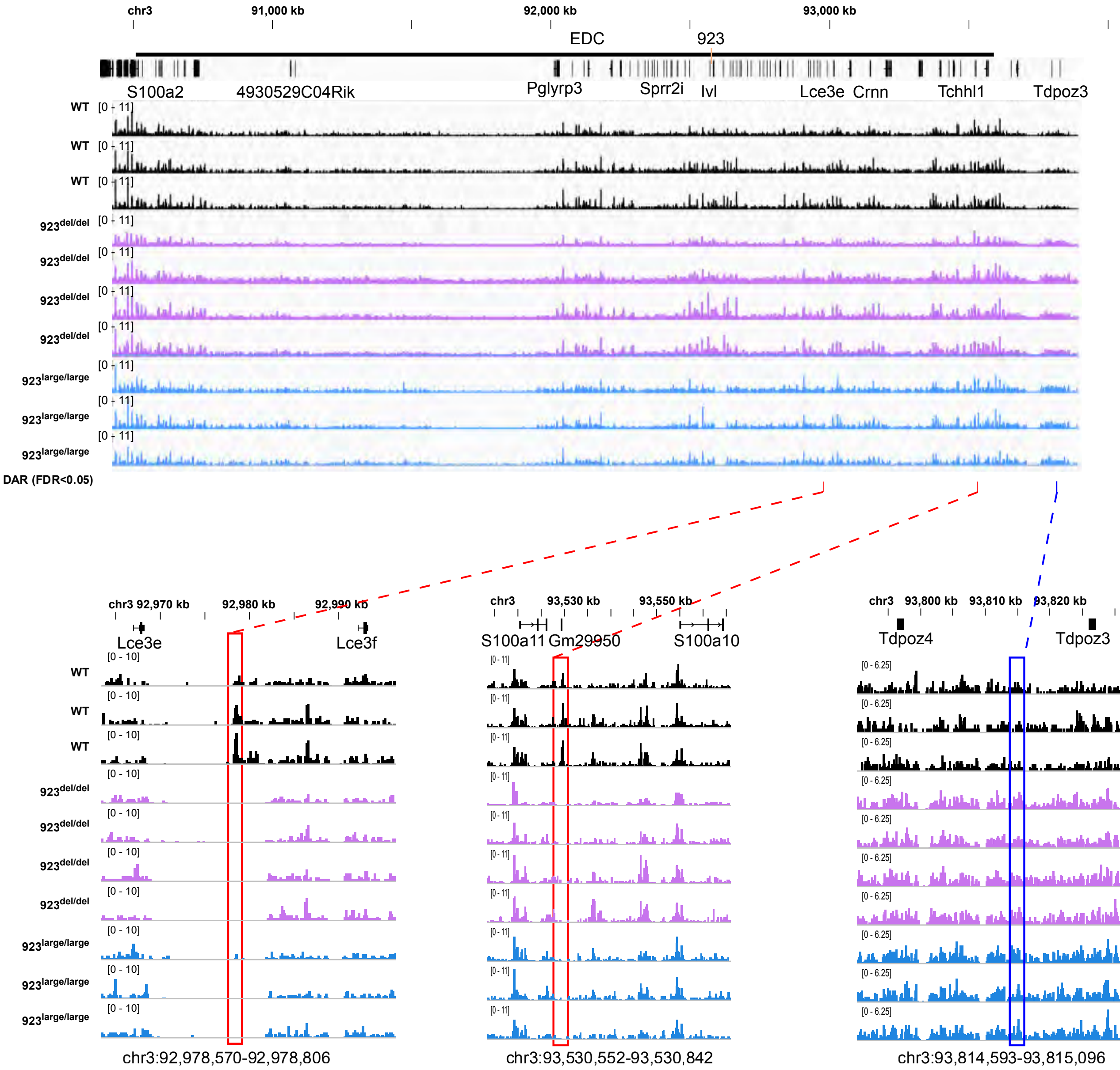

Supplementary Figure 4

a

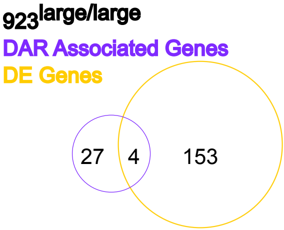


b

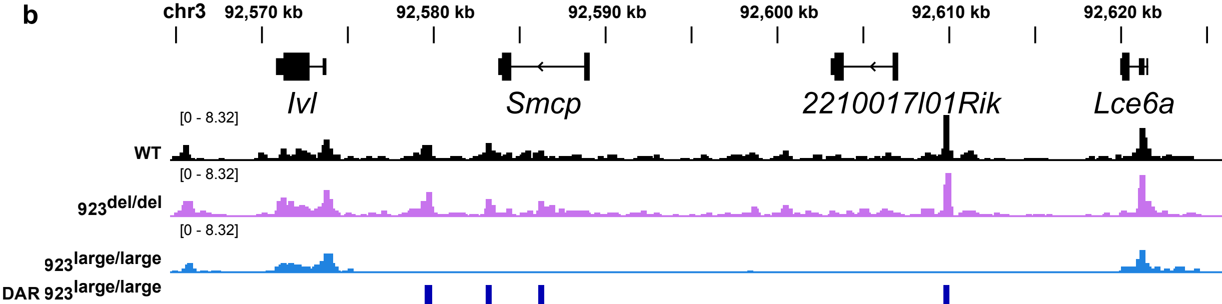

**Supplementary Figure 5**

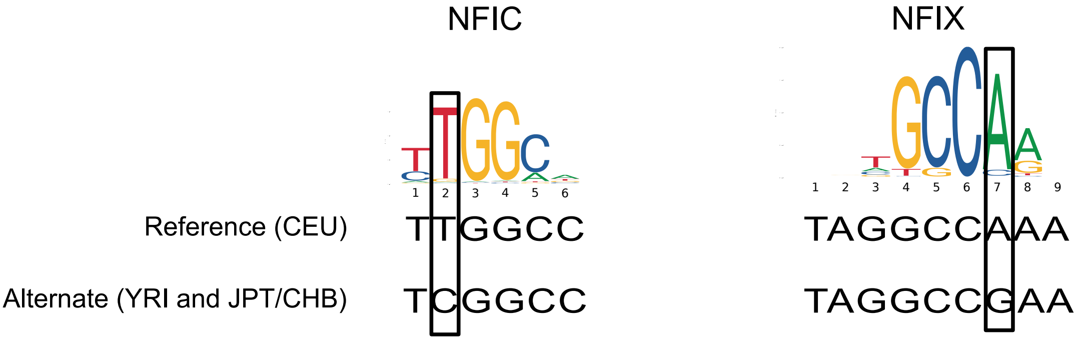

### Supplementary Figure 6

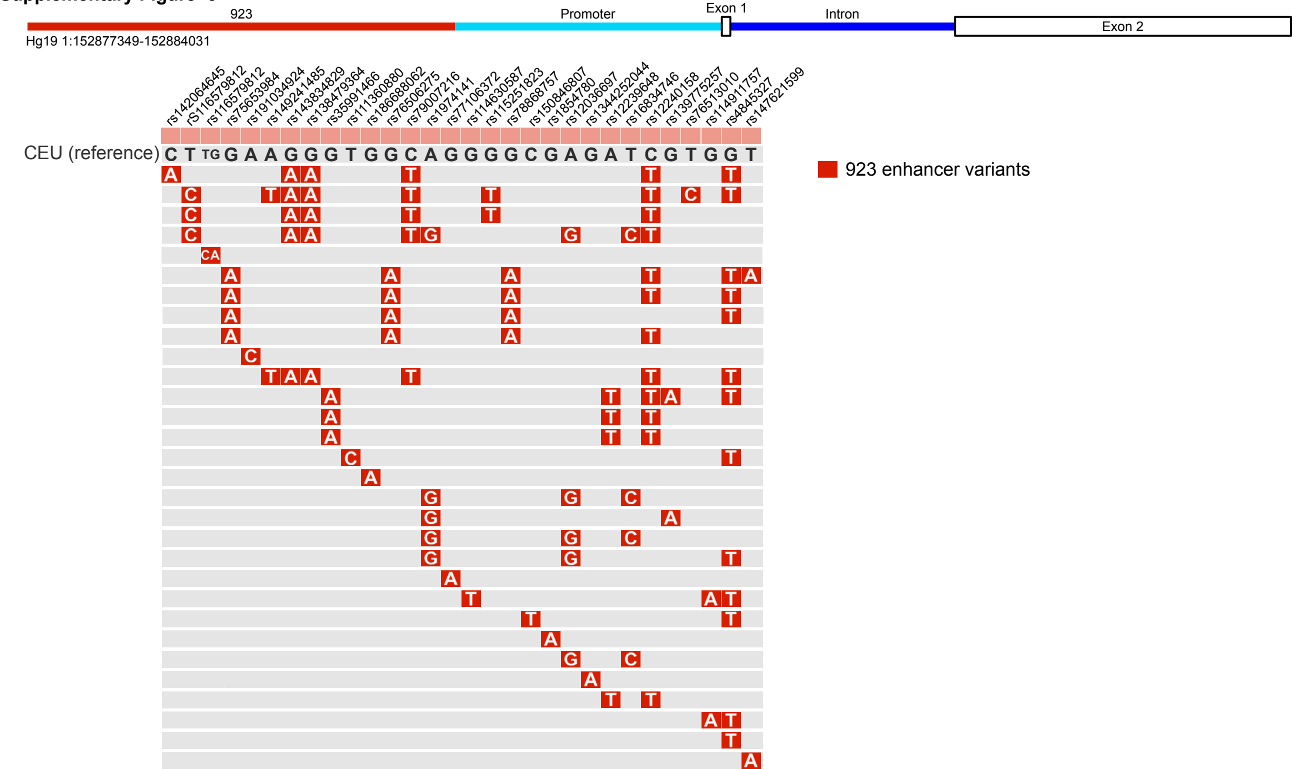
