## SupplementalTable1 for "An enhancer:involucrin regulatory module impacts human skin barrier adaptation out-of-Africa and modifies atopic dermatitis risk"

**Table S1. SNPs in clusters with high CMS scores (CMS>0) for each population.**

| *Population* | *SNP ID* | *CMS score* | *Region* |
|---|---|---|---|
| CEU | rs11205062 | 0.940825834 | *LCE2D-LCE2C* |
| CEU | rs1412547 | 0.904393281 | *LCE2D-LCE2C* |
| CEU | rs1412546 | 0.384145022 | *LCE2D-LCE2C* |
| CEU | rs12022319 | 5.109347946 | *SMCP* |
| CEU | rs4845490 | 8.172431499 | *SMCP* |
| CEU | rs4845491 | 6.968754084 | *SMCP* |
| CEU | rs3737861 | 4.328669894 | *SMCP* |
| CEU | rs16834728 | 3.18479759 | *SMCP* |
| CEU | rs4845327 | 1.707098701 | *IVL* |
| CEU | rs1854779 | 1.987061717 | *IVL* |
| CEU | rs7528862 | 1.987061717 | *IVL* |
| CEU | rs7539232 | 1.987061717 | *IVL* |
| CEU | rs11205132 | 1.987061717 | *IVL* |
| CEU | rs2229496 | 1.987061717 | *IVL* |
| CEU | rs7535306 | 1.987061717 | *IVL* |
| CEU | rs7545520 | 1.987061717 | *IVL* |
| CEU | rs55780839 | 1.224517473 | *SPRR2E-SPRR2F* |
| CEU | rs310128 | 0.483971238 | *SPRR2E-SPRR2F* |
| CEU | rs28410699 | 1.099842733 | *S100A6* |
| CEU | rs6587745 | 1.970059831 | *S100A6* |
| CEU | rs6587746 | 1.970059831 | *S100A6* |
| CEU | rs16835393 | 2.460390164 | *S100A6* |
| CEU | rs4845554 | 2.460390164 | *S100A6* |
| JPT/CHB | rs74127901 | 1.646499519 | *HRNR-FLG* |
| JPT/CHB | rs12748016 | 1.047502908 | *HRNR-FLG* |
| JPT/CHB | rs72696975 | 3.192900821 | *HRNR-FLG* |
| JPT/CHB | rs4511111 | 5.234197413 | *HRNR-FLG* |
| JPT/CHB | rs991231 | 2.246712246 | *HRNR-FLG* |
| JPT/CHB | rs11204948 | 0.699411258 | *HRNR-FLG* |
| JPT/CHB | rs11204949 | 1.693081979 | *HRNR-FLG* |
| JPT/CHB | rs11204971 | 0.631827205 | *HRNR-FLG* |
| JPT/CHB | rs4845767 | 0.763715164 | *CRNN-LCE5A* |
| JPT/CHB | rs6694145 | 1.013182206 | *CRNN-LCE5A* |
| JPT/CHB | rs6668295 | 2.027376788 | *IVL* |
| JPT/CHB | rs11205130 | 2.027376788 | *IVL* |
| JPT/CHB | rs4240864 | 0.798784213 | *LINC01527* |
| JPT/CHB | rs78730108 | 0.980450246 | *LINC01527* |
| YRI | rs11205031 | 0.572385476 | *5' of LCE3E* |
| YRI | rs11205032 | 0.572385476 | *5' of LCE3E* |
| YRI | rs11205033 | 0.80262217 | *5' of LCE3E* |
