## SupplementalTable2 for "An enhancer:involucrin regulatory module impacts human skin barrier adaptation out-of-Africa and modifies atopic dermatitis risk"

**Table S2. iSAFE scores for EDC SNPs in JPT/CHB.**

| *SNP ID* | *POS (hg19)* | *iSAFE* |
|---|---|---|
| rs71582209 | 151912718 | 0.000362735 |
| rs11589720 | 151921469 | 0.012363769 |
| rs61324873 | 151930351 | 0.001294859 |
| rs3007693 | 151932375 | 0.000234377 |
| rs562423432 | 151934243 | 0.012255957 |
| rs6662611 | 151936485 | 0.003634191 |
| rs2932564 | 151937894 | 0.000234377 |
| rs66914950 | 151938535 | 0.002964537 |
| rs6703772 | 151939856 | 0.0019977 |
| rs6691212 | 151939857 | 0.0019977 |
| rs6657658 | 151939864 | 0.0019977 |
| rs1552604 | 151940403 | 0.001765042 |
| rs11338554 | 151940738 | 0.000234377 |
| rs2999512 | 151946092 | 0.00825304 |
| rs112935859 | 151946656 | 0.021696458 |
| rs11204922 | 151961575 | 0.015953856 |
| rs200997618 | 151985014 | 0.002261026 |
| rs12410789 | 151985018 | 0.01016911 |
| rs11419775 | 152017637 | 0.003800938 |
| rs72696919 | 152021278 | 0.004069774 |
| rs138115683 | 152021391 | 0.004113482 |
| rs4333848 | 152028561 | 0.005854997 |
| rs3124315 | 152039711 | 0.00266311 |
| rs3124316 | 152042512 | 0.008005797 |
| rs563142014 | 152049767 | 0.00430791 |
| rs11204925 | 152073120 | 0.010108306 |
| rs3001974 | 152077574 | 0.006561969 |
| rs2935208 | 152077859 | 0.006561969 |
| rs1131471 | 152079989 | 0.006561969 |
| rs2496253 | 152081921 | 0.006561969 |
| rs112584962 | 152117599 | 0.005995556 |
| rs11204926 | 152125626 | 0.003579695 |
| rs3001978 | 152126467 | 0.00738698 |
| rs78544048 | 152129087 | 0.005484651 |
| rs76015112 | 152129094 | 0.005484651 |
| rs924086 | 152139660 | 0.005995556 |
| rs969359 | 152144260 | 0.011980286 |
| rs6696576 | 152157887 | 0.012826634 |
| rs11587676 | 152159961 | 0.025097311 |

| | | |
|---|---|---|
| rs1390489 | 152161103 | 0.064917565 |
| rs1390490 | 152161354 | 0.019207123 |
| rs11583524 | 152161657 | 0.024648598 |
| rs1496049 | 152161949 | 0.023418832 |
| rs372224762 | 152168479 | 0.00442125 |
| rs149885335 | 152170141 | 0.003218552 |
| rs28682375 | 152170149 | 0.003218552 |
| rs12408581 | 152175211 | 0.027132557 |
| rs12409762 | 152175398 | 0.022873097 |
| rs56013982 | 152176503 | 0.02614517 |
| rs11589532 | 152176539 | 0.023967401 |
| rs11204934 | 152176848 | 0.023967401 |
| rs868304 | 152178568 | 0.018648684 |
| rs12567463 | 152180128 | 0.023967401 |
| rs72696963 | 152181189 | 0.026830664 |
| rs4415568 | 152181207 | 0.018304118 |
| rs1390485 | 152182168 | 0.022873097 |
| rs11589587 | 152184129 | 0.027132557 |
| rs12726613 | 152185500 | 0.022873097 |
| rs12729662 | 152185750 | 0.023976039 |
| rs71625163 | 152190682 | 0.020594488 |
| rs74127901 | 152195579 | 0.025288617 |
| rs72696969 | 152196339 | 0.027789377 |
| rs12748016 | 152197235 | 0.023428115 |
| rs7534716 | 152198654 | 0.018216901 |
| rs10888470 | 152200472 | 0.018216901 |
| rs72696975 | 152206090 | 0.027789377 |
| rs10888472 | 152206896 | 0.026002739 |
| rs4511111 | 152211624 | 0.027789377 |
| rs991231 | 152211898 | 0.023428115 |
| rs12030180 | 152214406 | 0.011076695 |
| rs6666382 | 152217226 | 0.023428115 |
| rs11204947 | 152218257 | 0.023428115 |
| rs11204948 | 152218473 | 0.0258683 |
| rs11204949 | 152219797 | 0.027789377 |
| rs55957623 | 152223196 | 0.027789377 |
| rs6668499 | 152228927 | 0.027789377 |
| rs4582793 | 152229578 | 0.027789377 |
| rs200323326 | 152230104 | 0.027177951 |
| rs55654502 | 152231877 | 0.027789377 |
| rs12406105 | 152232635 | 0.027789377 |

| | | |
|---|---|---|
| rs12566867 | 152234888 | 0.027789377 |
| rs12408450 | 152238732 | 0.027789377 |
| rs12408186 | 152238785 | 0.02193372 |
| rs77937913 | 152238789 | 0.02193372 |
| rs12407905 | 152240439 | 0.022013092 |
| rs12407913 | 152240471 | 0.027402043 |
| rs12406777 | 152240472 | 0.027402043 |
| rs60742682 | 152246485 | 0.028302172 |
| rs72696989 | 152248674 | 0.027789377 |
| rs6670717 | 152249956 | 0.025684476 |
| rs72696991 | 152250235 | 0.027789377 |
| rs10888483 | 152252789 | 0.027789377 |
| rs11485508 | 152253167 | 0.003516912 |
| rs112311505 | 152253539 | 0.027789377 |
| rs12027807 | 152253726 | 0.003516912 |
| rs75533785 | 152253781 | 0.027789377 |
| rs113085728 | 152254010 | 0.027789377 |
| rs139965765 | 152254102 | 0.027789377 |
| rs148255037 | 152255714 | 0.025478793 |
| rs3126043 | 152256030 | 0.003679969 |
| rs72696997 | 152257099 | 0.027789377 |
| rs11204971 | 152259078 | 0.027789377 |
| rs1858477 | 152265261 | 0.003679969 |
| rs3126058 | 152269622 | 0.003193857 |
| rs144613541 | 152270875 | 0.00683659 |
| rs147652658 | 152270880 | 0.00683659 |
| rs3126060 | 152271307 | 0.00314081 |
| rs2184951 | 152273168 | 0.002059095 |
| rs12730241 | 152274016 | 0.002059095 |
| rs3126075 | 152276626 | 0.002305419 |
| rs531858654 | 152276823 | 0.004332663 |
| rs547915245 | 152276831 | 0.004332663 |
| rs3126066 | 152276871 | 0.002633292 |
| rs3091276 | 152277168 | 0.003491876 |
| rs9436066 | 152277717 | 0.00250463 |
| rs2065957 | 152277826 | 0.001608104 |
| rs2065958 | 152278049 | 0.002105027 |
| rs57672167 | 152278689 | 0.002267127 |
| rs192116923 | 152279406 | 0.003197711 |
| rs3126072 | 152279729 | 0.003329297 |
| rs3126074 | 152279841 | 0.003329297 |

| | | |
|---|---|---|
| rs11586631 | 152283283 | 0.027789377 |
| rs74129461 | 152285099 | 0.0258683 |
| rs12036682 | 152285807 | 0.010473399 |
| rs11584340 | 152285930 | 0.024533711 |
| rs11588170 | 152286032 | 0.027789377 |
| rs41267154 | 152286367 | 0.024533711 |
| rs3120659 | 152293202 | 0.003405591 |
| rs34188926 | 152293481 | 0.023488376 |
| rs3120661 | 152294402 | 0.003757001 |
| rs1858479 | 152294823 | 0.023488376 |
| rs12407156 | 152295942 | 0.02671193 |
| rs11204980 | 152296191 | 0.002314965 |
| rs34047456 | 152296573 | 0.002314965 |
| rs6587666 | 152296863 | 0.002314965 |
| rs6681457 | 152297117 | 0.024934274 |
| rs12407553 | 152297517 | 0.023488376 |
| rs11204981 | 152298212 | 0.002314965 |
| rs11586114 | 152299767 | 0.0263354 |
| rs11584427 | 152299788 | 0.02671193 |
| rs568564998 | 152302112 | 0.025063184 |
| rs72698906 | 152307184 | 0.025574305 |
| rs3126099 | 152314219 | 0.003718857 |
| rs3120665 | 152316590 | 0.003685254 |
| rs11588022 | 152316952 | 0.013822903 |
| rs10712489 | 152318040 | 0.003685254 |
| rs3120667 | 152318161 | 0.003685254 |
| rs72208466 | 152320127 | 0.003685254 |
| rs11204982 | 152322042 | 0.003685254 |
| rs6679449 | 152328411 | 0.004155004 |
| rs10788835 | 152330945 | 0.004155004 |
| rs3818831 | 152331240 | 0.003685254 |
| rs2275264 | 152331533 | 0.004155004 |
| rs11204984 | 152331987 | 0.00423666 |
| rs1858485 | 152334530 | 0.003793814 |
| rs72155677 | 152335006 | 0.003395203 |
| rs35385577 | 152337325 | 0.003756807 |
| rs12402680 | 152342904 | 0.003685254 |
| rs12029633 | 152347403 | 0.004037695 |
| rs10788836 | 152350135 | 0.004037695 |
| rs10888484 | 152352514 | 0.004037695 |
| rs4322234 | 152353768 | 0.004198769 |

| | | |
|---|---|---|
| rs6671146 | 152354472 | 0.004198769 |
| rs11204994 | 152358640 | 0.004198769 |
| rs1923506 | 152359700 | 0.004198769 |
| rs1923503 | 152360820 | 0.004198769 |
| rs7534951 | 152362972 | 0.004198769 |
| rs7514696 | 152363874 | 0.004198769 |
| rs11204998 | 152365655 | 0.004198769 |
| rs67661262 | 152368016 | 0.014016708 |
| rs12408401 | 152371801 | 0.014016708 |
| rs6660866 | 152372723 | 0.014343661 |
| rs202241217 | 152373605 | 0.014343661 |
| rs562029208 | 152375309 | 0.004299214 |
| rs151250169 | 152375430 | 0.014343661 |
| rs140242944 | 152380846 | 0.009821683 |
| rs12024486 | 152381085 | 0.009525764 |
| rs10888486 | 152382062 | 0.009525764 |
| rs12029431 | 152391547 | 0.008736315 |
| rs11590822 | 152392959 | 0.008736315 |
| rs6679899 | 152393847 | 0.008736315 |
| rs11583558 | 152397093 | 0.008736315 |
| rs726863 | 152400361 | 0.006947304 |
| rs11205002 | 152403339 | 0.008064809 |
| rs12749505 | 152404073 | 0.008064809 |
| rs113419595 | 152404275 | 0.008064809 |
| rs986903 | 152410569 | 0.008064809 |
| rs10888488 | 152411090 | 0.008064809 |
| rs6695950 | 152412687 | 0.008064809 |
| rs1923499 | 152413175 | 0.005452779 |
| rs4845433 | 152414439 | 0.010408948 |
| rs4240878 | 152414833 | 0.014441222 |
| rs4557976 | 152415066 | 0.008889926 |
| rs4845767 | 152415308 | 0.008889926 |
| rs4845768 | 152415473 | 0.007276978 |
| rs11205003 | 152416630 | 0.008093697 |
| rs141355348 | 152418162 | 0.005067258 |
| rs12125681 | 152418213 | 0.007276978 |
| rs1885527 | 152421545 | 0.006971554 |
| rs4845435 | 152423797 | 0.013313776 |
| rs6694145 | 152424825 | 0.008889926 |
| rs7531606 | 152426870 | 0.008889926 |
| rs6587673 | 152430152 | 0.011993154 |

| | | |
|---|---|---|
| rs11461245 | 152432643 | 0.006167098 |
| rs17597997 | 152432884 | 0.011993154 |
| rs12137893 | 152437397 | 0.01126446 |
| rs12116760 | 152437883 | 0.011993154 |
| rs952180 | 152439945 | 0.01126446 |
| rs6704503 | 152441295 | 0.01126446 |
| rs13373771 | 152443608 | 0.011083136 |
| rs34406913 | 152443670 | 0.011902594 |
| rs77033261 | 152444584 | 0.012033055 |
| rs61815580 | 152444619 | 0.01032022 |
| rs12733078 | 152444658 | 0.012033055 |
| rs61813862 | 152445162 | 0.013589968 |
| rs4845773 | 152445418 | 0.013589968 |
| rs4845774 | 152445621 | 0.011340885 |
| rs4845439 | 152446495 | 0.012265041 |
| rs6698520 | 152448967 | 0.012092207 |
| rs4845777 | 152450239 | 0.0133919 |
| rs4240882 | 152456358 | 0.021142728 |
| rs11205010 | 152457964 | 0.022505484 |
| rs112791653 | 152467061 | 0.020880315 |
| rs2181173 | 152474866 | 0.026353018 |
| rs7535188 | 152475899 | 0.025248529 |
| rs6587679 | 152481262 | 0.031461336 |
| rs11581947 | 152482587 | 0.026391206 |
| rs11205014 | 152483634 | 0.031678396 |
| rs2105117 | 152484129 | 0.027933751 |
| rs3753451 | 152485227 | 0.024095496 |
| rs4845441 | 152485421 | 0.027933751 |
| rs10666468 | 152485608 | 0.024095496 |
| rs6671924 | 152485830 | 0.027933751 |
| rs2282296 | 152487979 | 0.034092569 |
| rs4845780 | 152488846 | 0.034092569 |
| rs548252 | 152489742 | 0.034092569 |
| rs2181172 | 152491794 | 0.034092569 |
| rs12132469 | 152491856 | 0.034092569 |
| rs10888491 | 152492622 | 0.032733292 |
| rs11365304 | 152493337 | 0.020954093 |
| rs493133 | 152493875 | 0.034092569 |
| rs1001834 | 152495940 | 0.034092569 |
| rs11205018 | 152496249 | 0.034092569 |
| rs545418 | 152497338 | 0.034092569 |

| | | |
|---|---|---|
| rs12116609 | 152497685 | 0.034092569 |
| rs526099 | 152497811 | 0.034092569 |
| rs525960 | 152497866 | 0.034092569 |
| rs11584191 | 152500428 | 0.034092569 |
| rs71582241 | 152504491 | 0.035053533 |
| rs11404509 | 152506888 | 0.034658687 |
| rs61291402 | 152506889 | 0.034658687 |
| rs10666767 | 152506982 | 0.034314351 |
| rs4845443 | 152507156 | 0.034495755 |
| rs4845785 | 152510143 | 0.033645218 |
| rs4845788 | 152519496 | 0.031245575 |
| rs75513934 | 152522072 | 0.026298728 |
| rs144873910 | 152522469 | 0.026298728 |
| rs138657579 | 152522870 | 0.030651546 |
| rs12023930 | 152523949 | 0.027221093 |
| rs12034882 | 152525492 | 0.031245575 |
| rs4240883 | 152526260 | 0.031245575 |
| rs11205031 | 152527662 | 0.030045474 |
| rs11205032 | 152527690 | 0.030045474 |
| rs11205033 | 152527966 | 0.031245575 |
| rs7531317 | 152528439 | 0.027493768 |
| rs4240885 | 152529850 | 0.031245575 |
| rs4845444 | 152530125 | 0.031245575 |
| rs71626721 | 152532745 | 0.026054132 |
| rs11205035 | 152532832 | 0.031345698 |
| rs4112785 | 152533115 | 0.031345698 |
| rs5777817 | 152534105 | 0.049799766 |
| rs4074613 | 152534191 | 0.040455659 |
| rs79231283 | 152542115 | 0.008431342 |
| rs150573777 | 152543446 | 0.00097078 |
| rs10888502 | 152545993 | 0.041933001 |
| rs182958039 | 152553727 | 0.00097078 |
| rs74443569 | 152570460 | 0.00097078 |
| rs75196578 | 152588529 | 0.002345629 |
| rs200733667 | 152589240 | 0.010609839 |
| rs1581804 | 152592262 | 0.051849195 |
| rs11205045 | 152594826 | 0.054361924 |
| rs12041270 | 152596920 | 0.054255262 |
| rs11205046 | 152597043 | 0.054255262 |
| rs9970657 | 152597192 | 0.054255262 |
| rs1987306 | 152597804 | 0.054255262 |

| | | |
|---|---|---|
| rs1474169 | 152598168 | 0.054255262 |
| rs925977 | 152598977 | 0.054255262 |
| rs35655600 | 152599002 | 0.05264865 |
| rs11205048 | 152601676 | 0.054255262 |
| rs12564254 | 152602512 | 0.054255262 |
| rs11581621 | 152603011 | 0.054255262 |
| rs11587218 | 152603091 | 0.05713263 |
| rs11205050 | 152603441 | 0.054255262 |
| rs12073434 | 152603776 | 0.054255262 |
| rs373516735 | 152604900 | 0.051801624 |
| rs1575754 | 152605117 | 0.051801624 |
| rs12035776 | 152607461 | 0.054255262 |
| rs10127582 | 152608787 | 0.054548469 |
| rs7522480 | 152608876 | 0.067406374 |
| rs10127681 | 152608912 | 0.053511452 |
| rs6671975 | 152613631 | 0.067406374 |
| rs138421494 | 152619520 | 0.031316569 |
| rs66587527 | 152619643 | 0.044932599 |
| rs7540520 | 152619739 | 0.074335934 |
| rs10888505 | 152625148 | 0.05057526 |
| rs4287181 | 152626170 | 0.067406374 |
| rs373465501 | 152626743 | 0.054812738 |
| rs6587691 | 152634672 | 0.066915984 |
| rs1855080 | 152637150 | 0.066915984 |
| rs1855079 | 152638134 | 0.066915984 |
| rs11205065 | 152643998 | 0.067957158 |
| rs1930126 | 152647579 | 0.066915984 |
| rs1853805 | 152649114 | 0.066915984 |
| rs2014369 | 152651023 | 0.062175124 |
| rs2014368 | 152651029 | 0.062175124 |
| rs7546815 | 152651851 | 0.062175124 |
| rs12097462 | 152652137 | 0.066915984 |
| rs10788844 | 152653325 | 0.062175124 |
| rs1925663 | 152653506 | 0.061646737 |
| rs11205073 | 152654385 | 0.062175124 |
| rs1332507 | 152660493 | 0.062175124 |
| rs1332508 | 152660503 | 0.062175124 |
| rs1332509 | 152660746 | 0.062175124 |
| rs7525190 | 152663352 | 0.066915984 |
| rs10465797 | 152665640 | 0.062175124 |
| rs10788845 | 152668072 | 0.062175124 |

| | | |
|---|---|---|
| rs3904413 | 152668498 | 0.066915984 |
| rs11205078 | 152669459 | 0.062175124 |
| rs11581506 | 152673259 | 0.064451413 |
| rs11587581 | 152674066 | 0.064451413 |
| rs7543194 | 152674220 | 0.066915984 |
| rs7539239 | 152677727 | 0.064451413 |
| rs1412542 | 152678445 | 0.064451413 |
| rs1536157 | 152686773 | 0.067430051 |
| rs4845466 | 152690745 | 0.079633312 |
| rs4845467 | 152690825 | 0.079633312 |
| rs4845468 | 152691078 | 0.079633312 |
| rs4845469 | 152691308 | 0.079633312 |
| rs7545543 | 152691367 | 0.079633312 |
| rs7534164 | 152691438 | 0.079633312 |
| rs1332499 | 152691943 | 0.079633312 |
| rs1332500 | 152692074 | 0.079633312 |
| rs873775 | 152692472 | 0.079633312 |
| rs944682 | 152692588 | 0.079633312 |
| rs3814352 | 152692948 | 0.079633312 |
| rs3814353 | 152692952 | 0.079633312 |
| rs11205087 | 152695109 | 0.078687583 |
| rs10888511 | 152697881 | 0.067430051 |
| rs1888963 | 152701241 | 0.07116871 |
| rs10888514 | 152707929 | 0.073745383 |
| rs66659820 | 152719538 | 0.042442244 |
| rs28464852 | 152721454 | 0.045201221 |
| rs9729955 | 152721468 | 0.042002667 |
| rs1332506 | 152724053 | 0.039581526 |
| rs7515795 | 152725673 | 0.045271539 |
| rs10788848 | 152725723 | 0.045271539 |
| rs4845477 | 152727544 | 0.045271539 |
| rs4845478 | 152727803 | 0.042115965 |
| rs882827 | 152728795 | 0.045271539 |
| rs4845479 | 152730216 | 0.042115965 |
| rs17612167 | 152732106 | 0.008587329 |
| rs6673149 | 152734633 | 0.043645017 |
| rs11205099 | 152735155 | 0.045271539 |
| rs6680648 | 152737228 | 0.045271539 |
| rs6587702 | 152737703 | 0.045271539 |
| rs75366542 | 152738368 | 0.033472893 |
| rs4845481 | 152739631 | 0.045271539 |

| | | |
|---|---|---|
| rs6587703 | 152740858 | 0.032018981 |
| rs12077214 | 152743849 | 0.04496242 |
| **rs143284883** | **152745715** | **0.081453854** |
| **rs34384421** | **152746417** | **0.081453854** |
| rs11205102 | 152747126 | 0.044008315 |
| rs2339382 | 152749733 | 0.04496242 |
| rs913997 | 152750256 | 0.031574435 |
| **rs10157301** | **152751595** | **0.106244029** |
| rs11205103 | 152754916 | 0.04496242 |
| rs950338 | 152756806 | 0.040980475 |
| rs7530000 | 152757690 | 0.042576607 |
| rs11576947 | 152758519 | 0.041569783 |
| rs10888516 | 152759010 | 0.041569783 |
| rs1034109 | 152759678 | 0.041569783 |
| rs16834535 | 152761091 | 0.025354609 |
| rs1831639 | 152761454 | 0.025324947 |
| rs6587704 | 152761709 | 0.025324947 |
| rs4845485 | 152766239 | 0.024961018 |
| rs1537307 | 152769807 | 0.0246425 |
| rs41268492 | 152770521 | 0.017575604 |
| rs115138530 | 152770828 | 0.028686451 |
| rs2065206 | 152771293 | 0.020864249 |
| rs6587705 | 152774367 | 0.036634647 |
| rs11322194 | 152774515 | 0.020864249 |
| **rs1048535** | **152777465** | **0.077541958** |
| **rs35436039** | **152778118** | **0.082369934** |
| **rs4845488** | **152778576** | **0.063269509** |
| **rs1930127** | **152780277** | **0.100160336** |
| rs7524281 | 152783255 | 0.022887091 |
| rs12023196 | 152783724 | 0.030899517 |
| **rs11804609** | **152784255** | **0.121767753** |
| rs4845325 | 152798560 | 0.042989863 |
| rs11205114 | 152801416 | 0.005501101 |
| rs4240863 | 152843377 | 0.045751317 |
| rs12047888 | 152846621 | 0.047667711 |
| rs16834734 | 152853278 | 0.047667711 |
| rs377207121 | 152854429 | 0.047667711 |
| rs2339385 | 152859594 | 0.073494793 |
| rs547649180 | 152860981 | 0.051126286 |
| rs1590931 | 152861866 | 0.009106587 |
| rs6587709 | 152862466 | 0.073842659 |

| | | |
|---|---|---|
| rs1330703 | 152867349 | 0.009945117 |
| rs6668295 | 152869494 | 0.045998876 |
| rs58540020 | 152869713 | 0.034405125 |
| rs3856026 | 152869763 | 0.075911206 |
| rs75315086 | 152870259 | 0.034405125 |
| rs6671493 | 152870341 | 0.009945117 |
| rs6587710 | 152870716 | 0.01052925 |
| rs76570087 | 152873253 | 0.034405125 |
| rs77171551 | 152873859 | 0.034405125 |
| rs11804354 | 152874445 | 0.034405125 |
| rs79428665 | 152875207 | 0.038530312 |
| rs58290373 | 152875358 | 0.048282176 |
| rs1974141 | 152878222 | 0.048158566 |
| rs12036697 | 152878909 | 0.048158566 |
| rs2879484 | 152885500 | 0.043806241 |
| rs2879485 | 152885913 | 0.043806241 |
| rs7539610 | 152889327 | 0.032985558 |
| rs12239808 | 152889944 | 0.043806241 |
| rs12239812 | 152890078 | 0.043575629 |
| rs12239814 | 152890138 | 0.043806241 |
| rs11806470 | 152890604 | 0.03767148 |
| rs10788849 | 152902240 | 0.006788446 |
| rs4845497 | 152904162 | 0.006788446 |
| rs61614885 | 152906032 | 0.014566939 |
| rs57589052 | 152906033 | 0.014566939 |
| rs756303 | 152908222 | 0.007399796 |
| rs58298199 | 152908814 | 0.006021588 |
| rs71730778 | 152909296 | 0.006003999 |
| rs2050674 | 152913356 | 0.007399796 |
| rs34563282 | 152921985 | 0.009864223 |
| rs2066004 | 152922896 | 0.009865195 |
| rs2066005 | 152923113 | 0.011499072 |
| rs2050673 | 152923698 | 0.007969896 |
| rs4240865 | 152954839 | 0.022358481 |
| rs4240866 | 152954875 | 0.020339751 |
| rs1415964 | 152955085 | 0.022358481 |
| rs1999886 | 152955522 | 0.020339751 |
| rs1611759 | 152957439 | 0.020339751 |
| rs1611760 | 152957573 | 0.020339751 |
| rs1129654 | 152958238 | 0.020339751 |
| rs1129655 | 152958267 | 0.020339751 |

| | | |
|---|---|---|
| rs1984198 | 152958584 | 0.020339751 |
| rs1415962 | 152958731 | 0.018352234 |
| rs4255332 | 152958745 | 0.020407759 |
| rs6665575 | 152960506 | 0.020339751 |
| rs1577964 | 152961784 | 0.020339751 |
| rs1933383 | 152963468 | 0.020339751 |
| rs10788850 | 152963932 | 0.020339751 |
| rs6587716 | 152966705 | 0.020339751 |
| rs6665527 | 152967009 | 0.020339751 |
| rs61811427 | 152968419 | 0.017427187 |
| rs6667418 | 152969685 | 0.021463132 |
| rs9919227 | 152970867 | 0.019545639 |
| rs885096 | 152971313 | 0.019545639 |
| rs946100 | 152971663 | 0.019545639 |
| rs4319267 | 152972699 | 0.019545639 |
| rs3964619 | 152972811 | 0.019545639 |
| rs1415961 | 152973464 | 0.019545639 |
| rs3753454 | 152973776 | 0.019545639 |
| rs946097 | 152974732 | 0.019545639 |
| rs946098 | 152975038 | 0.019545639 |
| rs10399896 | 152975083 | 0.019545639 |
| rs2339494 | 152975143 | 0.019545639 |
| rs1055935 | 152975941 | 0.019545639 |
| rs12047099 | 152981048 | 0.046209443 |
| rs12024694 | 152986260 | 0.046209443 |
| rs11205165 | 152986928 | 0.019545639 |
| rs12029192 | 152988118 | 0.046209443 |
| rs4363385 | 152989321 | 0.020003018 |
| rs16834871 | 152996161 | 0.049125126 |
| rs4845513 | 152998066 | 0.02610463 |
| rs2711 | 153005156 | 0.014906078 |
| rs423692 | 153005681 | 0.014906078 |
| rs489323 | 153006047 | 0.014906078 |
| rs1415969 | 153006448 | 0.014906078 |
| rs382292 | 153006641 | 0.014906078 |
| rs451939 | 153006877 | 0.014906078 |
| rs419721 | 153006902 | 0.014906078 |
| rs1846857 | 153012765 | 0.037884919 |
| rs2339501 | 153013948 | 0.045889696 |
| rs1995308 | 153014082 | 0.023833119 |
| rs1995307 | 153014311 | 0.046249759 |

| | | |
|---|---|---|
| rs28647391 | 153014845 | 0.048689375 |
| rs4041337 | 153018913 | 0.048689375 |
| rs3856020 | 153019087 | 0.048689375 |
| rs310125 | 153020264 | 0.037037705 |
| rs649269 | 153023070 | 0.034826807 |
| rs12046394 | 153024634 | 0.034826807 |
| rs12047187 | 153025087 | 0.037202065 |
| rs12047225 | 153025233 | 0.048824097 |
| rs12037754 | 153025721 | 0.048824097 |
| rs634639 | 153025933 | 0.034826807 |
| rs6661063 | 153036048 | 0.048911444 |
| rs310096 | 153036391 | 0.034153114 |
| rs310099 | 153039294 | 0.036395634 |
| rs35722864 | 153040505 | 0.048441584 |
| rs6660797 | 153041726 | 0.059783868 |
| rs559423296 | 153042009 | 0.046796911 |
| rs200671030 | 153042010 | 0.046796911 |
| rs442209 | 153045330 | 0.034141584 |
| rs11205183 | 153048524 | 0.044650467 |
| rs10888529 | 153050654 | 0.057647814 |
| rs6587726 | 153052384 | 0.055147153 |
| rs376463294 | 153052588 | 0.055066785 |
| rs141309445 | 153052608 | 0.057647814 |
| rs4041381 | 153054285 | 0.025820071 |
| rs61811903 | 153054458 | 0.057647814 |
| rs72480203 | 153056884 | 0.041743232 |
| rs310111 | 153057176 | 0.032544027 |
| rs310112 | 153057762 | 0.031833828 |
| rs310113 | 153057789 | 0.031833828 |
| rs310116 | 153059105 | 0.033785001 |
| rs310117 | 153059801 | 0.031833828 |
| rs310118 | 153060120 | 0.031833828 |
| rs310121 | 153062634 | 0.032041998 |
| rs71582280 | 153063460 | 0.058229633 |
| rs6587728 | 153064996 | 0.029270931 |
| rs3913438 | 153066822 | 0.033530337 |
| rs12040837 | 153068267 | 0.036506824 |
| rs7536587 | 153073838 | 0.034679582 |
| rs12041391 | 153073840 | 0.034679582 |
| rs59429171 | 153074053 | 0.034679582 |
| rs12032237 | 153075423 | 0.034679582 |

| | | |
|---|---|---|
| rs55780839 | 153075561 | 0.034679582 |
| rs12075236 | 153076144 | 0.034768155 |
| rs144248807 | 153077351 | 0.036506824 |
| rs1119652 | 153077977 | 0.036506824 |
| rs61204539 | 153078499 | 0.036506824 |
| rs146071525 | 153080470 | 0.036506824 |
| rs11205201 | 153082864 | 0.034530598 |
| rs11205203 | 153084086 | 0.036094586 |
| rs74133286 | 153085029 | 0.035276917 |
| rs12037255 | 153090030 | 0.037453026 |
| rs11205205 | 153091767 | 0.037453026 |
| rs61811924 | 153094478 | 0.026826788 |
| rs58665653 | 153098180 | 0.037453026 |
| rs76087522 | 153098743 | 0.037453026 |
| rs549161726 | 153101118 | 0.032877956 |
| rs12409742 | 153101353 | 0.037453026 |
| rs12408875 | 153101566 | 0.037453026 |
| rs80334179 | 153102506 | 0.037453026 |
| rs16834971 | 153102724 | 0.037453026 |
| rs904949 | 153103772 | 0.037453026 |
| rs12045045 | 153104035 | 0.037453026 |
| rs11205209 | 153104182 | 0.026826788 |
| rs2928 | 153112806 | 0.026233778 |
| rs10888533 | 153114916 | 0.026826788 |
| rs12049044 | 153115109 | 0.033011273 |
| rs552242532 | 153115147 | 0.026736731 |
| rs12046633 | 153115172 | 0.026736731 |
| rs12045873 | 153115200 | 0.027875179 |
| rs11205211 | 153115253 | 0.03089015 |
| rs533437 | 153119969 | 0.025623741 |
| rs510277 | 153122310 | 0.025442096 |
| rs516280 | 153122930 | 0.025442096 |
| rs564787 | 153125400 | 0.024040545 |
| rs499893 | 153127880 | 0.024040545 |
| rs474907 | 153128269 | 0.026033653 |
| rs539269 | 153132115 | 0.024040545 |
| rs75183495 | 153132669 | 0.024040545 |
| rs112951761 | 153133079 | 0.021926278 |
| rs509061 | 153133123 | 0.021926278 |
| rs546160 | 153133124 | 0.021926278 |
| rs12403756 | 153133281 | 0.024276925 |

| | | |
|---|---|---|
| rs12406257 | 153133328 | 0.023804437 |
| rs12403764 | 153133355 | 0.023804437 |
| rs476761 | 153134300 | 0.026033653 |
| rs1697431 | 153134910 | 0.024076588 |
| rs503914 | 153135119 | 0.026033653 |
| rs555926 | 153135459 | 0.025388075 |
| rs474200 | 153136053 | 0.02472625 |
| rs57634145 | 153139358 | 0.018827199 |
| rs575860 | 153139890 | 0.016421175 |
| rs138114182 | 153139960 | 0.027213991 |
| rs9970592 | 153140118 | 0.026568693 |
| rs12406869 | 153140157 | 0.035955178 |
| rs498131 | 153143504 | 0.020576548 |
| rs75479578 | 153168834 | 0.002159947 |
| rs1831238 | 153169522 | 0.001613385 |
| rs56934782 | 153170035 | 0.002227746 |
| rs61427785 | 153170305 | 0.001461888 |
| rs78724691 | 153170401 | 0.001461888 |
| rs79925447 | 153172005 | 0.001461888 |
| rs1329107 | 153172189 | 0.001461888 |
| rs76717215 | 153175318 | 0.001461888 |
| rs7534334 | 153177852 | 0.001966923 |
| rs11205219 | 153181916 | 0.007427993 |
| rs10888536 | 153182911 | 0.007427993 |
| rs10494292 | 153183845 | 0.007204179 |
| rs10788860 | 153184789 | 0.009333602 |
| rs11205223 | 153184930 | 0.007204179 |
| rs10788861 | 153185394 | 0.009333602 |
| rs11205224 | 153185549 | 0.007204179 |
| rs11205225 | 153185608 | 0.007204179 |
| rs10888538 | 153186267 | 0.007204179 |
| rs12061473 | 153187269 | 0.007204179 |
| rs10888539 | 153187595 | 0.007204179 |
| rs11205228 | 153187807 | 0.007204179 |
| rs11205230 | 153188032 | 0.007204179 |
| rs12069007 | 153188869 | 0.007204179 |
| rs71582292 | 153189098 | 0.007204179 |
| rs1410860 | 153189978 | 0.009333602 |
| rs12062833 | 153190376 | 0.007204179 |
| rs1360100 | 153192214 | 0.009394697 |
| rs943134 | 153197786 | 0.009052919 |

| | | |
|---|---|---|
| rs1998845 | 153199733 | 0.007221176 |
| rs10788863 | 153200307 | 0.007722931 |
| rs4845341 | 153202786 | 0.007732265 |
| rs7525529 | 153205285 | 0.009131337 |
| rs7525704 | 153205421 | 0.003876199 |
| rs6674985 | 153206993 | 0.009131337 |
| rs1410863 | 153207274 | 0.009131337 |
| rs1536823 | 153210182 | 0.008578777 |
| rs6587733 | 153211557 | 0.008578777 |
| rs4845532 | 153212490 | 0.008578777 |
| rs12135864 | 153213098 | 0.00561155 |
| rs12136892 | 153214192 | 0.005596686 |
| rs12126039 | 153214352 | 0.005833169 |
| rs7523592 | 153214524 | 0.005596686 |
| rs11205246 | 153217329 | 0.008049303 |
| rs1926234 | 153217977 | 0.005400612 |
| rs554457965 | 153219336 | 0.00383104 |
| rs7532289 | 153222657 | 0.006434859 |
| rs1329097 | 153224122 | 0.006434859 |
| rs10567874 | 153230481 | 0.005875086 |
| rs943967 | 153231876 | 0.006434859 |
| rs11205255 | 153234602 | 0.006434859 |
| rs10888546 | 153236684 | 0.006232876 |
| rs4585927 | 153237467 | 0.005875086 |
| rs6686895 | 153237922 | 0.005875086 |
| rs1410861 | 153238503 | 0.005875086 |
| rs10047202 | 153239426 | 0.005875086 |
| rs2094636 | 153239586 | 0.005875086 |
| rs2094637 | 153239921 | 0.006268862 |
| rs1928341 | 153240013 | 0.005875086 |
| rs10788864 | 153242805 | 0.005875086 |
| rs10888549 | 153249072 | 0.021304748 |
| rs11205261 | 153254767 | 0.022209778 |
| rs2916238 | 153260711 | 0.021304748 |
| rs3795386 | 153270392 | 0.003801506 |
| rs2771111 | 153270857 | 0.008444578 |
| rs1433678 | 153271068 | 0.009799059 |
| rs821431 | 153271524 | 0.00951615 |
| rs821428 | 153275506 | 0.008661659 |
| rs843971 | 153277423 | 0.009859773 |
| rs1655320 | 153279177 | 0.00962637 |

| | | |
|---|---|---|
| rs821424 | 153282892 | 0.010725005 |
| rs821420 | 153283978 | 0.010725005 |
| rs821419 | 153284200 | 0.011060502 |
| rs821418 | 153284423 | 0.010725005 |
| rs821416 | 153287366 | 0.010725005 |
| rs1094362 | 153290524 | 0.01178993 |
| rs2916233 | 153290695 | 0.009788713 |
| rs821414 | 153292639 | 0.011060502 |
| rs821411 | 153294077 | 0.011454577 |
| rs67812211 | 153294196 | 0.009788713 |
| rs35395413 | 153294355 | 0.009592674 |
| rs2916229 | 153294653 | 0.009788713 |
| rs821408 | 153296027 | 0.011454577 |
| rs821407 | 153297131 | 0.01178993 |
| rs1836118 | 153298525 | 0.01178993 |
| rs2771115 | 153299965 | 0.011454577 |
| rs2771116 | 153300351 | 0.01178993 |
| rs2771118 | 153301527 | 0.011454577 |
| rs60799710 | 153301674 | 0.010239213 |
| rs2771120 | 153302417 | 0.009936811 |
| rs1655310 | 153303930 | 0.01178993 |
| rs1754133 | 153304560 | 0.010725005 |
| rs1754132 | 153304768 | 0.011060502 |
| rs1655308 | 153304769 | 0.011060502 |
| rs151252426 | 153308203 | 0.0084955 |
| rs1754134 | 153310747 | 0.010787211 |
| rs3006449 | 153314347 | 0.005865208 |
| rs3006450 | 153314534 | 0.005865208 |
| rs3014861 | 153314822 | 0.005865208 |
| rs3014862 | 153314999 | 0.005865208 |
| rs735012 | 153316260 | 0.005865208 |
| rs3006452 | 153317370 | 0.005865208 |
| rs2916208 | 153320125 | 0.005865208 |
| rs2916205 | 153320755 | 0.005865208 |
| rs3806231 | 153321348 | 0.005865208 |
| rs2916202 | 153321810 | 0.005865208 |
| rs4845345 | 153324212 | 0.005714629 |
| rs12561811 | 153324548 | 0.006218839 |
| rs35364949 | 153324802 | 0.006037278 |
| rs10888558 | 153325130 | 0.006285033 |
| rs1347250 | 153325954 | 0.006183106 |

| | | |
|---|---|---|
| rs11580993 | 153339314 | 0.006478449 |
| rs35195593 | 153343717 | 0.012848694 |
| rs3006475 | 153344636 | 0.014831338 |
| rs2916191 | 153345911 | 0.012848694 |
| rs4772 | 153346263 | 0.012848694 |
| rs3014885 | 153352127 | 0.014831338 |
| rs374815343 | 153352260 | 0.01335451 |
| rs2916221 | 153352865 | 0.012848694 |
| rs2916217 | 153353198 | 0.012848694 |
| rs3006479 | 153355594 | 0.014831338 |
| rs3006482 | 153356497 | 0.017768755 |
| rs371154676 | 153356719 | 0.012286034 |
| rs34944983 | 153357697 | 0.014831338 |
| rs6659104 | 153362146 | 0.013716754 |
| rs3006488 | 153362507 | 0.017768755 |
| rs73018470 | 153369532 | 0.014854516 |
| rs61803340 | 153369833 | 0.014854516 |
| rs59678238 | 153370659 | 0.014854516 |
| rs113420049 | 153374406 | 0.014854516 |
| rs6673905 | 153376145 | 0.014854516 |
| rs6686657 | 153376207 | 0.014854516 |
| rs11205283 | 153376350 | 0.014854516 |
| rs11205286 | 153376743 | 0.014854516 |
| rs11580424 | 153377319 | 0.014854516 |
| rs74436639 | 153382458 | 0.014854516 |
| rs11590633 | 153382941 | 0.014854516 |
| rs61803116 | 153384716 | 0.014854516 |
| rs4845543 | 153385086 | 0.014854516 |
| rs3014812 | 153388881 | 0.018883955 |
| rs4845348 | 153390245 | 0.010877904 |
| rs3006412 | 153390542 | 0.012414232 |
| rs3006414 | 153391729 | 0.015509894 |
| rs3014822 | 153400479 | 0.016083728 |
| rs3006422 | 153400672 | 0.016083728 |
| rs3014823 | 153401109 | 0.016083728 |
| rs3006423 | 153401664 | 0.016083728 |
| rs77140629 | 153403487 | 0.01339277 |
| rs3006426 | 153408031 | 0.00283968 |
| rs3014831 | 153411229 | 0.00290854 |
| rs10888562 | 153418785 | 0.005762286 |
| rs6587740 | 153419033 | 0.005762286 |

| | | |
|---|---|---|
| rs3014832 | 153419639 | 0.003172312 |
| rs3014833 | 153419984 | 0.003172312 |
| rs4845545 | 153422325 | 0.005562193 |
| rs6700785 | 153425008 | 0.005923138 |
| rs4418538 | 153426161 | 0.005560066 |
| rs3014835 | 153427569 | 0.003172312 |
| rs875501 | 153428683 | 0.003172312 |
| rs3014836 | 153431200 | 0.003172312 |
| rs3014838 | 153432265 | 0.003172312 |
| rs3014839 | 153432740 | 0.003172312 |
| rs3006433 | 153433692 | 0.003172312 |
| rs11205297 | 153433696 | 0.013118924 |
| rs4240869 | 153435744 | 0.005471755 |
| rs3006435 | 153437003 | 0.005123441 |
| rs3014841 | 153437344 | 0.005436836 |
| rs12759195 | 153437423 | 0.004913784 |
| rs3006436 | 153438422 | 0.005123441 |
| rs2986208 | 153440689 | 0.005123441 |
| rs983844 | 153442499 | 0.005123441 |
| rs10888563 | 153445024 | 0.005471755 |
| rs10888564 | 153445100 | 0.005471755 |
| rs71584111 | 153450133 | 0.005001029 |
| rs148011727 | 153458931 | 0.00192365 |
| rs7535089 | 153459023 | 0.005116012 |
| rs71584114 | 153460118 | 0.004644344 |
| rs6421468 | 153463764 | 0.004329714 |
| rs6672390 | 153466704 | 0.006837518 |
| rs6660055 | 153467074 | 0.006685497 |
| rs16835366 | 153468106 | 0.006685497 |
| rs28707433 | 153469765 | 0.006685497 |
| rs4845548 | 153470167 | 0.006685497 |
| rs11392697 | 153471344 | 0.002857059 |
| rs4845549 | 153472285 | 0.008017094 |
| rs4845550 | 153472556 | 0.007198883 |
| rs16835382 | 153475660 | 0.007135181 |
| rs9436095 | 153479777 | 0.006870854 |
| rs28635377 | 153479846 | 0.005341433 |
| rs28632016 | 153479905 | 0.005341433 |
| rs4845552 | 153479998 | 0.005341433 |
| rs28410699 | 153482272 | 0.006956251 |
| rs6587745 | 153485284 | 0.006956251 |

| | | |
|---|---|---|
| rs6587746 | 153485359 | 0.006956251 |
| rs16835393 | 153487225 | 0.006956251 |
| rs7544359 | 153487714 | 0.006656555 |
| rs7515940 | 153488027 | 0.006956251 |
| rs7537898 | 153488080 | 0.006956251 |
| rs7534823 | 153490810 | 0.006956251 |
| rs4845554 | 153492017 | 0.006956251 |
| rs7518306 | 153496247 | 0.01055791 |
| rs4845351 | 153496711 | 0.004814059 |
| rs477339 | 153510621 | 0.020082714 |
| rs1810765 | 153515120 | 0.02575354 |
| rs35932716 | 153535604 | 0.001818009 |
| rs144816475 | 153553932 | 0.003169118 |
| rs28857273 | 153560040 | 0.004884919 |
| rs2026564 | 153565636 | 0.005824511 |
| rs555939948 | 153567927 | 0.001429596 |
| rs9729962 | 153568986 | 0.005630574 |
| rs73009940 | 153569599 | 0.006481974 |
| rs28737329 | 153571023 | 0.007094086 |
| rs139734691 | 153572182 | 0.003241445 |
| rs148716453 | 153573725 | 0.001842747 |
| rs372416236 | 153574152 | 0.002353134 |
| rs9792963 | 153588208 | 0.000448029 |
| rs9792967 | 153588279 | 0.000448029 |
| rs558422909 | 153593635 | 0.000376989 |
