## SupplementalTable3 for "An enhancer:involucrin regulatory module impacts human skin barrier adaptation out-of-Africa and modifies atopic dermatitis risk"

**Table S3. iSAFE scores for EDC SNPs in CEU.**

| *SNP ID* | *POS (hg19)* | *iSAFE* |
|---|---|---|
| rs56313205 | 151904109 | 0.000182555 |
| rs6662602 | 151915155 | 0.000169534 |
| rs6663064 | 151915541 | 0.000169937 |
| rs61817722 | 151920862 | 0.000243151 |
| rs61817723 | 151920865 | 0.000188038 |
| rs369308985 | 151920971 | 0.000367134 |
| rs12095483 | 151921974 | 0.000243151 |
| rs61817725 | 151923177 | 0.000273378 |
| rs2932581 | 151933963 | 0.006230774 |
| rs562423432 | 151934243 | 0.006153104 |
| rs2999508 | 151934447 | 0.006230774 |
| rs11582739 | 151934541 | 0.000215814 |
| rs7554859 | 151937938 | 0.000215814 |
| rs1552604 | 151940403 | 0.007169235 |
| rs1038747 | 151942345 | 0.006230774 |
| rs2999512 | 151946092 | 0.010934301 |
| rs112935859 | 151946656 | 0.017035586 |
| rs3007702 | 151951600 | 0.012851222 |
| rs3007703 | 151951601 | 0.012851222 |
| rs2999526 | 151951695 | 0.007055298 |
| rs2999527 | 151951908 | 0.012851222 |
| rs2932592 | 151952065 | 0.012851222 |
| rs3007704 | 151952075 | 0.012851222 |
| rs2999528 | 151952885 | 0.011658907 |
| rs2932591 | 151953046 | 0.012851222 |
| rs2932590 | 151953763 | 0.011658907 |
| rs2999529 | 151953840 | 0.012851222 |
| rs11435688 | 151954033 | 0.012851222 |
| rs2999530 | 151954377 | 0.012851222 |
| rs2338019 | 151956814 | 0.011658907 |
| rs11204922 | 151961575 | 0.006293324 |
| rs11459749 | 151965357 | 0.012851222 |
| rs11542015 | 151966348 | 0.011658907 |
| rs3007708 | 151967836 | 0.011658907 |
| rs12736889 | 151968892 | 0.003813073 |
| rs2999531 | 151971497 | 0.007055298 |
| rs2999558 | 151972510 | 0.012851222 |
| rs882100 | 151977061 | 0.023147735 |

| | | |
|---|---|---|
| rs882099 | 151977172 | 0.023147735 |
| rs6587641 | 151978039 | 0.023147735 |
| rs2999561 | 151984547 | 0.007129832 |
| rs2932572 | 151984556 | 0.023147735 |
| rs12410789 | 151985018 | 0.005469575 |
| rs1490188 | 151985662 | 0.023147735 |
| rs2999539 | 151986627 | 0.007129832 |
| rs2932570 | 151986820 | 0.007129832 |
| rs1967085 | 151993065 | 0.023147735 |
| rs1967084 | 151993088 | 0.023147735 |
| rs2999548 | 151993314 | 0.023147735 |
| rs2932566 | 151993315 | 0.023147735 |
| rs115288876 | 152000117 | 0.025065752 |
| rs80298065 | 152016352 | 0.010480254 |
| rs75754657 | 152025632 | 0.004752172 |
| rs12130862 | 152027015 | 0.001960059 |
| rs55668963 | 152032745 | 0.001960059 |
| rs76408637 | 152038570 | 0.005553281 |
| rs2069258 | 152045626 | 0.002006003 |
| rs2999563 | 152046995 | 0.002997571 |
| rs116533286 | 152051550 | 0.005737945 |
| rs144964206 | 152052696 | 0.005553281 |
| rs376243217 | 152055232 | 0.005599475 |
| rs17646946 | 152062767 | 0.002124409 |
| rs76865868 | 152066130 | 0.004752172 |
| rs11334740 | 152068746 | 0.00853903 |
| rs1039088 | 152070131 | 0.004835624 |
| rs11803731 | 152083325 | 0.002346054 |
| rs36010924 | 152088844 | 0.002346054 |
| rs151256541 | 152097773 | 0.004835624 |
| rs12144907 | 152118217 | 0.002346054 |
| rs3001978 | 152126467 | 0.002867793 |
| rs145089986 | 152126933 | 0.00853903 |
| rs75957773 | 152127455 | 0.000369399 |
| rs903796 | 152144633 | 0.013316901 |
| rs12033248 | 152146648 | 0.021286534 |
| rs114394160 | 152152088 | 0.022550866 |
| rs1496042 | 152159773 | 0.031659373 |
| rs146355184 | 152162133 | 0.002497103 |
| **rs1496050** | **152163549** | **0.156574156** |

| | | |
|---|---|---|
| **rs1496051** | **152164923** | **0.156574156** |
| **rs1552991** | **152165878** | **0.062660718** |
| **rs10888466** | **152166223** | **0.156574156** |
| rs79717008 | 152167054 | 0.020802883 |
| **rs10788827** | **152168346** | **0.156574156** |
| **rs12117911** | **152168537** | **0.156574156** |
| **rs56283877** | **152168571** | **0.132865584** |
| **rs56407906** | **152168611** | **0.128232954** |
| **rs56210025** | **152168619** | **0.146687288** |
| **rs79853660** | **152169035** | **0.156574156** |
| **rs143634472** | **152169066** | **0.156574156** |
| **rs12047544** | **152169922** | **0.156574156** |
| **rs12133847** | **152170101** | **0.156574156** |
| **rs149885335** | **152170141** | **0.11115238** |
| **rs28682375** | **152170149** | **0.11115238** |
| **rs12144058** | **152170216** | **0.156574156** |
| rs115670257 | 152184907 | 0.022550866 |
| **rs4845426** | **152195359** | **0.162752703** |
| **rs7550106** | **152197202** | **0.162752703** |
| **rs1390488** | **152199500** | **0.162752703** |
| **rs10888471** | **152200849** | **0.162752703** |
| **rs11204939** | **152200942** | **0.162752703** |
| **rs4845750** | **152201217** | **0.162752703** |
| **rs145841925** | **152201899** | **0.162752703** |
| **rs35759221** | **152202787** | **0.157047566** |
| **rs11204941** | **152203343** | **0.162752703** |
| **rs11204943** | **152203906** | **0.162752703** |
| **rs10788830** | **152206898** | **0.162752703** |
| **rs11204945** | **152210698** | **0.162752703** |
| **rs71582223** | **152212083** | **0.162752703** |
| **rs1466757** | **152212469** | **0.162752703** |
| **rs1845791** | **152213772** | **0.162752703** |
| **rs2132418** | **152214783** | **0.162752703** |
| **rs10749670** | **152220467** | **0.162752703** |
| **rs12746538** | **152229296** | **0.168922796** |
| **rs138314100** | **152231760** | **0.157759196** |
| **rs4845753** | **152231834** | **0.168922796** |
| **rs10888476** | **152232321** | **0.162752703** |
| **rs4466659** | **152232566** | **0.162752703** |
| **rs4256811** | **152232733** | **0.162752703** |
| **rs4341364** | **152233713** | **0.162752703** |

| | | |
|---|---|---|
| **rs10888477** | **152234210** | **0.168922796** |
| **rs10888478** | **152235030** | **0.162752703** |
| **rs10788831** | **152235308** | **0.162752703** |
| **rs11204956** | **152236058** | **0.168922796** |
| **rs11204958** | **152236586** | **0.162752703** |
| **rs4528123** | **152239362** | **0.157384263** |
| **rs4394659** | **152239620** | **0.139844248** |
| **rs12730372** | **152240445** | **0.161575122** |
| **rs12747962** | **152240456** | **0.161575122** |
| **rs4595369** | **152248193** | **0.151640145** |
| **rs9887901** | **152248552** | **0.151640145** |
| **rs11204967** | **152249720** | **0.168922796** |
| **rs4634913** | **152249932** | **0.151640145** |
| **rs4534386** | **152251714** | **0.151640145** |
| **rs11485508** | **152253167** | **0.151640145** |
| **rs12065368** | **152253497** | **0.151640145** |
| **rs12027807** | **152253726** | **0.151640145** |
| **rs12045492** | **152253866** | **0.148281624** |
| **rs12027899** | **152253954** | **0.151640145** |
| **rs4845756** | **152254919** | **0.151640145** |
| **rs4845758** | **152255050** | **0.151640145** |
| **rs3126043** | **152256030** | **0.162752703** |
| **rs1858477** | **152265261** | **0.162752703** |
| **rs3126058** | **152269622** | **0.151640145** |
| rs144613541 | 152270875 | 0.049657763 |
| rs147652658 | 152270880 | 0.049657763 |
| **rs3126060** | **152271307** | **0.151640145** |
| **rs2184951** | **152273168** | **0.12775165** |
| **rs12730241** | **152274016** | **0.12775165** |
| **rs3126075** | **152276626** | **0.156314182** |
| **rs3126066** | **152276871** | **0.1421994** |
| **rs3091276** | **152277168** | **0.156594119** |
| **rs9436066** | **152277717** | **0.145992321** |
| **rs3126072** | **152279729** | **0.151640145** |
| **rs3126074** | **152279841** | **0.145992321** |
| **rs34224823** | **152287341** | **0.121044203** |
| **rs3120659** | **152293202** | **0.145992321** |
| **rs3120661** | **152294402** | **0.176136851** |
| **rs3126085** | **152300817** | **0.149556894** |
| **rs1858480** | **152301735** | **0.156594119** |
| **rs36142010** | **152302627** | **0.149556894** |

| | | |
|---|---|---|
| **rs35971937** | **152304644** | **0.149556894** |
| **rs3126088** | **152305785** | **0.149556894** |
| **rs1858482** | **152306921** | **0.149556894** |
| **rs3126091** | **152307338** | **0.176136851** |
| **rs3126092** | **152307955** | **0.149556894** |
| **rs3126094** | **152308243** | **0.156594119** |
| **rs2338555** | **152309380** | **0.149556894** |
| **rs2338556** | **152309427** | **0.149556894** |
| **rs3126097** | **152311026** | **0.176136851** |
| **rs3120664** | **152313206** | **0.14784571** |
| **rs6679449** | **152328411** | **0.166530558** |
| **rs10788835** | **152330945** | **0.166530558** |
| **rs2275264** | **152331533** | **0.166530558** |
| **rs11204984** | **152331987** | **0.166530558** |
| **rs35385577** | **152337325** | **0.119470668** |
| **rs4322234** | **152353768** | **0.157594698** |
| **rs6671146** | **152354472** | **0.149286768** |
| **rs11204994** | **152358640** | **0.149286768** |
| **rs1923506** | **152359700** | **0.157594698** |
| **rs1923503** | **152360820** | **0.157594698** |
| **rs7534951** | **152362972** | **0.149286768** |
| **rs10788838** | **152363201** | **0.121296247** |
| **rs7514696** | **152363874** | **0.149286768** |
| **rs11204998** | **152365655** | **0.149286768** |
| **rs2146119** | **152368898** | **0.157715011** |
| **rs7515448** | **152372035** | **0.151014629** |
| rs12048587 | 152378714 | 0.070733589 |
| rs1923491 | 152384945 | 0.070733589 |
| rs3753446 | 152388475 | 0.070733589 |
| rs549550056 | 152390743 | 0.068542538 |
| rs554599509 | 152390745 | 0.033493733 |
| rs12724738 | 152390751 | 0.027056549 |
| **rs113643562** | **152395654** | **0.127811092** |
| **rs11205001** | **152395948** | **0.127811092** |
| **rs726864** | **152400293** | **0.127811092** |
| rs11205002 | 152403339 | 0.058163219 |
| rs12749505 | 152404073 | 0.058163219 |
| rs113419595 | 152404275 | 0.058163219 |
| rs986903 | 152410569 | 0.058163219 |
| rs10888488 | 152411090 | 0.058163219 |
| rs6695950 | 152412687 | 0.058163219 |

| | | |
|---|---|---|
| rs4845433 | 152414439 | 0.058555705 |
| **rs111912262** | **152414878** | **0.101502831** |
| rs4845767 | 152415308 | 0.058555705 |
| rs6587673 | 152430152 | 0.044682486 |
| rs17597997 | 152432884 | 0.044682486 |
| rs57666146 | 152433876 | 0.044682486 |
| rs145572465 | 152436805 | 0.042408444 |
| rs12137893 | 152437397 | 0.044682486 |
| rs12134605 | 152437479 | 0.041568958 |
| rs12116760 | 152437883 | 0.044682486 |
| rs952180 | 152439945 | 0.041568958 |
| rs13373771 | 152443608 | 0.043190629 |
| rs34406913 | 152443670 | 0.043190629 |
| rs77033261 | 152444584 | 0.043190629 |
| rs12733078 | 152444658 | 0.040557191 |
| rs12757042 | 152444672 | 0.040557191 |
| rs61813862 | 152445162 | 0.043190629 |
| rs4845773 | 152445418 | 0.043190629 |
| rs4845439 | 152446495 | 0.043190629 |
| rs71582238 | 152447142 | 0.007523189 |
| rs12749879 | 152447991 | 0.051411672 |
| rs6698520 | 152448967 | 0.042403868 |
| rs4845777 | 152450239 | 0.042403868 |
| rs11205008 | 152450614 | 0.055293445 |
| rs4845440 | 152450898 | 0.055293445 |
| rs11578573 | 152452675 | 0.051411672 |
| **rs568184** | **152457223** | **0.142079557** |
| **rs1199153** | **152457624** | **0.142079557** |
| **rs1199154** | **152458435** | **0.142079557** |
| **rs1199157** | **152459811** | **0.142079557** |
| **rs1199160** | **152461668** | **0.142079557** |
| **rs1199161** | **152461813** | **0.142079557** |
| **rs145933516** | **152464741** | **0.142079557** |
| rs112791653 | 152467061 | 0.016617439 |
| **rs7552220** | **152479716** | **0.142079557** |
| rs6587679 | 152481262 | 0.00742897 |
| rs11205014 | 152483634 | 0.00742897 |
| rs4845780 | 152488846 | 0.00742897 |
| rs548252 | 152489742 | 0.00742897 |
| rs2181172 | 152491794 | 0.00742897 |

| | | |
|---|---|---|
| rs12132469 | 152491856 | 0.00742897 |
| rs4845782 | 152492225 | 0.00742897 |
| rs10888491 | 152492622 | 0.00742897 |
| rs11365304 | 152493337 | 0.042752493 |
| rs71582241 | 152504491 | 0.008694862 |
| rs10666767 | 152506982 | 0.008004378 |
| rs4845785 | 152510143 | 0.007797872 |
| rs908927 | 152513338 | 0.007797872 |
| rs12129646 | 152518816 | 0.005581745 |
| rs4845788 | 152519496 | 0.007797872 |
| rs138657579 | 152522870 | 0.007797872 |
| rs12034882 | 152525492 | 0.007797872 |
| rs4240883 | 152526260 | 0.007797872 |
| rs11205031 | 152527662 | 0.007797872 |
| rs11205032 | 152527690 | 0.007797872 |
| rs11205033 | 152527966 | 0.007797872 |
| rs4240885 | 152529850 | 0.007797872 |
| rs4845444 | 152530125 | 0.007797872 |
| rs11205035 | 152532832 | 0.007797872 |
| rs4112785 | 152533115 | 0.007797872 |
| rs4074613 | 152534191 | 0.006490981 |
| rs10888501 | 152537954 | 0.020440543 |
| rs7516108 | 152542229 | 0.001443546 |
| rs4085613 | 152550018 | 0.004058786 |
| rs477392 | 152555176 | 0.004058786 |
| esv3587563 | 152555495 | 0.004058786 |
| rs75567512 | 152587933 | 0.004058786 |
| rs61813888 | 152587979 | 0.004058786 |
| rs75196578 | 152588529 | 0.002405849 |
| rs78728337 | 152588771 | 0.004678449 |
| rs76930171 | 152588874 | 0.004678449 |
| rs11205042 | 152589643 | 0.004188768 |
| rs6677595 | 152590187 | 0.004058786 |
| rs6701307 | 152590444 | 0.004058786 |
| rs1886734 | 152591142 | 0.004058786 |
| rs4845450 | 152591200 | 0.004058786 |
| rs9633406 | 152591446 | 0.004058786 |
| rs4845453 | 152591953 | 0.004058786 |
| rs35655600 | 152599002 | 0.027687188 |
| rs7522480 | 152608876 | 0.050985645 |

| | | |
|---|---|---|
| rs10127684 | 152609072 | 0.060527953 |
| rs58237116 | 152609916 | 0.060527953 |
| rs16834273 | 152609951 | 0.060527953 |
| rs11205051 | 152610256 | 0.072167467 |
| rs11205053 | 152612045 | 0.060527953 |
| rs12030425 | 152612136 | 0.060527953 |
| rs1325508 | 152612910 | 0.064443085 |
| rs11205056 | 152613472 | 0.060527953 |
| rs12239456 | 152613593 | 0.060527953 |
| rs6671975 | 152613631 | 0.046982081 |
| rs142829848 | 152614880 | 0.060527953 |
| rs4353096 | 152616581 | 0.060527953 |
| rs4540662 | 152616737 | 0.060527953 |
| rs942827 | 152617423 | 0.060527953 |
| rs989834 | 152618544 | 0.061208458 |
| rs66587527 | 152619643 | 0.066092096 |
| rs12090832 | 152621249 | 0.060527953 |
| rs55952227 | 152621673 | 0.060527953 |
| **rs10888505** | **152625148** | **0.112907323** |
| **rs10888507** | **152626270** | **0.112907323** |
| **rs373465501** | **152626743** | **0.112907323** |
| **rs1332497** | **152632433** | **0.112907323** |
| **rs11205062** | **152635201** | **0.112907323** |
| **rs11205063** | **152635450** | **0.112907323** |
| rs11205065 | 152643998 | 0.055830444 |
| rs1930126 | 152647579 | 0.055998106 |
| rs1853805 | 152649114 | 0.051992108 |
| **rs1412547** | **152649574** | **0.112907323** |
| **rs1412546** | **152649645** | **0.112907323** |
| rs2014369 | 152651023 | 0.051992108 |
| rs2014368 | 152651029 | 0.051992108 |
| rs7546815 | 152651851 | 0.051992108 |
| rs12097462 | 152652137 | 0.051992108 |
| rs10788844 | 152653325 | 0.051992108 |
| rs1925663 | 152653506 | 0.051992108 |
| rs11205073 | 152654385 | 0.051992108 |
| rs1332507 | 152660493 | 0.051992108 |
| rs1332508 | 152660503 | 0.051992108 |
| rs1332509 | 152660746 | 0.051992108 |
| rs4845461 | 152661738 | 0.050808946 |

| | | |
|---|---|---|
| rs7522449 | 152662863 | 0.051992108 |
| rs7525190 | 152663352 | 0.051992108 |
| rs10465792 | 152664975 | 0.051992108 |
| rs10465797 | 152665640 | 0.051992108 |
| rs10788845 | 152668072 | 0.051992108 |
| rs3904413 | 152668498 | 0.051992108 |
| rs11205078 | 152669459 | 0.051992108 |
| rs11581506 | 152673259 | 0.051992108 |
| rs11587581 | 152674066 | 0.051992108 |
| rs1412542 | 152678445 | 0.051992108 |
| rs1536157 | 152686773 | 0.046992327 |
| rs4845466 | 152690745 | 0.029466529 |
| rs4845467 | 152690825 | 0.029466529 |
| rs4845468 | 152691078 | 0.029466529 |
| rs10888511 | 152697881 | 0.046992327 |
| rs55683499 | 152701156 | 0.010591871 |
| rs61812714 | 152708298 | 0.010772426 |
| rs11205094 | 152708619 | 0.048402414 |
| rs4845473 | 152709456 | 0.010772426 |
| rs4845474 | 152709548 | 0.010772426 |
| rs66659820 | 152719538 | 0.05756506 |
| rs28464852 | 152721454 | 0.051870095 |
| rs9729955 | 152721468 | 0.051870095 |
| rs6421466 | 152722707 | 0.056525161 |
| rs1412550 | 152723533 | 0.044310677 |
| rs1332504 | 152723672 | 0.044310677 |
| rs1332506 | 152724053 | 0.044310677 |
| rs35393920 | 152726908 | 0.009040456 |
| rs57629863 | 152727709 | 0.007439475 |
| rs56180022 | 152727768 | 0.009040456 |
| rs11578927 | 152728472 | 0.009040456 |
| rs1412548 | 152730656 | 0.007439475 |
| rs12725760 | 152735171 | 0.009040456 |
| rs35274012 | 152735687 | 0.009040456 |
| rs4539140 | 152736291 | 0.009040456 |
| rs6680648 | 152737228 | 0.055897885 |
| rs6587702 | 152737703 | 0.055897885 |
| rs10494277 | 152739226 | 0.010264643 |
| rs4845482 | 152746534 | 0.009040456 |
| rs12239774 | 152748253 | 0.009040456 |

| | | |
|---|---|---|
| rs913997 | 152750256 | 0.04647053 |
| rs11205103 | 152754916 | 0.052400806 |
| rs4845483 | 152755506 | 0.081309925 |
| rs950338 | 152756806 | 0.05756506 |
| rs7530000 | 152757690 | 0.05756506 |
| rs11576947 | 152758519 | 0.048893923 |
| rs10888516 | 152759010 | 0.050854831 |
| rs1034109 | 152759678 | 0.056327303 |
| rs201660535 | 152760075 | 0.058874425 |
| rs112600788 | 152760093 | 0.040574688 |
| rs114470093 | 152760558 | 0.019563669 |
| rs150561698 | 152760642 | 0.06614123 |
| rs16834535 | 152761091 | 0.065076414 |
| rs1831639 | 152761454 | 0.065076414 |
| rs6587704 | 152761709 | 0.072448261 |
| rs200638328 | 152763556 | 0.055481008 |
| rs4845485 | 152766239 | 0.065076414 |
| rs1537307 | 152769807 | 0.07224181 |
| rs41268492 | 152770521 | 0.042145927 |
| rs6587705 | 152774367 | 0.053233409 |
| rs7536191 | 152783560 | 0.065710472 |
| rs12023196 | 152783724 | 0.065710472 |
| **rs11804609** | **152784255** | **0.157320812** |
| **rs2339399** | **152788161** | **0.182071645** |
| **rs7550676** | **152793205** | **0.182071645** |
| **rs35160213** | **152793787** | **0.182071645** |
| **rs6659798** | **152795971** | **0.182071645** |
| **rs12565568** | **152823380** | **0.182071645** |
| **rs2339396** | **152842300** | **0.182071645** |
| **rs2339397** | **152842930** | **0.182071645** |
| **rs12022319** | **152847892** | **0.172370691** |
| **rs4845490** | **152849299** | **0.172370691** |
| **rs4845491** | **152849569** | **0.13878888** |
| **rs3737861** | **152850938** | **0.094030563** |
| **rs16834728** | **152853137** | **0.126226556** |
| **rs4845492** | **152853402** | **0.126226556** |
| rs6587710 | 152870716 | 0.027588312 |
| **rs4845327** | **152879512** | **0.097803393** |
| **rs1854779** | **152880672** | **0.097803393** |
| **rs7539232** | **152881802** | **0.097803393** |
| **rs11205132** | **152882135** | **0.097803393** |

| | | |
|---|---|---|
| **rs2229496** | **152882610** | **0.097803393** |
| **rs7520711** | **152882982** | **0.074447868** |
| **rs7535306** | **152883680** | **0.097803393** |
| **rs7545520** | **152883711** | **0.097803393** |
| **rs3820136** | **152884438** | **0.089076765** |
| rs4845496 | 152888178 | 0.028530482 |
| **rs3845340** | **152889661** | **0.07189708** |
| rs56335979 | 152891350 | 0.035843277 |
| rs11587269 | 152891805 | 0.035128149 |
| rs2339386 | 152892307 | 0.035128149 |
| rs7541328 | 152894142 | 0.035128149 |
| rs6663448 | 152894627 | 0.035128149 |
| rs1999885 | 152907785 | 0.057839036 |
| rs377142541 | 152912278 | 0.034684475 |
| rs2066005 | 152923113 | 0.038129675 |
| rs7556517 | 152925755 | 0.038129675 |
| rs61815715 | 152927706 | 0.032603274 |
| rs28608883 | 152929945 | 0.032603274 |
| rs6690549 | 152930426 | 0.038129675 |
| rs6587713 | 152932031 | 0.037814056 |
| rs56347903 | 152932175 | 0.032450633 |
| rs12748969 | 152932654 | 0.032450633 |
| rs4845500 | 152933011 | 0.037814056 |
| rs6587714 | 152933023 | 0.039617527 |
| rs6587715 | 152933960 | 0.041830426 |
| rs139250679 | 152936484 | 0.015102958 |
| rs6675352 | 152936749 | 0.039617527 |
| rs4845501 | 152937401 | 0.032450633 |
| rs1890284 | 152942190 | 0.036174852 |
| rs3170863 | 152944594 | 0.036174852 |
| rs2879490 | 152947163 | 0.036174852 |
| rs4240865 | 152954839 | 0.043156618 |
| rs1415964 | 152955085 | 0.043156618 |
| rs1611760 | 152957573 | 0.032409384 |
| rs1129654 | 152958238 | 0.032409384 |
| rs1129655 | 152958267 | 0.032409384 |
| rs1984198 | 152958584 | 0.032409384 |
| rs1415962 | 152958731 | 0.032409384 |
| rs1577964 | 152961784 | 0.034873353 |
| rs1577962 | 152962013 | 0.034873353 |

| | | |
|---|---|---|
| rs1933383 | 152963468 | 0.034873353 |
| rs10788850 | 152963932 | 0.034873353 |
| rs73014331 | 152964533 | 0.034873353 |
| rs6668311 | 152964586 | 0.034873353 |
| rs6587716 | 152966705 | 0.034873353 |
| rs6665527 | 152967009 | 0.034873353 |
| rs61811427 | 152968419 | 0.032745834 |
| rs6667418 | 152969685 | 0.039054139 |
| rs1415961 | 152973464 | 0.031614154 |
| rs1415960 | 152973571 | 0.028896629 |
| rs1415959 | 152973690 | 0.028896629 |
| rs3753454 | 152973776 | 0.031614154 |
| rs6671414 | 152974119 | 0.028896629 |
| rs946097 | 152974732 | 0.031614154 |
| rs946098 | 152975038 | 0.031614154 |
| rs10399896 | 152975083 | 0.031313434 |
| rs2339494 | 152975143 | 0.031313434 |
| rs1055935 | 152975941 | 0.031614154 |
| rs3737867 | 152976474 | 0.031614154 |
| rs6673691 | 152978289 | 0.031614154 |
| rs2339486 | 152983513 | 0.030294476 |
| rs11205163 | 152983677 | 0.031614154 |
| rs2339490 | 152984612 | 0.031614154 |
| rs11205165 | 152986928 | 0.029094808 |
| rs1591736 | 152987497 | 0.029094808 |
| rs9659389 | 152989844 | 0.029094808 |
| rs6685731 | 152990203 | 0.029094808 |
| rs2152994 | 152991060 | 0.029094808 |
| rs11205172 | 152992408 | 0.029094808 |
| rs17881363 | 153002334 | 0.030745734 |
| rs4845336 | 153003853 | 0.026078812 |
| rs4845516 | 153003892 | 0.026078812 |
| rs4845517 | 153003893 | 0.026078812 |
| rs4845518 | 153003903 | 0.028781481 |
| rs4845519 | 153003916 | 0.026078812 |
| rs1108410 | 153005662 | 0.032177057 |
| rs2937263 | 153006756 | 0.033876205 |
| rs3120744 | 153010330 | 0.025886112 |
| rs72704879 | 153012991 | 0.041824703 |
| rs310124 | 153020275 | 0.035257224 |

| | | |
|---|---|---|
| rs72704888 | 153020864 | 0.008688516 |
| rs6698361 | 153023184 | 0.028108044 |
| rs12082627 | 153023650 | 0.028108044 |
| rs12079087 | 153023888 | 0.028108044 |
| rs7551791 | 153024267 | 0.03107173 |
| rs11586559 | 153025714 | 0.03107173 |
| rs56180170 | 153026598 | 0.03107173 |
| rs6661059 | 153027257 | 0.028108044 |
| rs3737864 | 153029137 | 0.024770994 |
| rs6664380 | 153030365 | 0.028073277 |
| rs454245 | 153031979 | 0.024268286 |
| rs12083211 | 153032131 | 0.028073277 |
| rs582345 | 153033406 | 0.024268286 |
| rs6675009 | 153033783 | 0.031036862 |
| rs6587724 | 153034501 | 0.028073277 |
| rs608509 | 153034650 | 0.024268286 |
| rs61811886 | 153035743 | 0.031036862 |
| rs821755 | 153036245 | 0.024268286 |
| rs140372917 | 153040091 | 0.095834268 |
| rs576941 | 153040165 | 0.024268286 |
| rs71918470 | 153042468 | 0.095834268 |
| rs310103 | 153042554 | 0.030019627 |
| rs426147 | 153045011 | 0.024268286 |
| rs689008 | 153047100 | 0.024268286 |
| rs662506 | 153048386 | 0.024268286 |
| rs454186 | 153049149 | 0.024268286 |
| rs431242 | 153050414 | 0.030019627 |
| rs140957528 | 153051683 | 0.095834268 |
| rs148418088 | 153052414 | 0.009007131 |
| rs58755826 | 153057913 | 0.081539777 |
| rs11581933 | 153069407 | 0.041730665 |
| rs61811907 | 153070024 | 0.007326007 |
| rs61811908 | 153070662 | 0.041730665 |
| rs635505 | 153070981 | 0.037138429 |
| rs6679881 | 153073420 | 0.041730665 |
| rs11587454 | 153075171 | 0.041730665 |
| rs6704131 | 153077102 | 0.041730665 |
| rs11582619 | 153080966 | 0.041730665 |
| rs11584009 | 153081204 | 0.041730665 |
| rs540645152 | 153082377 | 0.041730665 |

| | | |
|---|---|---|
| rs28846828 | 153082810 | 0.041730665 |
| rs140524342 | 153083362 | 0.039828477 |
| rs2133961 | 153084795 | 0.048072837 |
| rs34115170 | 153085306 | 0.048823634 |
| rs1392806 | 153089634 | 0.046567642 |
| rs11586440 | 153091081 | 0.048823634 |
| rs6679004 | 153092124 | 0.046567642 |
| rs310133 | 153096423 | 0.038858688 |
| rs1119266 | 153097240 | 0.046567642 |
| rs1604264 | 153105819 | 0.046567642 |
| rs35225843 | 153106637 | 0.046567642 |
| rs12752746 | 153107314 | 0.046567642 |
| rs34863954 | 153115963 | 0.046567642 |
| rs77118675 | 153116692 | 0.046567642 |
| rs12734959 | 153117671 | 0.048823634 |
| rs11577955 | 153119519 | 0.04814242 |
| rs533437 | 153119969 | 0.039969588 |
| rs12744123 | 153121135 | 0.039969588 |
| rs12724705 | 153121274 | 0.043755163 |
| rs510277 | 153122310 | 0.039969588 |
| rs516280 | 153122930 | 0.039969588 |
| rs11578865 | 153126302 | 0.04814242 |
| rs476761 | 153134300 | 0.039969588 |
| rs475698 | 153134467 | 0.038124447 |
| rs473096 | 153134693 | 0.038124447 |
| rs77668680 | 153135100 | 0.038854242 |
| rs503914 | 153135119 | 0.039969588 |
| rs556824 | 153135361 | 0.038124447 |
| rs555926 | 153135459 | 0.038124447 |
| rs553175 | 153135780 | 0.038124447 |
| rs474200 | 153136053 | 0.039595543 |
| rs550463 | 153136056 | 0.037751824 |
| rs531699 | 153137229 | 0.038124447 |
| rs491959 | 153137806 | 0.038124447 |
| rs547119 | 153139236 | 0.043755163 |
| rs138114182 | 153139960 | 0.041375956 |
| rs9970592 | 153140118 | 0.040496684 |
| rs11585767 | 153142328 | 0.04814242 |
| rs722660 | 153168883 | 0.00377418 |
| rs1410868 | 153169338 | 0.003289111 |

| | | |
|---|---|---|
| rs1831238 | 153169522 | 0.003289111 |
| rs1410869 | 153169596 | 0.003289111 |
| rs1952465 | 153170276 | 0.003289111 |
| rs11205217 | 153170938 | 0.003289111 |
| rs10888535 | 153171148 | 0.003289111 |
| rs12145115 | 153179923 | 0.003414978 |
| rs12116563 | 153181362 | 0.002582042 |
| rs11205219 | 153181916 | 0.002171602 |
| rs1410858 | 153182116 | 0.002950512 |
| rs1980909 | 153183327 | 0.003255734 |
| rs6587731 | 153187680 | 0.00171837 |
| rs10788862 | 153191512 | 0.001599552 |
| rs11205232 | 153192119 | 0.00247113 |
| rs10494295 | 153195590 | 0.001599552 |
| rs71626758 | 153197738 | 0.001599552 |
| rs11205236 | 153200272 | 0.00306489 |
| rs140786948 | 153205310 | 0.00399169 |
| rs16835086 | 153206051 | 0.004947687 |
| rs4517302 | 153207416 | 0.004947687 |
| rs6696243 | 153208091 | 0.004947687 |
| rs6587732 | 153208387 | 0.004947687 |
| rs34846977 | 153208777 | 0.00306489 |
| rs12135864 | 153213098 | 0.001497096 |
| rs12126039 | 153214352 | 0.001552853 |
| rs111162279 | 153216195 | 0.001763792 |
| rs75437404 | 153216817 | 0.003139244 |
| rs12565904 | 153217580 | 0.003404511 |
| rs7522467 | 153219328 | 0.002368165 |
| rs12131158 | 153221315 | 0.002368165 |
| rs34961571 | 153221438 | 0.004666197 |
| rs150552072 | 153221488 | 0.002355575 |
| rs7554266 | 153222881 | 0.002204115 |
| rs7518709 | 153225883 | 0.002204115 |
| rs7518712 | 153225886 | 0.002204115 |
| rs6666892 | 153226579 | 0.002204115 |
| rs873234 | 153227177 | 0.003212798 |
| rs2094638 | 153227608 | 0.003067047 |
| rs75887175 | 153228359 | 0.002721681 |
| rs36063924 | 153228880 | 0.004090709 |
| rs12727071 | 153229447 | 0.003416321 |

| | | |
|---|---|---|
| rs11287299 | 153229673 | 0.003743779 |
| rs10888545 | 153229682 | 0.002371553 |
| rs10567874 | 153230481 | 0.003329244 |
| rs11205254 | 153231483 | 0.002719833 |
| rs6661601 | 153233510 | 0.002846301 |
| rs12138538 | 153235837 | 0.002128009 |
| rs4559442 | 153236224 | 0.003067047 |
| rs4319268 | 153243767 | 0.002409254 |
| rs10888548 | 153245625 | 0.003857515 |
| rs529510353 | 153255533 | 0.002007649 |
| rs3886614 | 153257106 | 0.001282372 |
| rs3856024 | 153258816 | 0.001282372 |
| rs67401364 | 153265396 | 0.027131689 |
| rs11585177 | 153266256 | 0.001282372 |
| rs61803281 | 153267873 | 0.000937695 |
| rs2771110 | 153270362 | 0.002679225 |
| rs1433679 | 153271048 | 0.002101599 |
| rs821430 | 153273259 | 0.002606657 |
| rs821429 | 153275443 | 0.004747341 |
| rs821427 | 153275977 | 0.004248812 |
| rs2771112 | 153277660 | 0.002208995 |
| rs2570443 | 153277768 | 0.002208995 |
| rs3006445 | 153280190 | 0.001186663 |
| rs3006447 | 153280303 | 0.001186663 |
| rs9970937 | 153280387 | 0.001186663 |
| rs527287616 | 153280476 | 0.001186663 |
| rs113944017 | 153285078 | 0.000568188 |
| rs3006424 | 153289558 | 0.000877522 |
| rs3006425 | 153289576 | 0.000877522 |
| rs1094362 | 153290524 | 0.002192292 |
| rs821414 | 153292639 | 0.002194431 |
| rs821411 | 153294077 | 0.001735986 |
| rs821408 | 153296027 | 0.001735986 |
| rs821407 | 153297131 | 0.002174857 |
| rs1836118 | 153298525 | 0.002192292 |
| rs2771115 | 153299965 | 0.001735986 |
| rs2771116 | 153300351 | 0.002192292 |
| rs2771118 | 153301527 | 0.001735986 |
| rs2771120 | 153302417 | 0.001570682 |
| rs1655310 | 153303930 | 0.002192292 |

| | | |
|---|---|---|
| rs1754132 | 153304768 | 0.002194431 |
| rs1655308 | 153304769 | 0.002194431 |
| rs151252426 | 153308203 | 0.009484832 |
| rs1754134 | 153310747 | 0.002194431 |
| rs3006449 | 153314347 | 0.009592667 |
| rs3006450 | 153314534 | 0.009592667 |
| rs3014861 | 153314822 | 0.009592667 |
| rs3014862 | 153314999 | 0.009592667 |
| rs735012 | 153316260 | 0.009592667 |
| rs3006452 | 153317370 | 0.009592667 |
| rs2916210 | 153319380 | 0.007174889 |
| rs6587737 | 153319811 | 0.007174889 |
| rs2916208 | 153320125 | 0.009592667 |
| rs2916205 | 153320755 | 0.009592667 |
| rs3806231 | 153321348 | 0.009592667 |
| rs2916202 | 153321810 | 0.009592667 |
| rs12070252 | 153323280 | 0.011550558 |
| rs72708726 | 153323574 | 0.011550558 |
| rs72708730 | 153323620 | 0.011550558 |
| rs72708731 | 153323734 | 0.011550558 |
| rs4845345 | 153324212 | 0.017037602 |
| rs12561811 | 153324548 | 0.018610259 |
| rs35364949 | 153324802 | 0.018610259 |
| rs10888558 | 153325130 | 0.018610259 |
| rs1347250 | 153325954 | 0.026595196 |
| rs11580993 | 153339314 | 0.01336903 |
| rs60752752 | 153339782 | 0.004330823 |
| rs58644524 | 153342575 | 0.004330823 |
| rs35195593 | 153343717 | 0.004330823 |
| rs3006475 | 153344636 | 0.004330823 |
| rs2916191 | 153345911 | 0.004330823 |
| rs4772 | 153346263 | 0.004330823 |
| rs141832834 | 153350207 | 0.004330823 |
| rs3014885 | 153352127 | 0.004330823 |
| rs374815343 | 153352260 | 0.004330823 |
| rs2916221 | 153352865 | 0.004330823 |
| rs2916217 | 153353198 | 0.004330823 |
| rs3006479 | 153355594 | 0.004330823 |
| rs3006482 | 153356497 | 0.004330823 |
| rs3006488 | 153362507 | 0.005215585 |

| | | |
|---|---|---|
| rs113420049 | 153374406 | 0.00560992 |
| rs112955890 | 153375455 | 0.00560992 |
| rs6673905 | 153376145 | 0.00560992 |
| rs6686657 | 153376207 | 0.00560992 |
| rs11205283 | 153376350 | 0.00560992 |
| rs11205286 | 153376743 | 0.00560992 |
| rs11580424 | 153377319 | 0.00560992 |
| rs74436639 | 153382458 | 0.00560992 |
| rs11590633 | 153382941 | 0.00560992 |
| rs3014812 | 153388881 | 0.0082072 |
| rs3006414 | 153391729 | 0.019924119 |
| rs3014822 | 153400479 | 0.019924119 |
| rs3006422 | 153400672 | 0.019924119 |
| rs3006423 | 153401664 | 0.019924119 |
| rs2026604 | 153408831 | 0.007208074 |
| rs10888561 | 153410822 | 0.007015693 |
| rs4845349 | 153411867 | 0.007208074 |
| rs10888562 | 153418785 | 0.007208074 |
| rs6587740 | 153419033 | 0.007208074 |
| rs4845545 | 153422325 | 0.007208074 |
| rs6700785 | 153425008 | 0.008097887 |
| rs4418538 | 153426161 | 0.007208074 |
| rs715754 | 153428325 | 0.007208074 |
| rs12759195 | 153437423 | 0.007379039 |
| rs10888563 | 153445024 | 0.008674239 |
| rs10888564 | 153445100 | 0.009220495 |
| rs2986213 | 153445736 | 0.008083421 |
| rs5777871 | 153449063 | 0.008974717 |
| rs2038928 | 153449174 | 0.008974717 |
| rs2038927 | 153449230 | 0.008974717 |
| rs12145434 | 153449498 | 0.008974717 |
| rs71584111 | 153450133 | 0.007525755 |
| rs10888566 | 153450379 | 0.008974717 |
| rs4845547 | 153450676 | 0.008974717 |
| rs1555885 | 153452831 | 0.008974717 |
| rs6657653 | 153454619 | 0.009966907 |
| rs4590630 | 153454674 | 0.009966907 |
| rs11205301 | 153456281 | 0.009966907 |
| rs913057 | 153457552 | 0.009966907 |
| rs7535089 | 153459023 | 0.014389334 |

| | | |
|---|---|---|
| rs7512616 | 153459971 | 0.009966907 |
| rs71584114 | 153460118 | 0.007375957 |
| rs6421468 | 153463764 | 0.012662695 |
| rs28410699 | 153482272 | 0.009691137 |
| rs6587745 | 153485284 | 0.009691137 |
| rs6587746 | 153485359 | 0.009691137 |
| rs16835393 | 153487225 | 0.009691137 |
| rs7544359 | 153487714 | 0.009691137 |
| rs7515940 | 153488027 | 0.009691137 |
| rs7537898 | 153488080 | 0.009691137 |
| rs7534823 | 153490810 | 0.009691137 |
| rs4845554 | 153492017 | 0.009691137 |
| rs7518306 | 153496247 | 0.009691137 |
| rs113973768 | 153497200 | 0.005229029 |
| rs9427239 | 153501202 | 0.017365632 |
| rs9426945 | 153502121 | 0.017365632 |
| rs71584119 | 153503281 | 0.011448668 |
| rs7535476 | 153514241 | 0.028146185 |
| rs1810765 | 153515120 | 0.035920621 |
| rs11488669 | 153517519 | 0.028744228 |
| rs56043239 | 153518483 | 0.028744228 |
| rs56089062 | 153520400 | 0.028744228 |
| rs28718953 | 153520784 | 0.018402856 |
| rs28594230 | 153520790 | 0.018402856 |
| rs28548347 | 153521018 | 0.018402856 |
| rs59376279 | 153521321 | 0.018402856 |
| rs57024596 | 153521496 | 0.018402856 |
| rs61265945 | 153521525 | 0.018402856 |
| rs4595398 | 153521653 | 0.018402856 |
| rs75928889 | 153521952 | 0.003546945 |
| rs3762268 | 153522350 | 0.022376662 |
| rs6427301 | 153524349 | 0.022376662 |
| rs74115476 | 153525417 | 0.022376662 |
| rs55955983 | 153526349 | 0.022376662 |
| rs8401 | 153533898 | 0.022376662 |
| rs6671398 | 153535509 | 0.022376662 |
| rs6703014 | 153540320 | 0.040423067 |
| rs144816475 | 153553932 | 0.004828032 |
| rs9658815 | 153555057 | 0.003268021 |
| rs138631403 | 153556248 | 0.005002505 |

| | | |
|---|---|---|
| rs28857273 | 153560040 | 0.005623494 |
| rs113117543 | 153563742 | 0.003268021 |
| rs56379573 | 153564661 | 0.005002505 |
| rs2026564 | 153565636 | 0.005623494 |
| rs555939948 | 153567927 | 0.00530615 |
| rs9729962 | 153568986 | 0.004441345 |
| rs73009940 | 153569599 | 0.005623494 |
| rs28737329 | 153571023 | 0.005623494 |
| rs139734691 | 153572182 | 0.005002505 |
| rs148716453 | 153573725 | 0.004574599 |
| rs143213672 | 153574099 | 0.001404284 |
| rs372416236 | 153574152 | 0.005002505 |
| rs566716273 | 153575424 | 0.00294072 |
