## SupplementalTable4 for "An enhancer:involucrin regulatory module impacts human skin barrier adaptation out-of-Africa and modifies atopic dermatitis risk"

**Table S4. CRISPR/Cas9 editing strategy reagents and mouse allele sequences.**

**a. Small Guide RNA (sgRNA) targeting sequences**

| | |
|---|---|
| Upstream 5' sgRNA | 5' - GAATACATCCCAGGAACAT -3' |
| Downstream 3' sgRNA | 5' - CAGTAAGCTAGCGCTAGAC -3' |

**b. ssODN sequences:** SphI restriction site, LoxP site, HindIII restriction site, **Homology arms**

| | |
|---|---|
| Upstream 5' ssODN | **5'- AGAAGTTTTTCAGTTCCCCATAGTTGTCCTGAGGAGCATATAATCTTTGTCTTAAGCAGATTTGTTTACAATAATTCCCT**ATAACTTCGTATAGCATACATTATACGAAGTTAT GCATGC**TTTAAAGAGATAGAGGACTGACATGACCCTCTGTCCTCTAAAACAAGTTTGCCAGGATTTCTCCATTCCCAGAGCCATGA**-3' |
| Downstream 3'ssODN | 5' - **TCTCTGTTGTTAGAGTCCATCTCCTACACCGATAGAGACTGATTCTGAAAAAAAAGGAAGCTCCCACTGTCCAAGTTCTA**AAGCTTATAACTTCGTATAGCATACATTATACGAAGTTATT**GGAAACCAGACACCCTGGCTGCTGCTCTGAAGGCAACTCTTCCCTATCAGGCTCCTTAATAGGATTTGATCAGTGTGAC**-3' |

**c. 923 WT**

TCTTTAGTGCTCAGTTAACAGCTTATTTTATGGAGTTCATCATTAACACTTTTTTATGAGA
TCATACAAAATAATATAGTAAAATAATGGAAAGATAAAACTCATTTCTAATTAGTCTTG **AG**
**AAGTTTTTCAGTTCCCCATAGTTGTCCTGAGGAGCATATAATCTTTGTCTTAAGCAGATTTGTTTA**
**CAATAATTCCCT**ATGTTCCTGGGATGTATTC**TTTAAAGAGATAGAGGACTGACATGACCCTCTGT**
**CCTCTAAAACAAGTTTGCCAGGATTTCTCCATTCCCAGAGCCATGA**GGCATCCTGAACACTACT
CTGAACTATATTTCTTTCTTCCTTTCTTTCTTCCTTCCTTCCTTCCTTCCTTCCTTCCTTCCT
TCCTTCCTTCCTTCCTTCCTTTCTTCCTTCCTTCTTTTCTTTTTCTTTCTTTCTTTCTTTCTTTCTTT
CTTTCTTTCTTTCTTTTTCTCTTTTTCTTTCCTTTTTTTGCTTTTATTTATTTTTTTTTCTTTTTCTATT
AGATATTTTCTTCATTTACATTTCAAATGTTATCCCCAAGCCCCCTATACCCTCCCCCAGTCCTGA
TCCCCAACCCACCCACACCCACTTCCTGGCCCTGGCACTACTTTCTATACCAAAGAAAGCATTC
CCCACCCCACCAGAAGAAGTAAGCAAGCATTCTCACATGAGCACTTATGCTCCACTTCTGACTT
CACATGGGAAGAATCTGACTCTCCTCAACCTGTGACAGTGCCAGGGCAGCAGACTGGTCAAAAA
GTCACACTGGTCTTATGGGTTCCCAGAGGCTCAGTATCTGCTCAATCTGTTTCCCACCAGCTGA
TTCAGAGTATGATAAGAATTCAGAAATGATACTGTGTGTGCGTGAGTGTTTGAGCACTGGGAAAA
GCTAAGGTGTGGGAATGAGGGCATAGGATAGAGCCCAGAAACCTGTGTGTAGTTGAAGGAGGG
GTTGAAGAAGCTCCAGACTTCTAATGCTCAAAGGTCACATATTTTGCCCTAGGATTATCCCACTT
AGCGACTGGGAATGCATGTCAATTTGGCATTTTTTTTTCAGTGTGCTGTGTGACTGACTTTATAA
GTCTCAGATCCTTGTGATGAATCCAAGAAACTATGCAATGCAAATTATACAAATTCTCCAGTGTA
ATGAAGGTAACCTTTCCCATAACCCATGAAGAGGCCTTGACCCAGCTCGGCCTCAGTGTTTAGG
AGGATAAGAGAAGGTGAAGGGATGAATATGACCAGAATGTGTGAAATTGGCAGAGAATGAATTA
TTTCTGAAAACTTGCCTTTGAAGAGTTTAGAGTGCTGCAGCTTCTTCAGAGAACATCA **TCTCTGT**
**TGTTAGAGTCCATCTCCTACACCGATAGAGACTGATTCTGAAAAAAAAGGAAGCTCCCACTGTC**
**CAAGTTCTA**CAGTAAGCTAGCGCTAGAC**TGGAAACCAGACACCCTGGCTGCTGCTCTGAAGGC**
**AACTCTTCCCTATCAGGCTCCTTAATAGGATTTGATCAGTGTGAC**AGGTTTCACTACATGACTAC
AGAGACATCCTCTAAGTCCAATAAGTTCCTGTGAGAATTTGGTGAGGCA

**d. 923del**

TCTTTAGTGCTCAGTTAACAGCTTATTTTATGGAGTTCATCATTAACACTTTTTTATGAGA
TCATACAAAATAATATAGTAAAATAATGGAAAGATAAAACTCATTTCTAATTAGTCTTG **AG**
**AAGTTTTTCAGTTCCCCATAGTTGTCCTGAGGAGCATATAATCTTTGTCTTAAGCAGATT**
**TGTTTACAATAATTCCC**ACTGTCCACTAAAGCTTATAACTTCGTATAGCATACATTATAC
GAAGTTAT**TGGAAACCAGACACCCTGGCTGCTGCTCTGAAGGCAACTCTTCCCTATCA**
**GGCTCCTTAATAGGATTTGATCAGTGTGAC**AGGTTTCACTACATGACTACAGAGACATC
CTCTAAGTCCAATAAGTTCCTGTGAGAATTTGGTGAGGCAA

**e. 923large**

TCCTCTGAATGCCCTAACTATCAGATTGTNTTCAGCTTTAATTAACTAATAAATTTTAGTT
ATTCTATCTATATTTATTTCATATTATTTATCTGTCTTCCACTGAAAAACAAGTTATATTTT
GAGAGAAATATTCTGGGTGTGCTTTCCATTGTCTCAAGGACCTATCAAAGTCACTCCAT
ACACTAAAACACTATCAGTATTAATTAAAAGAATAAATGACAGCAAATCTCATACCTACA
GACAACAACTTCCTAATATTTTAATGTCAAACAATCTTCATGTGTTTGAAAATGTGTGTG
CTAGGAAAATAAAGCTGAATTGTGGCTTATTTTTGTCTTTAGTGCTCAGTTAACAGCTTA
TTTTATGGAGTTCATCATTAACACTTTTTTATGAGATCATACAAAATAATATAGTAAAATA
ATGGAAAGATAAAACTCATTTCTAATTAGTCTTG **AGAAGTTTTTCAATTTCCCATAGTTGT**
**CCTGAGGAGCATATAATCTTTGTCTTAAGCAGATTTGTTTACAATAATTCCC**ATAACTTC
GTATAGCATACATTATACGAAGTTAT GCATGC**TTTAAAGAGATAGAGGACTGA**TCATGG
ACTTGGTAAATAGCCATATAAAATAGGAGCAGGTGGAAAAAAACATTTTCATTT **CTGATT**
**CTGAAAAAAAAGGAAGCTCCCACTGTCCAAGTTCTA**AAGCTTATAACTTCGTATAGCAT
ACATTATACGAAGTTAT**TGGAAACCAGACACCCTGGCTGCTGCTCTGAAGGCAACTCTT**
**CCCTATCAGGCTCCT**CAATAGGTGTTCTACATGAATGTATTGCTATGAAGCTACAGAGA
ACTGAAATACAAATTCCCAGAAATCTGTCCCTGAGAGGAGAAGAACCCACTTGAGGGTC
CTCTGCACTTCTGATCAGGGTCTCAAGAACTCACAGAAATCACAGTTATGCACCATGAT
CAATTTTATTGTTGTTGAACTGAAAGGTAGGCTAAAGAAAGAAACAAGAAATGTTTTTCT
AGCCAAGAGAGGTGGAGGG

**f. Genotyping primers**

| Mouse Allele | F Primer | R primer | Length |
|---|---|---|---|
| 923 WT | TCTTTAGTGCTCAGTTAACAGCT | AGAGTAGTGTTCAGGATG[illegible] | 319 |
| 923del | CAGTTCCCCATAGTTGTCCTG | TGCCTCACCAAATTCTCA[illegible] | 253 |
| 923large | CAGTTCCCCATAGTTGTCCTG*** | GGAAGAGTTGCCTTCAGA[illegible] | 317 |

***Primer doesn’t exactly match sanger sequence..(CAATTTCCCATAGTTGTCCTG)
