## SupplementalTable5 for "An enhancer:involucrin regulatory module impacts human skin barrier adaptation out-of-Africa and modifies atopic dermatitis risk"

**Table S5. Offspring genotype distribution from heterozygous 923del as well as 923large parental crosses.**

923del Het Intercrosses

| *Genotype* | *Observed* | *Expected* | | |
|---|---|---|---|---|
| +/+ | 22 | 18.75 | | |
| +/- | 40 | 37.5 | | |
| -/- | 13 | 18.75 | Chi-sqr test = | 0.2874614 |

923large Het Intercrosses

| *Genotype* | *Observed* | *Expected* | | |
|---|---|---|---|---|
| +/+ | 21 | 19.25 | | |
| +/- | 38 | 38.5 | | |
| -/- | 18 | 19.25 | Chi-sqr test = | 0.8839307 |
