## SupplementalTable6 for "An enhancer:involucrin regulatory module impacts human skin barrier adaptation out-of-Africa and modifies atopic dermatitis risk"

**Table S6. Ranked list of differentially expressed genes between 923del/del and WT mice whole skin from RNA-seq.**

| *Feature_ID* | *entrezgene* | *external_gene_name* | *gene_biotype* | *description* | *logFC* | *adj.P.Val* |
|---|---|---|---|---|---|---|
| ENSMUSG00000049128 | 16447 | *Ivl* | protein_coding | involucrin [Source:MGI Symbol;Acc:MGI:96626] | -5.119037 | 1.98E-06 |
| ENSMUSG00000028081 | 20091 | *Rps3a1* | protein_coding | ribosomal protein S3A1 [Source:MGI Symbol;Acc:MGI:1202063] | -4.878743 | 9.60E-14 |
| ENSMUSG00000081355 | NA | *Gm15264* | unprocessed_pseudogene | predicted gene 15264 [Source:MGI Symbol;Acc:MGI:3705845] | 2.481502 | 0.0031793 |
| ENSMUSG00000081303 | NA | *Gm16011* | processed_pseudogene | predicted gene 16011 [Source:MGI Symbol;Acc:MGI:3801796] | 4.154555 | 0.0345667 |
| ENSMUSG00000091383 | NA | *Hist1h2al* | processed_pseudogene | histone cluster 1, H2al [Source:MGI Symbol;Acc:MGI:3646032] | 7.649992 | 8.65E-10 |
| ENSMUSG00000059751 | NA | *Rps3a3* | processed_pseudogene | ribosomal protein S3A3 [Source:MGI Symbol;Acc:MGI:3643406] | 8.451245 | 5.17E-11 |
