## SupplementalTable7 for "An enhancer:involucrin regulatory module impacts human skin barrier adaptation out-of-Africa and modifies atopic dermatitis risk"

**Table S7. Ranked list of differentially expressed genes between 923del/+ and WT mice whole skin from RNA-seq.**

| *Feature_ID* | *entrezgene* | *external_gene_name* | *gene_biotype* | *description* | *logFC* | *adj.P.* |
|---|---|---|---|---|---|---|
| ENSMUSG00000040852 | 213556 | *Plekhh2* | protein_coding | pleckstrin homology domain containing, family H (with MyTH4 domain) member 2 [Source:MGI Symbol;Acc:MGI:2146813] | -4.543948 | 0.017 |
| ENSMUSG00000078122 | NA | *F630028O10Rik* | antisense | RIKEN cDNA F630028O10 gene [Source:MGI Symbol;Acc:MGI:3641813] | -2.150002 | 0.040 |
| ENSMUSG00000111912 | NA | *Gm48521* | lincRNA | predicted gene, 48521 [Source:MGI Symbol;Acc:MGI:6098057] | -2.14202 | 0.036 |
| ENSMUSG00000074634 | 633640 | *Tmem267* | protein_coding | transmembrane protein 267 [Source:MGI Symbol;Acc:MGI:3648543] | -2.009382 | 0.020 |
| ENSMUSG00000049103 | 12772 | *Ccr2* | protein_coding | chemokine (C-C motif) receptor 2 [Source:MGI Symbol;Acc:MGI:106185] | -2.000396 | 0.036 |
| ENSMUSG00000049128 | 16447 | *Ivl* | protein_coding | involucrin [Source:MGI Symbol;Acc:MGI:96626] | -1.826933 | 0.002 |
| ENSMUSG00000039252 | 246316 | *Lgi2* | protein_coding | leucine-rich repeat LGI family, member 2 [Source:MGI Symbol;Acc:MGI:2180196] | 2.103831 | 0.03 |
| ENSMUSG00000035769 | 102448 | *Xylb* | protein_coding | xylulokinase homolog (H. influenzae) [Source:MGI Symbol;Acc:MGI:2142985] | 2.601602 | 0.036 |
| ENSMUSG00000035184 | 629059 | *Fam124a* | protein_coding | family with sequence similarity 124, member A [Source:MGI Symbol;Acc:MGI:3645930] | 2.620848 | 0.047 |
| ENSMUSG00000040289 | 15213 | *Hey1* | protein_coding | hairy/enhancer-of-split related with YRPW motif 1 [Source:MGI Symbol;Acc:MGI:1341800] | 2.710703 | 0.017 |
| ENSMUSG00000016346 | 16536 | *Kcnq2* | protein_coding | potassium voltage-gated channel, subfamily Q, member 2 [Source:MGI Symbol;Acc:MGI:1309503] | 2.924024 | 0.032 |
| ENSMUSG00000066191 | 75691 | *Anks6* | protein_coding | ankyrin repeat and sterile alpha motif domain containing 6 [Source:MGI Symbol;Acc:MGI:1922941] | 2.976234 | 0.012 |
| ENSMUSG00000039137 | 73750 | *Whrn* | protein_coding | whirlin [Source:MGI Symbol;Acc:MGI:2682003] | 3.022618 | 0.000 |
| ENSMUSG00000069227 | 26913 | *Gprin1* | protein_coding | G protein-regulated inducer of neurite outgrowth 1 [Source:MGI Symbol;Acc:MGI:1349455] | 3.50514 | 0.036 |
| ENSMUSG00000006538 | 16147 | *Ihh* | protein_coding | Indian hedgehog [Source:MGI Symbol;Acc:MGI:96533] | 3.881618 | 0.030 |
| ENSMUSG00000027517 | 70065 | *Ankrd60* | protein_coding | ankyrin repeat domain 60 [Source:MGI Symbol;Acc:MGI:1917315] | 3.939413 | 0.043 |
| ENSMUSG00000092675 | NA | *Gm25262* | miRNA | predicted gene, 25262 [Source:MGI Symbol;Acc:MGI:5455039] | 3.973651 | 0.017 |
| ENSMUSG00000098973 | NA | *Mir6236* | miRNA | microRNA 6236 [Source:MGI Symbol;Acc:MGI:5530929] | 4.11892 | 0.040 |
| ENSMUSG00000033948 | 74464 | *Zswim5* | protein_coding | zinc finger SWIM-type containing 5 [Source:MGI Symbol;Acc:MGI:1921714] | 4.23693 | 0.017 |
| ENSMUSG00000076258 | NA | *Gm23935* | miRNA | predicted gene, 23935 [Source:MGI Symbol;Acc:MGI:5453712] | 4.258394 | 0.011 |
| ENSMUSG00000092909 | NA | *Gm25732* | miRNA | predicted gene, 25732 [Source:MGI Symbol;Acc:MGI:5455509] | 4.613336 | 0.023 |
| ENSMUSG00000112365 | NA | *Gm49782* | lincRNA | predicted gene, 49782 [Source:MGI Symbol;Acc:MGI:6215301] | 4.871473 | 0.032 |
| ENSMUSG00000044518 | 30923 | *Foxe3* | protein_coding | forkhead box E3 [Source:MGI Symbol;Acc:MGI:1353569] | 5.295595 | 0.032 |
| ENSMUSG00000091383 | NA | *Hist1h2al* | processed_pseudogene | histone cluster 1, H2al [Source:MGI Symbol;Acc:MGI:3646032] | 7.023739 | 7.31 |
| ENSMUSG00000059751 | NA | *Rps3a3* | processed_pseudogene | ribosomal protein S3A3 [Source:MGI Symbol;Acc:MGI:3643406] | 7.605159 | 9.56 |
