## SupplementalTable8 for "An enhancer:involucrin regulatory module impacts human skin barrier adaptation out-of-Africa and modifies atopic dermatitis risk"

**Table S8. Ranked list of differentially expressed genes between 923large/large and WT mice whole skin from RNA-seq.**

| *Feature_ID* | *entrezgene* | *external_gene_name* | *gene_biotype* | *description* | *logFC* | *adj.* |
|---|---|---|---|---|---|---|
| ENSMUSG00000103523 | NA | *2210017I01Rik* | protein_coding | RIKEN cDNA 2210017I01 gene [Source:MGI Symbol;Acc:MGI:3588251] | -5.881931 | 4 |
| ENSMUSG00000058172 | 16700 | *Krtap6-1* | protein_coding | keratin associated protein 6-1 [Source:MGI Symbol;Acc:MGI:1330228] | -5.700897 | 0.0 |
| ENSMUSG00000028081 | 20091 | *Rps3a1* | protein_coding | ribosomal protein S3A1 [Source:MGI Symbol;Acc:MGI:1202063] | -5.578549 | 1 |
| ENSMUSG00000096534 | 71369 | *Krtap16-3* | protein_coding | keratin associated protein 16-3 [Source:MGI Symbol;Acc:MGI:1918619] | -5.522284 | 0.0 |
| ENSMUSG00000058368 | 170656 | *Krtap21-1* | protein_coding | keratin associated protein 21-1 [Source:MGI Symbol;Acc:MGI:2157767] | -4.793331 | 0.0 |
| ENSMUSG00000086848 | 78382 | *Lce6a* | protein_coding | late cornified envelope 6A [Source:MGI Symbol;Acc:MGI:1925632] | -4.648181 | 3 |
| ENSMUSG00000069306 | 100041230 | *Hist1h4m* | protein_coding | histone cluster 1, H4m [Source:MGI Symbol;Acc:MGI:2448441] | -4.647076 | 0.0 |
| ENSMUSG00000062400 | 68484 | *Krtap6-5* | protein_coding | keratin associated protein 6-5 [Source:MGI Symbol;Acc:MGI:1915734] | -4.541823 | 0.0 |
| ENSMUSG00000068885 | 69520 | *Lce3f* | protein_coding | late cornified envelope 3F [Source:MGI Symbol;Acc:MGI:1916770] | -4.52936 | 0.0 |
| ENSMUSG00000086324 | NA | *Gm15564* | antisense | predicted gene 15564 [Source:MGI Symbol;Acc:MGI:3783013] | -4.385113 | 0.0 |
| ENSMUSG00000095992 | 68740 | *Krtap22-2* | protein_coding | keratin associated protein 22-2 [Source:MGI Symbol;Acc:MGI:1915990] | -4.294082 | 0.0 |
| ENSMUSG00000040852 | 213556 | *Plekhh2* | protein_coding | pleckstrin homology domain containing, family H (with MyTH4 domain) member 2 [Source:MGI Symbol;Acc:MGI:2146813] | -3.916212 | 0.0 |
| ENSMUSG00000057174 | 170939 | *Krtap19-9b* | protein_coding | keratin associated protein 19-9B [Source:MGI Symbol;Acc:MGI:2181750] | -3.873255 | 0 |
| ENSMUSG00000068075 | 100040201 | *Gm10229* | protein_coding | predicted gene 10229 [Source:MGI Symbol;Acc:MGI:3711943] | -3.829142 | 0.0 |
| ENSMUSG00000049128 | 16447 | *Ivl* | protein_coding | involucrin [Source:MGI Symbol;Acc:MGI:96626] | -3.751251 | 1 |
| ENSMUSG00000022931 | 26560 | *Krtap15* | protein_coding | keratin associated protein 15 [Source:MGI Symbol;Acc:MGI:1347350] | -3.739274 | 0 |
| ENSMUSG00000060469 | 77918 | *Krtap19-3* | protein_coding | keratin associated protein 19-3 [Source:MGI Symbol;Acc:MGI:1925168] | -3.581363 | 0.0 |
| ENSMUSG00000081855 | NA | *Rpl17-ps5* | processed_pseudogene | ribosomal protein L17, pseudogene 5 [Source:MGI Symbol;Acc:MGI:3704246] | -3.575253 | 0.0 |
| ENSMUSG00000035202 | 102436 | *Lars2* | protein_coding | leucyl-tRNA synthetase, mitochondrial [Source:MGI Symbol;Acc:MGI:2142973] | -3.404943 | 0.0 |
| ENSMUSG00000068074 | 100040214 | *Gm10228* | protein_coding | predicted gene 10228 [Source:MGI Symbol;Acc:MGI:3704467] | -3.40229 | 0.0 |
| ENSMUSG00000099842 | NA | *Gm4034* | processed_pseudogene | predicted gene 4034 [Source:MGI Symbol;Acc:MGI:3782208] | -3.372075 | 3 |
| ENSMUSG00000059632 | 16703 | *Krtap8-1* | protein_coding | keratin associated protein 8-1 [Source:MGI Symbol;Acc:MGI:1330293] | -3.33129 | 0.0 |
| ENSMUSG00000049809 | 75586 | *Krtap9-3* | protein_coding | keratin associated protein 9-3 [Source:MGI Symbol;Acc:MGI:1922836] | -3.159599 | 0.0 |
| ENSMUSG00000077506 | NA | *Scarna9* | snoRNA | small Cajal body-specific RNA 9 [Source:MGI Symbol;Acc:MGI:3819489] | -3.150323 | 5 |
| ENSMUSG00000027908 | 71325 | *Tchhl1* | protein_coding | trichohyalin-like 1 [Source:MGI Symbol;Acc:MGI:1918575] | -3.047137 | 0.0 |
| ENSMUSG00000117900 | NA | *AC125101.1* | antisense | Novel transcript, antisense to Scd3 | -3.008209 | 0.0 |
| ENSMUSG00000074433 | 69514 | *Lce3e* | protein_coding | late cornified envelope 3E [Source:MGI Symbol;Acc:MGI:1916764] | -3.007469 | 0.0 |
| ENSMUSG00000022229 | 192113 | *Atp12a* | protein_coding | ATPase, H+/K+ transporting, nongastric, alpha polypeptide [Source:MGI Symbol;Acc:MGI:1926943] | -3.000971 | 0.0 |
| ENSMUSG00000017300 | 21925 | *Tnnc2* | protein_coding | troponin C2, fast [Source:MGI Symbol;Acc:MGI:98780] | -2.968038 | 0.0 |
| ENSMUSG00000082699 | NA | *Gm12736* | processed_pseudogene | predicted gene 12736 [Source:MGI Symbol;Acc:MGI:3650723] | -2.899085 | 0.0 |
| ENSMUSG00000087968 | NA | *Gm25395* | scaRNA | predicted gene, 25395 [Source:MGI Symbol;Acc:MGI:5455172] | -2.862775 | 6 |
| ENSMUSG00000024411 | 11829 | *Aqp4* | protein_coding | aquaporin 4 [Source:MGI Symbol;Acc:MGI:107387] | -2.854831 | 0.0 |
| ENSMUSG00000062433 | 16701 | *Krtap6-2* | protein_coding | keratin associated protein 6-2 [Source:MGI Symbol;Acc:MGI:1330280] | -2.849751 | 0.0 |
| ENSMUSG00000030237 | 28250 | *Slco1a4* | protein_coding | solute carrier organic anion transporter family, member 1a4 [Source:MGI Symbol;Acc:MGI:1351896] | -2.817944 | 0.0 |
| ENSMUSG00000079641 | 67248 | *Rpl39* | protein_coding | ribosomal protein L39 [Source:MGI Symbol;Acc:MGI:1914498] | -2.786459 | 9 |
| ENSMUSG00000069308 | 319188 | *Hist1h2bp* | protein_coding | histone cluster 1, H2bp [Source:MGI Symbol;Acc:MGI:2448409] | -2.784327 | 8 |
| ENSMUSG00000065037 | 19817 | *Rn7sk* | misc_RNA | RNA, 7SK, nuclear [Source:MGI Symbol;Acc:MGI:103186] | -2.73028 | 3 |
| ENSMUSG00000089281 | NA | *Scarna6* | scaRNA | small Cajal body-specific RNA 6 [Source:MGI Symbol;Acc:MGI:3819487] | -2.724599 | 8 |
| ENSMUSG00000069722 | 66380 | *Krtap3-3* | protein_coding | keratin associated protein 3-3 [Source:MGI Symbol;Acc:MGI:1913630] | -2.719274 | 0.0 |
| ENSMUSG00000057322 | 67671 | *Rpl38* | protein_coding | ribosomal protein L38 [Source:MGI Symbol;Acc:MGI:1914921] | -2.699279 | 5 |
| ENSMUSG00000092837 | NA | *Rpph1* | ribozyme | ribonuclease P RNA component H1 [Source:MGI Symbol;Acc:MGI:1934664] | -2.685727 | 2 |
| ENSMUSG00000064999 | NA | *Gm26035* | misc_RNA | predicted gene, 26035 [Source:MGI Symbol;Acc:MGI:5455812] | -2.675681 | 0.0 |
| ENSMUSG00000069309 | 319170 | *Hist1h2an* | protein_coding | histone cluster 1, H2an [Source:MGI Symbol;Acc:MGI:2448300] | -2.660542 | 1 |
| ENSMUSG00000001773 | 53320 | *Folh1* | protein_coding | folate hydrolase 1 [Source:MGI Symbol;Acc:MGI:1858193] | -2.552246 | 0 |
| ENSMUSG00000115016 | 102636984 | *AC158982.1* | processed_transcript | novel transcript | -2.542871 | 0.0 |
| ENSMUSG00000036231 | 403205 | *Agr3* | protein_coding | anterior gradient 3 [Source:MGI Symbol;Acc:MGI:2685734] | -2.527515 | 0 |
| ENSMUSG00000067288 | 54127 | *Rps28* | protein_coding | ribosomal protein S28 [Source:MGI Symbol;Acc:MGI:1859516] | -2.520922 | 5 |
| ENSMUSG00000047246 | 319179 | *Hist1h2be* | protein_coding | histone cluster 1, H2be [Source:MGI Symbol;Acc:MGI:2448380] | -2.504356 | 8 |
| ENSMUSG00000047253 | 69664 | *Krtap1-5* | protein_coding | keratin associated protein 1-5 [Source:MGI Symbol;Acc:MGI:1916914] | -2.503888 | 0.0 |
| ENSMUSG00000079013 | 238395 | *Serpina3j* | protein_coding | serine (or cysteine) peptidase inhibitor, clade A (alpha-1 antiproteinase, antitrypsin), member 3J [Source:MGI Symbol;Acc:MGI:2182 | -2.450798 | 0.0 |
| ENSMUSG00000093314 | NA | *Mir5136* | miRNA | microRNA 5136 [Source:MGI Symbol;Acc:MGI:4950461] | -2.411673 | 0.0 |
| ENSMUSG00000056706 | 71363 | *Krtap7-1* | protein_coding | keratin associated protein 7-1 [Source:MGI Symbol;Acc:MGI:1918613] | -2.406509 | 0.0 |
| ENSMUSG00000101355 | 319152 | *Hist1h3h* | protein_coding | histone cluster 1, H3h [Source:MGI Symbol;Acc:MGI:2448349] | -2.396619 | 7 |
| ENSMUSG00000060981 | 69386 | *Hist1h4h* | protein_coding | histone cluster 1, H4h [Source:MGI Symbol;Acc:MGI:2448427] | -2.395177 | 1 |
| ENSMUSG00000068855 | 319176 | *Hist2h2ac* | protein_coding | histone cluster 2, H2ac [Source:MGI Symbol;Acc:MGI:2448316] | -2.390388 | 1 |
| ENSMUSG00000103084 | NA | *Gm38119* | protein_coding | predicted gene, 38119 [Source:MGI Symbol;Acc:MGI:5611347] | -2.387959 | 0.0 |
| ENSMUSG00000023078 | 55985 | *Cxcl13* | protein_coding | chemokine (C-X-C motif) ligand 13 [Source:MGI Symbol;Acc:MGI:1888499] | -2.384844 | 0.0 |
| ENSMUSG00000047894 | 11731 | *Ang2* | protein_coding | angiogenin, ribonuclease A family, member 2 [Source:MGI Symbol;Acc:MGI:104984] | -2.382766 | 0.0 |
| ENSMUSG00000071478 | 319165 | *Hist1h2ad* | protein_coding | histone cluster 1, H2ad [Source:MGI Symbol;Acc:MGI:2448289] | -2.374623 | 7 |
| ENSMUSG00000060678 | 319155 | *Hist1h4c* | protein_coding | histone cluster 1, H4c [Source:MGI Symbol;Acc:MGI:2448421] | -2.371969 | 6 |
| ENSMUSG00000067455 | 319159 | *Hist1h4j* | protein_coding | histone cluster 1, H4j [Source:MGI Symbol;Acc:MGI:2448436] | -2.368957 | 9 |
| ENSMUSG00000093218 | NA | *Gm25252* | miRNA | predicted gene, 25252 [Source:MGI Symbol;Acc:MGI:5455029] | -2.341425 | 0.0 |
| ENSMUSG00000015665 | 245533 | *Awat1* | protein_coding | acyl-CoA wax alcohol acyltransferase 1 [Source:MGI Symbol;Acc:MGI:3588200] | -2.331783 | 0.0 |
| ENSMUSG00000056270 | 109314 | *Prr9* | protein_coding | proline rich 9 [Source:MGI Symbol;Acc:MGI:1925680] | -2.328088 | 0.0 |
| ENSMUSG00000069267 | 319150 | *Hist1h3b* | protein_coding | histone cluster 1, H3b [Source:MGI Symbol;Acc:MGI:2448319] | -2.305767 | 2 |
| ENSMUSG00000045566 | 229562 | *Sprr4* | protein_coding | small proline-rich protein 4 [Source:MGI Symbol;Acc:MGI:2654508] | -2.301675 | 0.0 |
| ENSMUSG00000064694 | NA | *Gm24146* | misc_RNA | predicted gene, 24146 [Source:MGI Symbol;Acc:MGI:5453923] | -2.266976 | 1 |
| ENSMUSG00000028011 | 56720 | *Tdo2* | protein_coding | tryptophan 2,3-dioxygenase [Source:MGI Symbol;Acc:MGI:1928486] | -2.259513 | 0.0 |
| ENSMUSG00000101972 | 319153 | *Hist1h3i* | protein_coding | histone cluster 1, H3i [Source:MGI Symbol;Acc:MGI:2448350] | -2.25449 | 5 |
| ENSMUSG00000058385 | 319181 | *Hist1h2bg* | protein_coding | histone cluster 1, H2bg [Source:MGI Symbol;Acc:MGI:2448386] | -2.236547 | 1 |
| ENSMUSG00000041841 | 100502825 | *Rpl37* | protein_coding | ribosomal protein L37 [Source:MGI Symbol;Acc:MGI:1914531] | -2.229471 | 5 |
| ENSMUSG00000041841 | 67281 | *Rpl37* | protein_coding | ribosomal protein L37 [Source:MGI Symbol;Acc:MGI:1914531] | -2.229471 | 5 |
| ENSMUSG00000064899 | NA | *Snord118* | snoRNA | small nucleolar RNA, C/D box 118 [Source:MGI Symbol;Acc:MGI:3819519] | -2.225969 | 0.0 |
| ENSMUSG00000094443 | 244495 | *Sgo2b* | protein_coding | shugoshin 2B [Source:MGI Symbol;Acc:MGI:3644562] | -2.22534 | 0.0 |
| ENSMUSG00000062727 | 319184 | *Hist1h2bk* | protein_coding | histone cluster 1, H2bk [Source:MGI Symbol;Acc:MGI:2448399] | -2.217886 | 1 |
| ENSMUSG00000069972 | NA | *Rps13-ps2* | processed_pseudogene | ribosomal protein S13, pseudogene 2 [Source:MGI Symbol;Acc:MGI:3704295] | -2.208372 | 0.0 |
| ENSMUSG00000078655 | NA | *Gm10972* | protein_coding | predicted gene 10972 [Source:MGI Symbol;Acc:MGI:3779183] | -2.190514 | 0.0 |
| ENSMUSG00000101389 | 666907 | *Ms4a4a* | protein_coding | membrane-spanning 4-domains, subfamily A, member 4A [Source:MGI Symbol;Acc:MGI:3643932] | -2.189783 | 0.0 |
| ENSMUSG00000066364 | 271047 | *Serpina3b* | protein_coding | serine (or cysteine) peptidase inhibitor, clade A, member 3B [Source:MGI Symbol;Acc:MGI:2182835] | -2.174769 | 0 |
| ENSMUSG00000027824 | 56544 | *Vmn2r1* | protein_coding | vomeronasal 2, receptor 1 [Source:MGI Symbol;Acc:MGI:3645892] | -2.172085 | 0.0 |
| ENSMUSG00000028359 | 18407 | *Orm3* | protein_coding | orosomucoid 3 [Source:MGI Symbol;Acc:MGI:97445] | -2.161917 | 0.0 |
| ENSMUSG00000018102 | 68024 | *Hist1h2bc* | protein_coding | histone cluster 1, H2bc [Source:MGI Symbol;Acc:MGI:1915274] | -2.146902 | 1 |
| ENSMUSG00000069305 | 319161 | *Hist1h4n* | protein_coding | histone cluster 1, H4n [Source:MGI Symbol;Acc:MGI:4843992] | -2.139598 | 0.0 |
| ENSMUSG00000048483 | 238331 | *Zdhhc22* | protein_coding | zinc finger, DHHC-type containing 22 [Source:MGI Symbol;Acc:MGI:2685108] | -2.131746 | 0.0 |
| ENSMUSG00000069302 | 319168 | *Hist1h2ah* | protein_coding | histone cluster 1, H2ah [Source:MGI Symbol;Acc:MGI:2448295] | -2.116599 | 1 |
| ENSMUSG00000069266 | 326620 | *Hist1h4b* | protein_coding | histone cluster 1, H4b [Source:MGI Symbol;Acc:MGI:2448420] | -2.104418 | 7 |
| ENSMUSG00000052013 | 208154 | *Btla* | protein_coding | B and T lymphocyte associated [Source:MGI Symbol;Acc:MGI:2658978] | -2.079617 | 0.0 |
| ENSMUSG00000056054 | 20201 | *S100a8* | protein_coding | S100 calcium binding protein A8 (calgranulin A) [Source:MGI Symbol;Acc:MGI:88244] | -2.07057 | 0.0 |
| ENSMUSG00000039760 | 237310 | *Il22ra2* | protein_coding | interleukin 22 receptor, alpha 2 [Source:MGI Symbol;Acc:MGI:2665114] | -2.046628 | 0.0 |
| ENSMUSG00000094338 | 319185 | *Hist1h2bl* | protein_coding | histone cluster 1, H2bl [Source:MGI Symbol;Acc:MGI:2448403] | -2.043779 | 7 |
| ENSMUSG00000114456 | 319182 | *Hist1h2bh* | protein_coding | histone cluster 1, H2bh [Source:MGI Symbol;Acc:MGI:2448387] | -2.033872 | 1 |
| ENSMUSG00000114279 | 319186 | *Hist1h2bm* | protein_coding | histone cluster 1, H2bm [Source:MGI Symbol;Acc:MGI:2448404] | -2.02139 | 1 |
| ENSMUSG00000037563 | 20055 | *Rps16* | protein_coding | ribosomal protein S16 [Source:MGI Symbol;Acc:MGI:98118] | -2.004548 | 2 |
| ENSMUSG00000078252 | 77914 | *Krtap17-1* | protein_coding | keratin associated protein 17-1 [Source:MGI Symbol;Acc:MGI:1925164] | -2.000689 | 0.0 |
| ENSMUSG00000108790 | NA | *Gm44806* | TEC | predicted gene 44806 [Source:MGI Symbol;Acc:MGI:5753382] | 2.014853 | 0.0 |
| ENSMUSG00000023176 | 71756 | *Cpn2* | protein_coding | carboxypeptidase N, polypeptide 2 [Source:MGI Symbol;Acc:MGI:1919006] | 2.016662 | 0.0 |
| ENSMUSG00000038522 | 215929 | *Mfsd4b1* | protein_coding | major facilitator superfamily domain containing 4B1 [Source:MGI Symbol;Acc:MGI:2143575] | 2.018912 | 0.0 |
| ENSMUSG00000005373 | 58805 | *Mlxipl* | protein_coding | MLX interacting protein-like [Source:MGI Symbol;Acc:MGI:1927999] | 2.019612 | 0.0 |
| ENSMUSG00000000183 | 14177 | *Fgf6* | protein_coding | fibroblast growth factor 6 [Source:MGI Symbol;Acc:MGI:95520] | 2.025462 | 0.0 |

| | | | | | | |
|---|---|---|---|---|---|---|
| ENSMUSG00000097442 | NA | *Gm26632* | antisense | predicted gene, 26632 [Source:MGI Symbol;Acc:MGI:5477126] | 2.028084 | 0.[illegible] |
| ENSMUSG00000079261 | NA | *Gm15217* | protein_coding | predicted gene 15217 [Source:MGI Symbol;Acc:MGI:3705233] | 2.055163 | 0.[illegible] |
| ENSMUSG00000040146 | 71746 | *Rgl3* | protein_coding | ral guanine nucleotide dissociation stimulator-like 3 [Source:MGI Symbol;Acc:MGI:1918996] | 2.07165 | 0.[illegible] |
| ENSMUSG00000028427 | 11832 | *Aqp7* | protein_coding | aquaporin 7 [Source:MGI Symbol;Acc:MGI:1314647] | 2.07465 | 0[illegible] |
| ENSMUSG00000038508 | 23886 | *Gdf15* | protein_coding | growth differentiation factor 15 [Source:MGI Symbol;Acc:MGI:1346047] | 2.090234 | 0.[illegible] |
| ENSMUSG00000030546 | 103968 | *Plin1* | protein_coding | perilipin 1 [Source:MGI Symbol;Acc:MGI:1890505] | 2.091874 | 0.[illegible] |
| ENSMUSG00000086167 | NA | *Gm13827* | processed_pseudogene | predicted gene 13827 [Source:MGI Symbol;Acc:MGI:3651518] | 2.093876 | 0.[illegible] |
| ENSMUSG00000020963 | 22095 | *Tshr* | protein_coding | thyroid stimulating hormone receptor [Source:MGI Symbol;Acc:MGI:98849] | 2.111705 | 0.[illegible] |
| ENSMUSG00000094012 | 100009614 | *Gm10024* | protein_coding | predicted gene 10024 [Source:MGI Symbol;Acc:MGI:3641784] | 2.117729 | 0.[illegible] |
| ENSMUSG00000044667 | 229791 | *Plppr4* | protein_coding | phospholipid phosphatase related 4 [Source:MGI Symbol;Acc:MGI:106530] | 2.122175 | 0[illegible] |
| ENSMUSG00000027335 | 11550 | *Adra1d* | protein_coding | adrenergic receptor, alpha 1d [Source:MGI Symbol;Acc:MGI:106673] | 2.129465 | 0.[illegible] |
| ENSMUSG00000097983 | NA | *Gm26971* | sense_intronic | predicted gene, 26971 [Source:MGI Symbol;Acc:MGI:5504086] | 2.146034 | 0.[illegible] |
| ENSMUSG00000095817 | 100041261 | *Gm3238* | protein_coding | predicted gene 3238 [Source:MGI Symbol;Acc:MGI:3781416] | 2.146129 | 0.[illegible] |
| ENSMUSG00000085794 | NA | *Vax2os* | antisense | ventral anterior homeobox 2, opposite strand [Source:MGI Symbol;Acc:MGI:3583301] | 2.148413 | 0.[illegible] |
| ENSMUSG00000104971 | NA | *9430087B13Rik* | TEC | RIKEN cDNA 9430087B13 gene [Source:MGI Symbol;Acc:MGI:1924687] | 2.169937 | 0.[illegible] |
| ENSMUSG00000004892 | 12032 | *Bcan* | protein_coding | brevican [Source:MGI Symbol;Acc:MGI:1096385] | 2.188172 | 0.[illegible] |
| ENSMUSG00000020926 | 11488 | *Adam11* | protein_coding | a disintegrin and metallopeptidase domain 11 [Source:MGI Symbol;Acc:MGI:1098667] | 2.188704 | 0.[illegible] |
| ENSMUSG00000018554 | 53422 | *Ybx2* | protein_coding | Y box protein 2 [Source:MGI Symbol;Acc:MGI:1096372] | 2.192596 | 0.[illegible] |
| ENSMUSG00000101588 | NA | *Gm28265* | lincRNA | predicted gene 28265 [Source:MGI Symbol;Acc:MGI:5578971] | 2.194802 | 0.[illegible] |
| ENSMUSG00000054667 | 16370 | *Irs4* | protein_coding | insulin receptor substrate 4 [Source:MGI Symbol;Acc:MGI:1338009] | 2.195335 | 0.[illegible] |
| ENSMUSG00000104860 | NA | *Gm42510* | TEC | predicted gene 42510 [Source:MGI Symbol;Acc:MGI:5662647] | 2.210299 | 0.[illegible] |
| ENSMUSG00000027983 | 71519 | *Cyp2u1* | protein_coding | cytochrome P450, family 2, subfamily u, polypeptide 1 [Source:MGI Symbol;Acc:MGI:1918769] | 2.220056 | 0.[illegible] |
| ENSMUSG00000020264 | 246049 | *Slc36a2* | protein_coding | solute carrier family 36 (proton/amino acid symporter), member 2 [Source:MGI Symbol;Acc:MGI:1891430] | 2.243022 | 0.[illegible] |
| ENSMUSG00000113764 | NA | *Gm48617* | sense_intronic | predicted gene, 48617 [Source:MGI Symbol;Acc:MGI:6098210] | 2.247111 | 0.[illegible] |
| ENSMUSG00000110631 | NA | *Gm42047* | lincRNA | predicted gene, 42047 [Source:MGI Symbol;Acc:MGI:5624932] | 2.252189 | 0.[illegible] |
| ENSMUSG00000107794 | NA | *Gm44095* | TEC | predicted gene, 44095 [Source:MGI Symbol;Acc:MGI:5690487] | 2.256119 | 0.[illegible] |
| ENSMUSG00000112170 | 670895 | *Gm9508* | protein_coding | predicted gene 9508 [Source:MGI Symbol;Acc:MGI:3779918] | 2.281089 | 0.[illegible] |
| ENSMUSG00000026834 | 269275 | *Acvr1c* | protein_coding | activin A receptor, type IC [Source:MGI Symbol;Acc:MGI:2661081] | 2.289884 | 0.[illegible] |
| ENSMUSG00000087104 | 67351 | *Tmem132cos* | antisense | transmembrane protein 132C, opposite strand [Source:MGI Symbol;Acc:MGI:1914601] | 2.300357 | 0.[illegible] |
| ENSMUSG00000031535 | 234130 | *Dkk4* | protein_coding | dickkopf WNT signaling pathway inhibitor 4 [Source:MGI Symbol;Acc:MGI:2385299] | 2.305373 | 0.[illegible] |
| ENSMUSG00000045875 | 11549 | *Adra1a* | protein_coding | adrenergic receptor, alpha 1a [Source:MGI Symbol;Acc:MGI:104773] | 2.323263 | 0.[illegible] |
| ENSMUSG00000102461 | NA | *Gm37166* | TEC | predicted gene, 37166 [Source:MGI Symbol;Acc:MGI:5610394] | 2.334491 | 0.[illegible] |
| ENSMUSG00000112830 | NA | *Gm47765* | processed_transcript | predicted gene, 47765 [Source:MGI Symbol;Acc:MGI:6096917] | 2.340376 | 0.[illegible] |
| ENSMUSG00000083382 | NA | *Gm6433* | processed_pseudogene | predicted gene 6433 [Source:MGI Symbol;Acc:MGI:3645394] | 2.343639 | 0[illegible] |
| ENSMUSG00000049241 | 243270 | *Hcar1* | protein_coding | hydrocarboxylic acid receptor 1 [Source:MGI Symbol;Acc:MGI:2441671] | 2.375732 | 0[illegible] |
| ENSMUSG00000032387 | 71973 | *Rbpms2* | protein_coding | RNA binding protein with multiple splicing 2 [Source:MGI Symbol;Acc:MGI:1919223] | 2.390549 | 0.[illegible] |
| ENSMUSG00000031489 | 11556 | *Adrb3* | protein_coding | adrenergic receptor, beta 3 [Source:MGI Symbol;Acc:MGI:87939] | 2.391318 | 0.[illegible] |
| ENSMUSG00000023019 | 14555 | *Gpd1* | protein_coding | glycerol-3-phosphate dehydrogenase 1 (soluble) [Source:MGI Symbol;Acc:MGI:95679] | 2.410334 | 0.[illegible] |
| ENSMUSG00000118330 | NA | *AC132148.1* | antisense | novel transcript, antisense to Dagla | 2.414843 | 0.[illegible] |
| ENSMUSG00000049265 | 16527 | *Kcnk3* | protein_coding | potassium channel, subfamily K, member 3 [Source:MGI Symbol;Acc:MGI:1100509] | 2.437493 | 0.[illegible] |
| ENSMUSG00000033882 | 633285 | *Rbm46* | protein_coding | RNA binding motif protein 46 [Source:MGI Symbol;Acc:MGI:3645057] | 2.455302 | 0.[illegible] |
| ENSMUSG00000108141 | NA | *Gm44079* | antisense | predicted gene, 44079 [Source:MGI Symbol;Acc:MGI:5690471] | 2.500186 | 0.[illegible] |
| ENSMUSG00000048070 | 193003 | *Pirt* | protein_coding | phosphoinositide-interacting regulator of transient receptor potential channels [Source:MGI Symbol;Acc:MGI:2443635] | 2.542469 | 0[illegible] |
| ENSMUSG00000096421 | 100502953 | *Gm10100* | protein_coding | predicted gene 10100 [Source:MGI Symbol;Acc:MGI:3642388] | 2.679939 | 0[illegible] |
| ENSMUSG00000064225 | 75552 | *Paqr9* | protein_coding | progestin and adipoQ receptor family member IX [Source:MGI Symbol;Acc:MGI:1922802] | 2.726758 | 0.[illegible] |
| ENSMUSG00000026347 | 72160 | *Tmem163* | protein_coding | transmembrane protein 163 [Source:MGI Symbol;Acc:MGI:1919410] | 2.736005 | 0.[illegible] |
| ENSMUSG00000067017 | NA | *Gm3608* | pseudogene | predicted gene 3608 [Source:MGI Symbol;Acc:MGI:3804932] | 2.797161 | 0.[illegible] |
| ENSMUSG00000090955 | NA | *Gm17097* | processed_pseudogene | predicted gene 17097 [Source:MGI Symbol;Acc:MGI:4937924] | 2.827473 | 0.[illegible] |
| ENSMUSG00000104696 | NA | *Gm42946* | TEC | predicted gene 42946 [Source:MGI Symbol;Acc:MGI:5663083] | 2.986255 | 0.[illegible] |
| ENSMUSG00000081303 | NA | *Gm16011* | processed_pseudogene | predicted gene 16011 [Source:MGI Symbol;Acc:MGI:3801796] | 3.386614 | 0.[illegible] |
| ENSMUSG00000087382 | NA | *Ctcflos* | antisense | CCCTC-binding factor (zinc finger protein)-like, opposite strand [Source:MGI Symbol;Acc:MGI:1921411] | 3.479359 | 0.[illegible] |
| ENSMUSG00000069584 | NA | *Gm10272* | protein_coding | predicted gene 10272 [Source:MGI Symbol;Acc:MGI:3642183] | 3.502495 | 0.[illegible] |
| ENSMUSG00000078234 | 242721 | *Klhdc7a* | protein_coding | kelch domain containing 7A [Source:MGI Symbol;Acc:MGI:2444612] | 3.649402 | 0.[illegible] |
| ENSMUSG00000032401 | 235435 | *Lctl* | protein_coding | lactase-like [Source:MGI Symbol;Acc:MGI:2183549] | 3.699934 | 0.[illegible] |
| ENSMUSG00000055197 | 260298 | *Fev* | protein_coding | FEV (ETS oncogene family) [Source:MGI Symbol;Acc:MGI:2449712] | 3.973698 | 0.[illegible] |
| ENSMUSG00000115702 | NA | *Gm5206* | transcribed_processed_pseudogene | predicted pseudogene 5206 [Source:MGI Symbol;Acc:MGI:3645516] | 4.183463 | 4[illegible] |
| ENSMUSG00000091383 | NA | *Hist1h2al* | processed_pseudogene | histone cluster 1, H2al [Source:MGI Symbol;Acc:MGI:3646032] | 7.454856 | 4[illegible] |
| ENSMUSG00000059751 | NA | *Rps3a3* | processed_pseudogene | ribosomal protein S3A3 [Source:MGI Symbol;Acc:MGI:3643406] | 7.741016 | 1[illegible] |
