## SupplementalTable8 for "An enhancer:involucrin regulatory module impacts human skin barrier adaptation out-of-Africa and modifies atopic dermatitis risk"

**Table S9. Ranked list of differentially expressed genes between 923large/+ and WT mice whole skin from RNA-seq.**

| *Feature_ID* | *entrezgene* | *external_gene_name* | *gene_biotype* | *description* | *logFC* | *adj* |
|---|---|---|---|---|---|---|
| ENSMUSG00000069306 | 100041230 | *Hist1h4m* | protein_coding | histone cluster 1, H4m [Source:MGI Symbol;Acc:MGI:2448441] | -7.401242 | 0. |
| ENSMUSG00000086324 | NA | *Gm15564* | antisense | predicted gene 15564 [Source:MGI Symbol;Acc:MGI:3783013] | -5.188587 | 0. |
| ENSMUSG00000027908 | 71325 | *Tchhl1* | protein_coding | trichohyalin-like 1 [Source:MGI Symbol;Acc:MGI:1918575] | -5.015061 | 0. |
| ENSMUSG00000058368 | 170656 | *Krtap21-1* | protein_coding | keratin associated protein 21-1 [Source:MGI Symbol;Acc:MGI:2157767] | -4.939582 | 0. |
| ENSMUSG00000096534 | 71369 | *Krtap16-3* | protein_coding | keratin associated protein 16-3 [Source:MGI Symbol;Acc:MGI:1918619] | -4.892855 | 0. |
| ENSMUSG00000113267 | NA | *Gm47969* | protein_coding | predicted gene, 47969 [Source:MGI Symbol;Acc:MGI:6097249] | -4.874263 | 0. |
| ENSMUSG00000057174 | 170939 | *Krtap19-9b* | protein_coding | keratin associated protein 19-9B [Source:MGI Symbol;Acc:MGI:2181750] | -4.071018 | 0. |
| ENSMUSG00000040852 | 213556 | *Plekhh2* | protein_coding | pleckstrin homology domain containing, family H (with MyTH4 domain) member 2 [Source:MGI Symbol;Acc:MGI:2146813] | -3.948906 | 0. |
| ENSMUSG00000068075 | 100040201 | *Gm10229* | protein_coding | predicted gene 10229 [Source:MGI Symbol;Acc:MGI:3711943] | -3.790833 | 0. |
| ENSMUSG00000035202 | 102436 | *Lars2* | protein_coding | leucyl-tRNA synthetase, mitochondrial [Source:MGI Symbol;Acc:MGI:2142973] | -3.6555 | 0. |
| ENSMUSG00000060469 | 77918 | *Krtap19-3* | protein_coding | keratin associated protein 19-3 [Source:MGI Symbol;Acc:MGI:1925168] | -3.648565 | 0. |
| ENSMUSG00000045566 | 229562 | *Sprr4* | protein_coding | small proline-rich protein 4 [Source:MGI Symbol;Acc:MGI:2654508] | -3.592875 | 0. |
| ENSMUSG00000058172 | 16700 | *Krtap6-1* | protein_coding | keratin associated protein 6-1 [Source:MGI Symbol;Acc:MGI:1330228] | -3.373585 | 0. |
| ENSMUSG00000062456 | NA | *Rpl9-ps6* | protein_coding | ribosomal protein L9, pseudogene 6 [Source:MGI Symbol;Acc:MGI:3642682] | -3.182746 | 0. |
| ENSMUSG00000077506 | NA | *Scarna9* | snoRNA | small Cajal body-specific RNA 9 [Source:MGI Symbol;Acc:MGI:3819489] | -3.033554 | 1 |
| ENSMUSG00000017300 | 21925 | *Tnnc2* | protein_coding | troponin C2, fast [Source:MGI Symbol;Acc:MGI:98780] | -3.018984 | 0. |
| ENSMUSG00000069309 | 319170 | *Hist1h2an* | protein_coding | histone cluster 1, H2an [Source:MGI Symbol;Acc:MGI:2448300] | -3.01664 | 1 |
| ENSMUSG00000062400 | 68484 | *Krtap6-5* | protein_coding | keratin associated protein 6-5 [Source:MGI Symbol;Acc:MGI:1915734] | -3.010397 | 0. |
| ENSMUSG00000069308 | 319188 | *Hist1h2bp* | protein_coding | histone cluster 1, H2bp [Source:MGI Symbol;Acc:MGI:2448409] | -2.996099 | 1 |
| ENSMUSG00000099842 | NA | *Gm4034* | processed_pseudogene | predicted gene 4034 [Source:MGI Symbol;Acc:MGI:3782208] | -2.972916 | 0. |
| ENSMUSG00000057322 | 67671 | *Rpl38* | protein_coding | ribosomal protein L38 [Source:MGI Symbol;Acc:MGI:1914921] | -2.845785 | 1 |
| ENSMUSG00000062727 | 319184 | *Hist1h2bk* | protein_coding | histone cluster 1, H2bk [Source:MGI Symbol;Acc:MGI:2448399] | -2.759047 | 2 |
| ENSMUSG00000095992 | 68740 | *Krtap22-2* | protein_coding | keratin associated protein 22-2 [Source:MGI Symbol;Acc:MGI:1915990] | -2.743409 | 0. |
| ENSMUSG00000087968 | NA | *Gm25395* | scaRNA | predicted gene, 25395 [Source:MGI Symbol;Acc:MGI:5455172] | -2.67975 | 3 |
| ENSMUSG00000069266 | 326620 | *Hist1h4b* | protein_coding | histone cluster 1, H4b [Source:MGI Symbol;Acc:MGI:2448420] | -2.675207 | 1 |
| ENSMUSG00000069972 | NA | *Rps13-ps2* | processed_pseudogene | ribosomal protein S13, pseudogene 2 [Source:MGI Symbol;Acc:MGI:3704295] | -2.663459 | 0. |
| ENSMUSG00000068074 | 100040214 | *Gm10228* | protein_coding | predicted gene 10228 [Source:MGI Symbol;Acc:MGI:3704467] | -2.660009 | 0 |
| ENSMUSG00000067288 | 54127 | *Rps28* | protein_coding | ribosomal protein S28 [Source:MGI Symbol;Acc:MGI:1859516] | -2.657763 | 1 |
| ENSMUSG00000109973 | NA | *Gm45397* | lincRNA | predicted gene 45397 [Source:MGI Symbol;Acc:MGI:5791233] | -2.635053 | 0. |
| ENSMUSG00000047246 | 319179 | *Hist1h2be* | protein_coding | histone cluster 1, H2be [Source:MGI Symbol;Acc:MGI:2448380] | -2.577282 | 1 |
| ENSMUSG00000069267 | 319150 | *Hist1h3b* | protein_coding | histone cluster 1, H3b [Source:MGI Symbol;Acc:MGI:2448319] | -2.549473 | 1 |
| ENSMUSG00000101355 | 319152 | *Hist1h3h* | protein_coding | histone cluster 1, H3h [Source:MGI Symbol;Acc:MGI:2448349] | -2.523819 | 1 |
| ENSMUSG00000092837 | NA | *Rpph1* | ribozyme | ribonuclease P RNA component H1 [Source:MGI Symbol;Acc:MGI:1934664] | -2.518094 | 1 |
| ENSMUSG00000043753 | 242523 | *Dmrta1* | protein_coding | doublesex and mab-3 related transcription factor like family A1 [Source:MGI Symbol;Acc:MGI:2653627] | -2.517907 | 0. |
| ENSMUSG00000093218 | NA | *Gm25252* | miRNA | predicted gene, 25252 [Source:MGI Symbol;Acc:MGI:5455029] | -2.480508 | 0. |
| ENSMUSG00000071478 | 319165 | *Hist1h2ad* | protein_coding | histone cluster 1, H2ad [Source:MGI Symbol;Acc:MGI:2448289] | -2.477184 | 8 |
| ENSMUSG00000079641 | 67248 | *Rpl39* | protein_coding | ribosomal protein L39 [Source:MGI Symbol;Acc:MGI:1914498] | -2.473789 | 1 |
| ENSMUSG00000059632 | 16703 | *Krtap8-1* | protein_coding | keratin associated protein 8-1 [Source:MGI Symbol;Acc:MGI:1330293] | -2.460032 | 0. |
| ENSMUSG00000067455 | 319159 | *Hist1h4j* | protein_coding | histone cluster 1, H4j [Source:MGI Symbol;Acc:MGI:2448436] | -2.453508 | 1 |
| ENSMUSG00000101972 | 319153 | *Hist1h3i* | protein_coding | histone cluster 1, H3i [Source:MGI Symbol;Acc:MGI:2448350] | -2.444522 | 5 |
| ENSMUSG00000061584 | 332427 | *Lyg2* | protein_coding | lysozyme G-like 2 [Source:MGI Symbol;Acc:MGI:2685622] | -2.443755 | 0. |
| ENSMUSG00000060981 | 69386 | *Hist1h4h* | protein_coding | histone cluster 1, H4h [Source:MGI Symbol;Acc:MGI:2448427] | -2.441691 | 2 |
| ENSMUSG00000065037 | 19817 | *Rn7sk* | misc_RNA | RNA, 7SK, nuclear [Source:MGI Symbol;Acc:MGI:103186] | -2.43123 | 8 |
| ENSMUSG00000060678 | 319155 | *Hist1h4c* | protein_coding | histone cluster 1, H4c [Source:MGI Symbol;Acc:MGI:2448421] | -2.425398 | 1 |
| ENSMUSG00000114279 | 319186 | *Hist1h2bm* | protein_coding | histone cluster 1, H2bm [Source:MGI Symbol;Acc:MGI:2448404] | -2.333092 | 4 |
| ENSMUSG00000069722 | 66380 | *Krtap3-3* | protein_coding | keratin associated protein 3-3 [Source:MGI Symbol;Acc:MGI:1913630] | -2.319279 | 0. |
| ENSMUSG00000089281 | NA | *Scarna6* | scaRNA | small Cajal body-specific RNA 6 [Source:MGI Symbol;Acc:MGI:3819487] | -2.305016 | 1 |
| ENSMUSG00000056054 | 20201 | *S100a8* | protein_coding | S100 calcium binding protein A8 (calgranulin A) [Source:MGI Symbol;Acc:MGI:88244] | -2.268103 | 0. |
| ENSMUSG00000041841 | 100502825 | *Rpl37* | protein_coding | ribosomal protein L37 [Source:MGI Symbol;Acc:MGI:1914531] | -2.257742 | 1 |
| ENSMUSG00000041841 | 67281 | *Rpl37* | protein_coding | ribosomal protein L37 [Source:MGI Symbol;Acc:MGI:1914531] | -2.257742 | 1 |
| ENSMUSG00000114456 | 319182 | *Hist1h2bh* | protein_coding | histone cluster 1, H2bh [Source:MGI Symbol;Acc:MGI:2448387] | -2.254283 | 1 |
| ENSMUSG00000064694 | NA | *Gm24146* | misc_RNA | predicted gene, 24146 [Source:MGI Symbol;Acc:MGI:5453923] | -2.248391 | 2 |
| ENSMUSG00000058385 | 319181 | *Hist1h2bg* | protein_coding | histone cluster 1, H2bg [Source:MGI Symbol;Acc:MGI:2448386] | -2.236667 | 2 |
| ENSMUSG00000069301 | 319167 | *Hist1h2ag* | protein_coding | histone cluster 1, H2ag [Source:MGI Symbol;Acc:MGI:2448293] | -2.231177 | 1 |
| ENSMUSG00000021403 | 20706 | *Serpinb9b* | protein_coding | serine (or cysteine) peptidase inhibitor, clade B, member 9b [Source:MGI Symbol;Acc:MGI:894668] | -2.22929 | 0. |
| ENSMUSG00000068855 | 319176 | *Hist2h2ac* | protein_coding | histone cluster 2, H2ac [Source:MGI Symbol;Acc:MGI:2448316] | -2.225914 | 1 |
| ENSMUSG00000094338 | 319185 | *Hist1h2bl* | protein_coding | histone cluster 1, H2bl [Source:MGI Symbol;Acc:MGI:2448403] | -2.2024 | 7 |
| ENSMUSG00000069305 | 319161 | *Hist1h4n* | protein_coding | histone cluster 1, H4n [Source:MGI Symbol;Acc:MGI:4843992] | -2.20155 | 0. |
| ENSMUSG00000065254 | NA | *Gm23973* | misc_RNA | predicted gene, 23973 [Source:MGI Symbol;Acc:MGI:5453750] | -2.185648 | 3 |
| ENSMUSG00000018102 | 68024 | *Hist1h2bc* | protein_coding | histone cluster 1, H2bc [Source:MGI Symbol;Acc:MGI:1915274] | -2.181749 | 3 |
| ENSMUSG00000069302 | 319168 | *Hist1h2ah* | protein_coding | histone cluster 1, H2ah [Source:MGI Symbol;Acc:MGI:2448295] | -2.174081 | 2 |
| ENSMUSG00000056270 | 109314 | *Prr9* | protein_coding | proline rich 9 [Source:MGI Symbol;Acc:MGI:1925680] | -2.157458 | 0. |
| ENSMUSG00000063767 | 381493 | *S100a7a* | protein_coding | S100 calcium binding protein A7A [Source:MGI Symbol;Acc:MGI:2687194] | -2.144488 | 0. |
| ENSMUSG00000086848 | 78382 | *Lce6a* | protein_coding | late cornified envelope 6A [Source:MGI Symbol;Acc:MGI:1925632] | -2.13625 | 4 |
| ENSMUSG00000093314 | NA | *Mir5136* | miRNA | microRNA 5136 [Source:MGI Symbol;Acc:MGI:4950461] | -2.104303 | 0. |
| ENSMUSG00000079597 | 433016 | *Gm5483* | protein_coding | predicted gene 5483 [Source:MGI Symbol;Acc:MGI:3645124] | -2.103801 | 0. |
| ENSMUSG00000048483 | 238331 | *Zdhhc22* | protein_coding | zinc finger, DHHC-type containing 22 [Source:MGI Symbol;Acc:MGI:2685108] | -2.097674 | 0. |
| ENSMUSG00000064288 | 319160 | *Hist1h4k* | protein_coding | histone cluster 1, H4k [Source:MGI Symbol;Acc:MGI:2448439] | -2.09539 | 1 |
| ENSMUSG00000074183 | 14857 | *Gsta1* | protein_coding | glutathione S-transferase, alpha 1 (Ya) [Source:MGI Symbol;Acc:MGI:1095417] | -2.086436 | 0. |
| ENSMUSG00000086801 | NA | *Gm15943* | antisense | predicted gene 15943 [Source:MGI Symbol;Acc:MGI:3802102] | -2.070638 | 0 |
| ENSMUSG00000075031 | 319178 | *Hist1h2bb* | protein_coding | histone cluster 1, H2bb [Source:MGI Symbol;Acc:MGI:2448377] | -2.054542 | 2 |
| ENSMUSG00000037953 | 333424 | *A4gnt* | protein_coding | alpha-1,4-N-acetylglucosaminyltransferase [Source:MGI Symbol;Acc:MGI:2143261] | -2.027703 | 0. |
| ENSMUSG00000037563 | 20055 | *Rps16* | protein_coding | ribosomal protein S16 [Source:MGI Symbol;Acc:MGI:98118] | -2.01195 | 8 |
| ENSMUSG00000064899 | NA | *Snord118* | snoRNA | small nucleolar RNA, C/D box 118 [Source:MGI Symbol;Acc:MGI:3819519] | -2.011581 | |
| ENSMUSG00000050063 | 19144 | *Klk6* | protein_coding | kallikrein related-peptidase 6 [Source:MGI Symbol;Acc:MGI:1343166] | -2.007204 | 0. |
| ENSMUSG00000008683 | 267019 | *Rps15a* | protein_coding | ribosomal protein S15A [Source:MGI Symbol;Acc:MGI:2389091] | -2.00257 | 4 |
| ENSMUSG00000095217 | 319187 | *Hist1h2bn* | protein_coding | histone cluster 1, H2bn [Source:MGI Symbol;Acc:MGI:2448407] | -2.00008 | 9 |
| ENSMUSG00000081838 | NA | *Gm13038* | processed_pseudogene | predicted gene 13038 [Source:MGI Symbol;Acc:MGI:3650684] | 2.005304 | 0. |
| ENSMUSG00000074217 | 70134 | *2210011C24Rik* | protein_coding | RIKEN cDNA 2210011C24 gene [Source:MGI Symbol;Acc:MGI:1917384] | 2.038815 | 0. |
| ENSMUSG00000075304 | 64406 | *Sp5* | protein_coding | trans-acting transcription factor 5 [Source:MGI Symbol;Acc:MGI:1927715] | 2.05191 | 0. |
| ENSMUSG00000072589 | NA | *Gm10371* | antisense | predicted gene 10371 [Source:MGI Symbol;Acc:MGI:3642716] | 2.05314 | 0. |
| ENSMUSG00000039683 | 330222 | *Sdk1* | protein_coding | sidekick cell adhesion molecule 1 [Source:MGI Symbol;Acc:MGI:2444413] | 2.177336 | 0. |
| ENSMUSG00000096349 | NA | *Gm22513* | snRNA | predicted gene, 22513 [Source:MGI Symbol;Acc:MGI:5452290] | 2.192986 | 0. |
| ENSMUSG00000116657 | NA | *Gm49774* | antisense | predicted gene, 49774 [Source:MGI Symbol;Acc:MGI:6215288] | 2.210757 | 0. |
| ENSMUSG00000095817 | 100041261 | *Gm3238* | protein_coding | predicted gene 3238 [Source:MGI Symbol;Acc:MGI:3781416] | 2.220815 | 0. |
| ENSMUSG00000117465 | NA | *AC102496.1* | lincRNA | novel transcript | 2.271198 | 0. |
| ENSMUSG00000097983 | NA | *Gm26971* | sense_intronic | predicted gene, 26971 [Source:MGI Symbol;Acc:MGI:5504086] | 2.312075 | 0. |
| ENSMUSG00000090955 | NA | *Gm17097* | processed_pseudogene | predicted gene 17097 [Source:MGI Symbol;Acc:MGI:4937924] | 2.31641 | 0. |
| ENSMUSG00000104040 | NA | *Gm37563* | TEC | predicted gene, 37563 [Source:MGI Symbol;Acc:MGI:5610791] | 2.368809 | 0. |
| ENSMUSG00000107559 | NA | *Gm44193* | TEC | predicted gene, 44193 [Source:MGI Symbol;Acc:MGI:5690585] | 2.375337 | 0. |
| ENSMUSG00000002012 | 93843 | *Pnck* | protein_coding | pregnancy upregulated non-ubiquitously expressed CaM kinase [Source:MGI Symbol;Acc:MGI:1347357] | 2.408702 | 0. |
| ENSMUSG00000096421 | 100502953 | *Gm10100* | protein_coding | predicted gene 10100 [Source:MGI Symbol;Acc:MGI:3642388] | 2.412681 | 0. |
| ENSMUSG00000098678 | 223645 | *Mroh6* | protein_coding | maestro heat-like repeat family member 6 [Source:MGI Symbol;Acc:MGI:5011755] | 2.491765 | 0 |
| ENSMUSG00000104696 | NA | *Gm42946* | TEC | predicted gene 42946 [Source:MGI Symbol;Acc:MGI:5663083] | 2.723052 | 0. |
| ENSMUSG00000084897 | 50518 | *Gm14226* | protein_coding | predicted gene 14226 [Source:MGI Symbol;Acc:MGI:3649244] | 2.729843 | 0 |
| ENSMUSG00000022957 | 16443 | *Itsn1* | protein_coding | intersectin 1 (SH3 domain protein 1A) [Source:MGI Symbol;Acc:MGI:1338069] | 2.829821 | 2 |
| ENSMUSG00000094841 | NA | *Gm10610* | lincRNA | predicted gene 10610 [Source:MGI Symbol;Acc:MGI:3642045] | 2.923944 | 0. |
| ENSMUSG00000081303 | NA | *Gm16011* | processed_pseudogene | predicted gene 16011 [Source:MGI Symbol;Acc:MGI:3801796] | 3.217223 | 0. |
| ENSMUSG00000104232 | NA | *Gm37590* | TEC | predicted gene, 37590 [Source:MGI Symbol;Acc:MGI:5610818] | 3.26393 | 0. |

| | | | | | | |
|---|---|---|---|---|---|---|
| ENSMUSG00000018263 | 21388 | *Tbx5* | protein_coding | T-box 5 [Source:MGI Symbol;Acc:MGI:102541] | 3.285104 | 0.[illegible] |
| ENSMUSG00000069584 | NA | *Gm10272* | protein_coding | predicted gene 10272 [Source:MGI Symbol;Acc:MGI:3642183] | 3.311198 | 0.[illegible] |
| ENSMUSG00000100510 | NA | *Hand2os1* | processed_transcript | Hand2, opposite strand 1 [Source:MGI Symbol;Acc:MGI:5578769] | 3.571063 | 0.[illegible] |
| ENSMUSG00000055197 | 260298 | *Fev* | protein_coding | FEV (ETS oncogene family) [Source:MGI Symbol;Acc:MGI:2449712] | 4.157327 | 0.[illegible] |
| ENSMUSG00000026347 | 72160 | *Tmem163* | protein_coding | transmembrane protein 163 [Source:MGI Symbol;Acc:MGI:1919410] | 4.281242 | 2[illegible] |
| ENSMUSG00000115702 | NA | *Gm5206* | transcribed_processed_pseudogene | predicted pseudogene 5206 [Source:MGI Symbol;Acc:MGI:3645516] | 4.991942 | 5[illegible] |
| ENSMUSG00000091383 | NA | *Hist1h2al* | processed_pseudogene | histone cluster 1, H2al [Source:MGI Symbol;Acc:MGI:3646032] | 6.466509 | 1[illegible] |
| ENSMUSG00000059751 | NA | *Rps3a3* | processed_pseudogene | ribosomal protein S3A3 [Source:MGI Symbol;Acc:MGI:3643406] | 6.86999 | 2[illegible] |
