## SupplementalTable10 for "An enhancer:involucrin regulatory module impacts human skin barrier adaptation out-of-Africa and modifies atopic dermatitis risk"

**Table S10. Ranked list of differentially expressed genes between Ivl -/- and WT mice whole skin from RNA-seq.**

| *Feature_ID* | *entrezgene* | *external_gene_name* | *gene_biotype* | *description* | *logFC* | *adj.P.Val* |
|---|---|---|---|---|---|---|
| ENSMUSG00000069306 | 100041230 | *Hist1h4m* | protein_coding | histone cluster 1, H4m [Source:MGI Symbol;Acc:MGI:2448441] | -11.114 | 0.00164645 |
| ENSMUSG00000049128 | 16447 | *Ivl* | protein_coding | involucrin [Source:MGI Symbol;Acc:MGI:96626] | -7.674693 | 1.96E-1 |
| ENSMUSG00000086324 | NA | *Gm15564* | antisense | predicted gene 15564 [Source:MGI Symbol;Acc:MGI:3783013] | -5.179489 | 0.00415718 |
| ENSMUSG00000057174 | 170939 | *Krtap19-9b* | protein_coding | keratin associated protein 19-9B [Source:MGI Symbol;Acc:MGI:2181750] | -5.004569 | 0.00066712 |
| ENSMUSG00000092909 | NA | *Gm25732* | miRNA | predicted gene, 25732 [Source:MGI Symbol;Acc:MGI:5455509] | -4.925226 | 0.0272112 |
| ENSMUSG00000028081 | 20091 | *Rps3a1* | protein_coding | ribosomal protein S3A1 [Source:MGI Symbol;Acc:MGI:1202063] | -4.882306 | 5.70E-1 |
| ENSMUSG00000068885 | 69520 | *Lce3f* | protein_coding | late cornified envelope 3F [Source:MGI Symbol;Acc:MGI:1916770] | -4.647524 | 0.0039097 |
| ENSMUSG00000032080 | 11808 | *Apoa4* | protein_coding | apolipoprotein A-IV [Source:MGI Symbol;Acc:MGI:88051] | -4.485311 | 0.0152514 |
| ENSMUSG00000022931 | 26560 | *Krtap15* | protein_coding | keratin associated protein 15 [Source:MGI Symbol;Acc:MGI:1347350] | -4.327468 | 0.00395693 |
| ENSMUSG00000074928 | 23927 | *Krtap14* | protein_coding | keratin associated protein 14 [Source:MGI Symbol;Acc:MGI:1346079] | -4.283546 | 0.00842211 |
| ENSMUSG00000049972 | 329918 | *Skint9* | protein_coding | selection and upkeep of intraepithelial T cells 9 [Source:MGI Symbol;Acc:MGI:3045341] | -4.260082 | 0.00060955 |
| ENSMUSG00000027908 | 71325 | *Tchhl1* | protein_coding | trichohyalin-like 1 [Source:MGI Symbol;Acc:MGI:1918575] | -4.212048 | 1.19E-0 |
| ENSMUSG00000070868 | 195564 | *Skint3* | protein_coding | selection and upkeep of intraepithelial T cells 3 [Source:MGI Symbol;Acc:MGI:3045331] | -4.165558 | 0.00063026 |
| ENSMUSG00000055960 | 320640 | *Skint4* | protein_coding | selection and upkeep of intraepithelial T cells 4 [Source:MGI Symbol;Acc:MGI:2444425] | -4.078137 | 0.00046739 |
| ENSMUSG00000032083 | 11806 | *Apoa1* | protein_coding | apolipoprotein A-I [Source:MGI Symbol;Acc:MGI:88049] | -4.030253 | 0.0377021 |
| ENSMUSG00000096534 | 71369 | *Krtap16-3* | protein_coding | keratin associated protein 16-3 [Source:MGI Symbol;Acc:MGI:1918619] | -3.886214 | 0.0015339 |
| ENSMUSG00000079013 | 238395 | *Serpina3j* | protein_coding | serine (or cysteine) peptidase inhibitor, clade A (alpha-1 antiproteinase, antitrypsin), member 3J [Source:MGI Symbol;Acc:MGI:2182843] | -3.872281 | 0.00112861 |
| ENSMUSG00000038567 | 13081 | *Cyp24a1* | protein_coding | cytochrome P450, family 24, subfamily a, polypeptide 1 [Source:MGI Symbol;Acc:MGI:88593] | -3.8445 | 0.00178060 |
| ENSMUSG00000060469 | 77918 | *Krtap19-3* | protein_coding | keratin associated protein 19-3 [Source:MGI Symbol;Acc:MGI:1925168] | -3.745647 | 0.010719 |
| ENSMUSG00000082699 | NA | *Gm12736* | processed_pseudogene | predicted gene 12736 [Source:MGI Symbol;Acc:MGI:3650723] | -3.7071 | 0.0002653 |
| ENSMUSG00000062400 | 68484 | *Krtap6-5* | protein_coding | keratin associated protein 6-5 [Source:MGI Symbol;Acc:MGI:1915734] | -3.639797 | 0.00159488 |
| ENSMUSG00000017300 | 21925 | *Tnnc2* | protein_coding | troponin C2, fast [Source:MGI Symbol;Acc:MGI:98780] | -3.636399 | 1.32E-0 |
| ENSMUSG00000058368 | 170656 | *Krtap21-1* | protein_coding | keratin associated protein 21-1 [Source:MGI Symbol;Acc:MGI:2157767] | -3.563505 | 0.00110427 |
| ENSMUSG00000069372 | 629147 | *Ctxn3* | protein_coding | cortexin 3 [Source:MGI Symbol;Acc:MGI:3642816] | -3.527552 | 0.0482299 |
| ENSMUSG00000035202 | 102436 | *Lars2* | protein_coding | leucyl-tRNA synthetase, mitochondrial [Source:MGI Symbol;Acc:MGI:2142973] | -3.517735 | 0.00901382 |
| ENSMUSG00000058172 | 16700 | *Krtap6-1* | protein_coding | keratin associated protein 6-1 [Source:MGI Symbol;Acc:MGI:1330228] | -3.486279 | 0.00166418 |
| ENSMUSG00000098973 | NA | *Mir6236* | miRNA | microRNA 6236 [Source:MGI Symbol;Acc:MGI:5530929] | -3.40162 | 0.0200108 |
| ENSMUSG00000095992 | 68740 | *Krtap22-2* | protein_coding | keratin associated protein 22-2 [Source:MGI Symbol;Acc:MGI:1915990] | -3.369604 | 0.00070083 |
| ENSMUSG00000087194 | 230622 | *Skint6* | protein_coding | selection and upkeep of intraepithelial T cells 6 [Source:MGI Symbol;Acc:MGI:3649262] | -3.303689 | 0.00055862 |
| ENSMUSG00000033948 | 74464 | *Zswim5* | protein_coding | zinc finger SWIM-type containing 5 [Source:MGI Symbol;Acc:MGI:1921714] | -3.267198 | 0.0311364 |
| ENSMUSG00000082678 | NA | *Gm12818* | processed_pseudogene | predicted gene 12818 [Source:MGI Symbol;Acc:MGI:3649634] | -3.2244 | 0.00459194 |
| ENSMUSG00000040852 | 213556 | *Plekhh2* | protein_coding | pleckstrin homology domain containing, family H (with MyTH4 domain) member 2 [Source:MGI Symbol;Acc:MGI:2146813] | -3.198944 | 0.00730020 |
| ENSMUSG00000028834 | 433766 | *Trim63* | protein_coding | tripartite motif-containing 63 [Source:MGI Symbol;Acc:MGI:2447992] | -3.188662 | 0.0187187 |
| ENSMUSG00000045566 | 229562 | *Sprr4* | protein_coding | small proline-rich protein 4 [Source:MGI Symbol;Acc:MGI:2654508] | -3.149035 | 0.00065859 |
| ENSMUSG00000068075 | 100040201 | *Gm10229* | protein_coding | predicted gene 10229 [Source:MGI Symbol;Acc:MGI:3711943] | -3.120352 | 0.00098679 |
| ENSMUSG00000091573 | NA | *Serpina3d-ps* | unprocessed_pseudogene | serine (or cysteine) peptidase inhibitor, clade A, member 3D, pseudogene [Source:MGI Symbol;Acc:MGI:2182836] | -3.118783 | 0.0148988 |
| ENSMUSG00000024411 | 11829 | *Aqp4* | protein_coding | aquaporin 4 [Source:MGI Symbol;Acc:MGI:107387] | -3.101655 | 0.00451522 |
| ENSMUSG00000065037 | 19817 | *Rn7sk* | misc_RNA | RNA, 7SK, nuclear [Source:MGI Symbol;Acc:MGI:103186] | -3.083254 | 2.24E-1 |
| ENSMUSG00000023078 | 55985 | *Cxcl13* | protein_coding | chemokine (C-X-C motif) ligand 13 [Source:MGI Symbol;Acc:MGI:1888499] | -3.012573 | 0.00015769 |
| ENSMUSG00000106174 | NA | *Gm43235* | unprocessed_pseudogene | predicted gene 43235 [Source:MGI Symbol;Acc:MGI:5663372] | -2.982233 | 0.039440 |
| ENSMUSG00000093218 | NA | *Gm25252* | miRNA | predicted gene, 25252 [Source:MGI Symbol;Acc:MGI:5455029] | -2.922841 | 0.00019577 |
| ENSMUSG00000062433 | 16701 | *Krtap6-2* | protein_coding | keratin associated protein 6-2 [Source:MGI Symbol;Acc:MGI:1330280] | -2.871519 | 0.0114425 |
| ENSMUSG00000023267 | 14409 | *Gabrr2* | protein_coding | gamma-aminobutyric acid (GABA) C receptor, subunit rho 2 [Source:MGI Symbol;Acc:MGI:95626] | -2.85728 | 0.0132710 |
| ENSMUSG00000060678 | 319155 | *Hist1h4c* | protein_coding | histone cluster 1, H4c [Source:MGI Symbol;Acc:MGI:2448421] | -2.851275 | 4.63E-1 |
| ENSMUSG00000049809 | 75586 | *Krtap9-3* | protein_coding | keratin associated protein 9-3 [Source:MGI Symbol;Acc:MGI:1922836] | -2.796016 | 0.0142539 |
| ENSMUSG00000045515 | 18993 | *Pou3f3* | protein_coding | POU domain, class 3, transcription factor 3 [Source:MGI Symbol;Acc:MGI:102564] | -2.788303 | 0.0167244 |
| ENSMUSG00000060560 | 234677 | *Ces4a* | protein_coding | carboxylesterase 4A [Source:MGI Symbol;Acc:MGI:2384581] | -2.769375 | 0.00119194 |
| ENSMUSG00000086298 | NA | *Gm11716* | antisense | predicted gene 11716 [Source:MGI Symbol;Acc:MGI:3649215] | -2.763502 | 0.00020474 |
| ENSMUSG00000057977 | 230623 | *Skint11* | protein_coding | selection and upkeep of intraepithelial T cells 11 [Source:MGI Symbol;Acc:MGI:2685415] | -2.739083 | 0.00218199 |
| ENSMUSG00000042985 | 100647 | *Upk3b* | protein_coding | uroplakin 3B [Source:MGI Symbol;Acc:MGI:2140882] | -2.716239 | 0.00319035 |
| ENSMUSG00000055298 | 26898 | *Ctsj* | protein_coding | cathepsin J [Source:MGI Symbol;Acc:MGI:1349426] | -2.715879 | 0.00410616 |
| ENSMUSG00000060981 | 69386 | *Hist1h4h* | protein_coding | histone cluster 1, H4h [Source:MGI Symbol;Acc:MGI:2448427] | -2.688193 | 3.59E-1 |
| ENSMUSG00000057322 | 67671 | *Rpl38* | protein_coding | ribosomal protein L38 [Source:MGI Symbol;Acc:MGI:1914921] | -2.683652 | 3.72E-1 |
| ENSMUSG00000059632 | 16703 | *Krtap8-1* | protein_coding | keratin associated protein 8-1 [Source:MGI Symbol;Acc:MGI:1330293] | -2.661073 | 0.00330994 |
| ENSMUSG00000090527 | 433597 | *Gm5538* | protein_coding | predicted gene 5538 [Source:MGI Symbol;Acc:MGI:3779495] | -2.607808 | 0.00367359 |
| ENSMUSG00000074433 | 69514 | *Lce3e* | protein_coding | late cornified envelope 3E [Source:MGI Symbol;Acc:MGI:1916764] | -2.60615 | 0.0114425 |
| ENSMUSG00000024846 | 73720 | *Cst6* | protein_coding | cystatin E/M [Source:MGI Symbol;Acc:MGI:1920970] | -2.594556 | 7.32E-0 |
| ENSMUSG00000042045 | 66402 | *Sln* | protein_coding | sarcolipin [Source:MGI Symbol;Acc:MGI:1913652] | -2.575175 | 0.00170649 |
| ENSMUSG00000052180 | 97848 | *Serpinb6c* | protein_coding | serine (or cysteine) peptidase inhibitor, clade B, member 6c [Source:MGI Symbol;Acc:MGI:2145481] | -2.556487 | 0.00043318 |
| ENSMUSG00000099842 | NA | *Gm4034* | processed_pseudogene | predicted gene 4034 [Source:MGI Symbol;Acc:MGI:3782208] | -2.531045 | 1.09E-0 |
| ENSMUSG00000081169 | NA | *Gm12551* | unprocessed_pseudogene | predicted gene 12551 [Source:MGI Symbol;Acc:MGI:3651664] | -2.518646 | 0.00160891 |
| ENSMUSG00000063767 | 381493 | *S100a7a* | protein_coding | S100 calcium binding protein A7A [Source:MGI Symbol;Acc:MGI:2687194] | -2.511191 | 0.00028292 |
| ENSMUSG00000005716 | 19293 | *Pvalb* | protein_coding | parvalbumin [Source:MGI Symbol;Acc:MGI:97821] | -2.497871 | 0.0253554 |
| ENSMUSG00000089948 | NA | *Far2os1* | antisense | fatty acyl CoA reductase 2, opposite strand 1 [Source:MGI Symbol;Acc:MGI:4415003] | -2.497209 | 0.030818 |
| ENSMUSG00000064347 | NA | *mt-Ta* | Mt_tRNA | mitochondrially encoded tRNA alanine [Source:MGI Symbol;Acc:MGI:102491] | -2.434675 | 7.53E-0 |
| ENSMUSG00000093674 | 67945 | *Rpl41* | protein_coding | ribosomal protein L41 [Source:MGI Symbol;Acc:MGI:1915195] | -2.432223 | 1.66E-1 |
| ENSMUSG00000053719 | 16618 | *Klk1b26* | protein_coding | kallikrein 1-related petidase b26 [Source:MGI Symbol;Acc:MGI:891981] | -2.431491 | 0.0181499 |
| ENSMUSG00000038670 | 233199 | *Mybpc2* | protein_coding | myosin binding protein C, fast-type [Source:MGI Symbol;Acc:MGI:1336170] | -2.4145 | 0.016370 |
| ENSMUSG00000114279 | 319186 | *Hist1h2bm* | protein_coding | histone cluster 1, H2bm [Source:MGI Symbol;Acc:MGI:2448404] | -2.407702 | 8.74E-0 |
| ENSMUSG00000066364 | 271047 | *Serpina3b* | protein_coding | serine (or cysteine) peptidase inhibitor, clade A, member 3B [Source:MGI Symbol;Acc:MGI:2182835] | -2.402826 | 0.00015859 |
| ENSMUSG00000021403 | 20706 | *Serpinb9b* | protein_coding | serine (or cysteine) peptidase inhibitor, clade B, member 9b [Source:MGI Symbol;Acc:MGI:894668] | -2.389597 | 0.00186218 |
| ENSMUSG00000030237 | 28250 | *Slco1a4* | protein_coding | solute carrier organic anion transporter family, member 1a4 [Source:MGI Symbol;Acc:MGI:1351896] | -2.388574 | 0.0208339 |
| ENSMUSG00000069718 | 100040248 | *Gm11563* | protein_coding | predicted gene 11563 [Source:MGI Symbol;Acc:MGI:3650330] | -2.38387 | 0.0321563 |
| ENSMUSG00000075031 | 319178 | *Hist1h2bb* | protein_coding | histone cluster 1, H2bb [Source:MGI Symbol;Acc:MGI:2448377] | -2.380083 | 4.92E-0 |
| ENSMUSG00000069792 | 100034251 | *Wfdc17* | protein_coding | WAP four-disulfide core domain 17 [Source:MGI Symbol;Acc:MGI:3649773] | -2.35702 | 0.00831589 |
| ENSMUSG00000037563 | 20055 | *Rps16* | protein_coding | ribosomal protein S16 [Source:MGI Symbol;Acc:MGI:98118] | -2.345048 | 2.51E-1 |
| ENSMUSG00000030785 | 12862 | *Cox6a2* | protein_coding | cytochrome c oxidase subunit 6A2 [Source:MGI Symbol;Acc:MGI:104649] | -2.341498 | 0.00135547 |
| ENSMUSG00000043219 | 15403 | *Hoxa6* | protein_coding | homeobox A6 [Source:MGI Symbol;Acc:MGI:96178] | -2.340478 | 0.0126634 |
| ENSMUSG00000042212 | 20758 | *Sprr2d* | protein_coding | small proline-rich protein 2D [Source:MGI Symbol;Acc:MGI:1330347] | -2.329071 | 0.0116360 |
| ENSMUSG00000082766 | NA | *1700064H15Rik* | protein_coding | RIKEN cDNA 1700064H15 gene [Source:MGI Symbol;Acc:MGI:1920674] | -2.326279 | 3.66E-0 |
| ENSMUSG00000026390 | 17167 | *Marco* | protein_coding | macrophage receptor with collagenous structure [Source:MGI Symbol;Acc:MGI:1309998] | -2.324156 | 0.0354361 |
| ENSMUSG00000062727 | 319184 | *Hist1h2bk* | protein_coding | histone cluster 1, H2bk [Source:MGI Symbol;Acc:MGI:2448399] | -2.323362 | 8.12E-0 |
| ENSMUSG00000103084 | NA | *Gm38119* | protein_coding | predicted gene, 38119 [Source:MGI Symbol;Acc:MGI:5611347] | -2.321923 | 0.00643862 |
| ENSMUSG00000056054 | 20201 | *S100a8* | protein_coding | S100 calcium binding protein A8 (calgranulin A) [Source:MGI Symbol;Acc:MGI:88244] | -2.320829 | 0.00294415 |
| ENSMUSG00000068855 | 319176 | *Hist2h2ac* | protein_coding | histone cluster 2, H2ac [Source:MGI Symbol;Acc:MGI:2448316] | -2.319494 | 1.44E-1 |
| ENSMUSG00000112188 | NA | *Gm47708* | lincRNA | predicted gene, 47708 [Source:MGI Symbol;Acc:MGI:6096827] | -2.317008 | 0.0031876 |
| ENSMUSG00000031097 | 21953 | *Tnni2* | protein_coding | troponin I, skeletal, fast 2 [Source:MGI Symbol;Acc:MGI:105070] | -2.316066 | 0.00447574 |
| ENSMUSG00000021456 | 14120 | *Fbp2* | protein_coding | fructose bisphosphatase 2 [Source:MGI Symbol;Acc:MGI:95491] | -2.312927 | 0.00112025 |
| ENSMUSG00000067455 | 319159 | *Hist1h4j* | protein_coding | histone cluster 1, H4j [Source:MGI Symbol;Acc:MGI:2448436] | -2.309552 | 1.06E-0 |
| ENSMUSG00000080885 | NA | *Rpl10-ps6* | processed_pseudogene | ribosomal protein L10, pseudogene 6 [Source:MGI Symbol;Acc:MGI:3782343] | -2.307117 | 9.60E-0 |
| ENSMUSG00000047253 | 69664 | *Krtap1-5* | protein_coding | keratin associated protein 1-5 [Source:MGI Symbol;Acc:MGI:1916914] | -2.305145 | 0.0157685 |
| ENSMUSG00000092746 | NA | *Rn7s6* | misc_RNA | 7S RNA 6 [Source:MGI Symbol;Acc:MGI:97956] | -2.296659 | 7.71E-0 |
| ENSMUSG00000062077 | 58522 | *Trim54* | protein_coding | tripartite motif-containing 54 [Source:MGI Symbol;Acc:MGI:1889623] | -2.293314 | 0.00059540 |
| ENSMUSG00000056999 | 15925 | *Ide* | protein_coding | insulin degrading enzyme [Source:MGI Symbol;Acc:MGI:96412] | -2.284985 | 2.29E-0 |
| ENSMUSG00000056706 | 71363 | *Krtap7-1* | protein_coding | keratin associated protein 7-1 [Source:MGI Symbol;Acc:MGI:1918613] | -2.28395 | 0.011423 |
| ENSMUSG00000031779 | 20299 | *Ccl22* | protein_coding | chemokine (C-C motif) ligand 22 [Source:MGI Symbol;Acc:MGI:1306779] | -2.283168 | 0.00371238 |
| ENSMUSG00000096010 | 320332 | *Hist4h4* | protein_coding | histone cluster 4, H4 [Source:MGI Symbol;Acc:MGI:2448443] | -2.277637 | 3.59E-1 |
| ENSMUSG00000052819 | 212989 | *Best2* | protein_coding | bestrophin 2 [Source:MGI Symbol;Acc:MGI:2387588] | -2.25215 | 0.0273016 |
| ENSMUSG00000038236 | 15404 | *Hoxa7* | protein_coding | homeobox A7 [Source:MGI Symbol;Acc:MGI:96179] | -2.242497 | 0.0121759 |

| | | | | | | |
|---|---|---|---|---|---|---|
| ENSMUSG00000015665 | 245533 | *Awat1* | protein_coding | acyl-CoA wax alcohol acyltransferase 1 [Source:MGI Symbol;Acc:MGI:3588200] | -2.240769 | 0.00060021 |
| ENSMUSG00000030672 | 17907 | *Mylpf* | protein_coding | myosin light chain, phosphorylatable, fast skeletal muscle [Source:MGI Symbol;Acc:MGI:97273] | -2.232251 | 0.00398243 |
| ENSMUSG00000037953 | 333424 | *A4gnt* | protein_coding | alpha-1,4-N-acetylglucosaminyltransferase [Source:MGI Symbol;Acc:MGI:2143261] | -2.225887 | 6.20E-0 |
| ENSMUSG00000031382 | 68854 | *Asb11* | protein_coding | ankyrin repeat and SOCS box-containing 11 [Source:MGI Symbol;Acc:MGI:1916104] | -2.219595 | 0.00882161 |
| ENSMUSG00000100190 | NA | *Krtap28-10* | protein_coding | keratin associated protein 28-10 [Source:MGI Symbol;Acc:MGI:3779575] | -2.216962 | 0.0486597 |
| ENSMUSG00000100511 | 74305 | *1700111N16Rik* | antisense | RIKEN cDNA 1700111N16 gene [Source:MGI Symbol;Acc:MGI:1921555] | -2.206489 | 0.00136372 |
| ENSMUSG00000077391 | NA | *Gm24336* | snoRNA | predicted gene, 24336 [Source:MGI Symbol;Acc:MGI:5454113] | -2.201206 | 2.27E-0 |
| ENSMUSG00000092837 | NA | *Rpph1* | ribozyme | ribonuclease P RNA component H1 [Source:MGI Symbol;Acc:MGI:1934664] | -2.19566 | 6.51E-0 |
| ENSMUSG00000068877 | 20342 | *Selenbp2* | protein_coding | selenium binding protein 2 [Source:MGI Symbol;Acc:MGI:104859] | -2.192436 | 0.00106876 |
| ENSMUSG00000087968 | NA | *Gm25395* | scaRNA | predicted gene, 25395 [Source:MGI Symbol;Acc:MGI:5455172] | -2.189961 | 1.30E-0 |
| ENSMUSG00000107451 | NA | *Gm44421* | lincRNA | predicted gene, 44421 [Source:MGI Symbol;Acc:MGI:5690813] | -2.16974 | 0.00734905 |
| ENSMUSG00000061482 | 319156 | *Hist1h4d* | protein_coding | histone cluster 1, H4d [Source:MGI Symbol;Acc:MGI:2448423] | -2.161726 | 6.72E-0 |
| ENSMUSG00000078131 | 435273 | *Krtap1-3* | protein_coding | keratin associated protein 1-3 [Source:MGI Symbol;Acc:MGI:3650443] | -2.15899 | 0.0383477 |
| ENSMUSG00000020475 | 56012 | *Pgam2* | protein_coding | phosphoglycerate mutase 2 [Source:MGI Symbol;Acc:MGI:1933118] | -2.157761 | 0.00183229 |
| ENSMUSG00000068074 | 100040214 | *Gm10228* | protein_coding | predicted gene 10228 [Source:MGI Symbol;Acc:MGI:3704467] | -2.156763 | 0.0231990 |
| ENSMUSG00000001773 | 53320 | *Folh1* | protein_coding | folate hydrolase 1 [Source:MGI Symbol;Acc:MGI:1858193] | -2.147778 | 0.00424968 |
| ENSMUSG00000023153 | 69671 | *Tmem52* | protein_coding | transmembrane protein 52 [Source:MGI Symbol;Acc:MGI:1916921] | -2.136695 | 0.0144206 |
| ENSMUSG00000069722 | 66380 | *Krtap3-3* | protein_coding | keratin associated protein 3-3 [Source:MGI Symbol;Acc:MGI:1913630] | -2.121211 | 0.0137142 |
| ENSMUSG00000030399 | 12715 | *Ckm* | protein_coding | creatine kinase, muscle [Source:MGI Symbol;Acc:MGI:88413] | -2.110767 | 0.0109697 |
| ENSMUSG00000064288 | 319160 | *Hist1h4k* | protein_coding | histone cluster 1, H4k [Source:MGI Symbol;Acc:MGI:2448439] | -2.106847 | 8.73E-0 |
| ENSMUSG00000109973 | NA | *Gm45397* | lincRNA | predicted gene 45397 [Source:MGI Symbol;Acc:MGI:5791233] | -2.103577 | 0.00028943 |
| ENSMUSG00000111912 | NA | *Gm48521* | lincRNA | predicted gene, 48521 [Source:MGI Symbol;Acc:MGI:6098057] | -2.09826 | 0.00090550 |
| ENSMUSG00000026985 | 69677 | *Il1f8* | protein_coding | interleukin 1 family, member 8 [Source:MGI Symbol;Acc:MGI:1916927] | -2.096863 | 1.42E-0 |
| ENSMUSG00000069267 | 319150 | *Hist1h3b* | protein_coding | histone cluster 1, H3b [Source:MGI Symbol;Acc:MGI:2448319] | -2.090672 | 1.17E-0 |
| ENSMUSG00000050063 | 19144 | *Klk6* | protein_coding | kallikrein related-peptidase 6 [Source:MGI Symbol;Acc:MGI:1343166] | -2.090533 | 0.00029306 |
| ENSMUSG00000051748 | 66107 | *Wfdc21* | protein_coding | WAP four-disulfide core domain 21 [Source:MGI Symbol;Acc:MGI:1913357] | -2.086112 | 7.89E-0 |
| ENSMUSG00000056328 | 17879 | *Myh1* | protein_coding | myosin, heavy polypeptide 1, skeletal muscle, adult [Source:MGI Symbol;Acc:MGI:1339711] | -2.085837 | 0.00842991 |
| ENSMUSG00000007877 | 21393 | *Tcap* | protein_coding | titin-cap [Source:MGI Symbol;Acc:MGI:1330233] | -2.083346 | 0.00836103 |
| ENSMUSG00000079017 | 76933 | *Ifi27l2a* | protein_coding | interferon, alpha-inducible protein 27 like 2A [Source:MGI Symbol;Acc:MGI:1924183] | -2.079273 | 4.78E-0 |
| ENSMUSG00000044748 | 13214 | *Defb1* | protein_coding | defensin beta 1 [Source:MGI Symbol;Acc:MGI:1096878] | -2.076503 | 0.00011978 |
| ENSMUSG00000042254 | 214425 | *Cilp* | protein_coding | cartilage intermediate layer protein, nucleotide pyrophosphohydrolase [Source:MGI Symbol;Acc:MGI:2444507] | -2.069957 | 0.00054630 |
| ENSMUSG00000089281 | NA | *Scarna6* | scaRNA | small Cajal body-specific RNA 6 [Source:MGI Symbol;Acc:MGI:3819487] | -2.057776 | 9.56E-0 |
| ENSMUSG00000019933 | 69563 | *Mrln* | protein_coding | myoregulin [Source:MGI Symbol;Acc:MGI:1916813] | -2.055473 | 0.00093973 |
| ENSMUSG00000030996 | 11870 | *Art1* | protein_coding | ADP-ribosyltransferase 1 [Source:MGI Symbol;Acc:MGI:107511] | -2.050561 | 0.00645593 |
| ENSMUSG00000062694 | 12391 | *Cav3* | protein_coding | caveolin 3 [Source:MGI Symbol;Acc:MGI:107570] | -2.046603 | 0.00114301 |
| ENSMUSG00000067288 | 54127 | *Rps28* | protein_coding | ribosomal protein S28 [Source:MGI Symbol;Acc:MGI:1859516] | -2.042956 | 9.26E-1 |
| ENSMUSG00000096965 | 78512 | *3300005D01Rik* | lincRNA | RIKEN cDNA 3300005D01 gene [Source:MGI Symbol;Acc:MGI:1925762] | -2.042858 | 0.00017204 |
| ENSMUSG00000069274 | 319157 | *Hist1h4f* | protein_coding | histone cluster 1, H4f [Source:MGI Symbol;Acc:MGI:2448425] | -2.036614 | 1.83E-0 |
| ENSMUSG00000006587 | 30927 | *Snai3* | protein_coding | snail family zinc finger 3 [Source:MGI Symbol;Acc:MGI:1353563] | -2.034214 | 0.0253424 |
| ENSMUSG00000069309 | 319170 | *Hist1h2an* | protein_coding | histone cluster 1, H2an [Source:MGI Symbol;Acc:MGI:2448300] | -2.029664 | 2.68E-0 |
| ENSMUSG00000087881 | NA | *Gm22442* | scaRNA | predicted gene, 22442 [Source:MGI Symbol;Acc:MGI:5452219] | -2.027412 | 8.26E-1 |
| ENSMUSG00000047246 | 319179 | *Hist1h2be* | protein_coding | histone cluster 1, H2be [Source:MGI Symbol;Acc:MGI:2448380] | -2.027286 | 1.16E-0 |
| ENSMUSG00000041841 | 100502825 | *Rpl37* | protein_coding | ribosomal protein L37 [Source:MGI Symbol;Acc:MGI:1914531] | -2.023278 | 3.79E-1 |
| ENSMUSG00000041841 | 67281 | *Rpl37* | protein_coding | ribosomal protein L37 [Source:MGI Symbol;Acc:MGI:1914531] | -2.023278 | 3.79E-1 |
| ENSMUSG00000022215 | 68680 | *Fitm1* | protein_coding | fat storage-inducing transmembrane protein 1 [Source:MGI Symbol;Acc:MGI:1915930] | -2.016988 | 0.00313641 |
| ENSMUSG00000006457 | 11474 | *Actn3* | protein_coding | actinin alpha 3 [Source:MGI Symbol;Acc:MGI:99678] | -2.013423 | 0.0171623 |
| ENSMUSG00000043681 | 69134 | *Fam25c* | protein_coding | family with sequence similarity 25, member C [Source:MGI Symbol;Acc:MGI:1916384] | -2.007213 | 6.75E-0 |
| ENSMUSG00000069266 | 326620 | *Hist1h4b* | protein_coding | histone cluster 1, H4b [Source:MGI Symbol;Acc:MGI:2448420] | -2.002982 | 6.72E-0 |
| ENSMUSG00000043753 | 242523 | *Dmrta1* | protein_coding | doublesex and mab-3 related transcription factor like family A1 [Source:MGI Symbol;Acc:MGI:2653627] | -2.00125 | 0.00015371 |
| ENSMUSG00000048483 | 238331 | *Zdhhc22* | protein_coding | zinc finger, DHHC-type containing 22 [Source:MGI Symbol;Acc:MGI:2685108] | -2.00045 | 9.49E-0 |
| ENSMUSG00000081142 | NA | *Gm15497* | processed_pseudogene | predicted gene 15497 [Source:MGI Symbol;Acc:MGI:3782944] | 2.001126 | 0.00078816 |
| ENSMUSG00000079654 | 101359 | *Prrt4* | protein_coding | proline-rich transmembrane protein 4 [Source:MGI Symbol;Acc:MGI:2141677] | 2.003046 | 0.00269412 |
| ENSMUSG00000090255 | 66737 | *4921534H16Rik* | antisense | RIKEN cDNA 4921534H16 gene [Source:MGI Symbol;Acc:MGI:1913987] | 2.005352 | 0.0019161 |
| ENSMUSG00000097983 | NA | *Gm26971* | sense_intronic | predicted gene, 26971 [Source:MGI Symbol;Acc:MGI:5504086] | 2.008283 | 0.00830269 |
| ENSMUSG00000102160 | NA | *Gm36944* | TEC | predicted gene, 36944 [Source:MGI Symbol;Acc:MGI:5610172] | 2.013043 | 0.00390445 |
| ENSMUSG00000069583 | 16694 | *Krtap12-1* | protein_coding | keratin associated protein 12-1 [Source:MGI Symbol;Acc:MGI:1328315] | 2.020037 | 0.00589742 |
| ENSMUSG00000083382 | NA | *Gm6433* | processed_pseudogene | predicted gene 6433 [Source:MGI Symbol;Acc:MGI:3645394] | 2.024659 | 0.00969881 |
| ENSMUSG00000097520 | NA | *4930488L21Rik* | antisense | RIKEN cDNA 4930488L21 gene [Source:MGI Symbol;Acc:MGI:1923059] | 2.025266 | 0.00011055 |
| ENSMUSG00000105265 | 320478 | *Sox2ot* | processed_transcript | SOX2 overlapping transcript (non-protein coding) [Source:MGI Symbol;Acc:MGI:2444112] | 2.036342 | 0.00279392 |
| ENSMUSG00000097442 | NA | *Gm26632* | antisense | predicted gene, 26632 [Source:MGI Symbol;Acc:MGI:5477126] | 2.05817 | 0.00057593 |
| ENSMUSG00000086485 | NA | *Gm15458* | lincRNA | predicted gene 15458 [Source:MGI Symbol;Acc:MGI:3705291] | 2.068661 | 0.00702198 |
| ENSMUSG00000085785 | 66760 | *Sox5os3* | antisense | SRY (sex determining region Y)-box 5, opposite strand 3 [Source:MGI Symbol;Acc:MGI:1914010] | 2.072049 | 0.00314339 |
| ENSMUSG00000057715 | 320492 | *A830018L16Rik* | protein_coding | RIKEN cDNA A830018L16 gene [Source:MGI Symbol;Acc:MGI:2444149] | 2.072475 | 0.0463393 |
| ENSMUSG00000104801 | NA | *Gm43834* | TEC | predicted gene 43834 [Source:MGI Symbol;Acc:MGI:5663971] | 2.089419 | 0.0162712 |
| ENSMUSG00000097245 | NA | *Gm5421* | processed_pseudogene | predicted gene 5421 [Source:MGI Symbol;Acc:MGI:3645718] | 2.097842 | 0.0187688 |
| ENSMUSG00000107794 | NA | *Gm44095* | TEC | predicted gene, 44095 [Source:MGI Symbol;Acc:MGI:5690487] | 2.097998 | 0.0383494 |
| ENSMUSG00000022622 | 11434 | *Acr* | protein_coding | acrosin prepropeptide [Source:MGI Symbol;Acc:MGI:87884] | 2.109631 | 0.00573082 |
| ENSMUSG00000020963 | 22095 | *Tshr* | protein_coding | thyroid stimulating hormone receptor [Source:MGI Symbol;Acc:MGI:98849] | 2.118431 | 0.0142339 |
| ENSMUSG00000118330 | NA | *AC132148.1* | antisense | novel transcript, antisense to Dagla | 2.130168 | 0.00359827 |
| ENSMUSG00000109648 | NA | *Svet1* | TEC | subventricular expressed transcript 1 [Source:MGI Symbol;Acc:MGI:2385655] | 2.154123 | 0.0347457 |
| ENSMUSG00000045875 | 11549 | *Adra1a* | protein_coding | adrenergic receptor, alpha 1a [Source:MGI Symbol;Acc:MGI:104773] | 2.158667 | 0.0362557 |
| ENSMUSG00000086785 | NA | *Gm6081* | processed_pseudogene | predicted gene 6081 [Source:MGI Symbol;Acc:MGI:3645683] | 2.189778 | 0.00111700 |
| ENSMUSG00000102250 | NA | *Gm38260* | TEC | predicted gene, 38260 [Source:MGI Symbol;Acc:MGI:5611488] | 2.20764 | 0.0060334 |
| ENSMUSG00000112990 | NA | *Gm47372* | TEC | predicted gene, 47372 [Source:MGI Symbol;Acc:MGI:6096284] | 2.21299 | 0.00040232 |
| ENSMUSG00000064225 | 75552 | *Paqr9* | protein_coding | progestin and adipoQ receptor family member IX [Source:MGI Symbol;Acc:MGI:1922802] | 2.238633 | 0.037831 |
| ENSMUSG00000035226 | 241770 | *Rims4* | protein_coding | regulating synaptic membrane exocytosis 4 [Source:MGI Symbol;Acc:MGI:2674366] | 2.249374 | 0.0261590 |
| ENSMUSG00000117465 | NA | *AC102496.1* | lincRNA | novel transcript | 2.251607 | 0.00469068 |
| ENSMUSG00000081838 | NA | *Gm13038* | processed_pseudogene | predicted gene 13038 [Source:MGI Symbol;Acc:MGI:3650684] | 2.260867 | 0.00014872 |
| ENSMUSG00000089783 | NA | *Gm454* | lincRNA | predicted gene 454 [Source:MGI Symbol;Acc:MGI:2685300] | 2.280125 | 0.0273077 |
| ENSMUSG00000070390 | 637515 | *Nlrp1b* | protein_coding | NLR family, pyrin domain containing 1B [Source:MGI Symbol;Acc:MGI:3582959] | 2.30229 | 8.36E-0 |
| ENSMUSG00000108141 | NA | *Gm44079* | antisense | predicted gene, 44079 [Source:MGI Symbol;Acc:MGI:5690471] | 2.325855 | 0.00305294 |
| ENSMUSG00000105726 | NA | *Gm42443* | TEC | predicted gene 42443 [Source:MGI Symbol;Acc:MGI:5662580] | 2.34345 | 0.0008189 |
| ENSMUSG00000039200 | 75329 | *Atf7ip2* | protein_coding | activating transcription factor 7 interacting protein 2 [Source:MGI Symbol;Acc:MGI:1922579] | 2.358692 | 0.00025685 |
| ENSMUSG00000044349 | 319317 | *Snhg11* | protein_coding | small nucleolar RNA host gene 11 [Source:MGI Symbol;Acc:MGI:2441845] | 2.377717 | 0.00108775 |
| ENSMUSG00000117666 | 225852 | *AC131692.1* | protein_coding | Mus musculus predicted 550 (Gm550), mRNA. [Source:RefSeq mRNA;Acc:NM_001362427] | 2.37872 | 0.00242976 |
| ENSMUSG00000029167 | 19017 | *Ppargc1a* | protein_coding | peroxisome proliferative activated receptor, gamma, coactivator 1 alpha [Source:MGI Symbol;Acc:MGI:1342774] | 2.380016 | 0.0149969 |
| ENSMUSG00000111207 | NA | *Gm8125* | processed_pseudogene | predicted gene 8125 [Source:MGI Symbol;Acc:MGI:3648562] | 2.380968 | 0.0149164 |
| ENSMUSG00000053054 | 69117 | *Adh6a* | protein_coding | alcohol dehydrogenase 6A (class V) [Source:MGI Symbol;Acc:MGI:1916367] | 2.38639 | 0.0447617 |
| ENSMUSG00000094841 | NA | *Gm10610* | lincRNA | predicted gene 10610 [Source:MGI Symbol;Acc:MGI:3642045] | 2.387216 | 0.00527598 |
| ENSMUSG00000108442 | NA | *Rpl15-ps5* | processed_pseudogene | ribosomal protein L15, pseudogene 5 [Source:MGI Symbol;Acc:MGI:5010232] | 2.387714 | 0.00010218 |
| ENSMUSG00000050423 | 76487 | *Ppp1r3g* | protein_coding | protein phosphatase 1, regulatory subunit 3G [Source:MGI Symbol;Acc:MGI:1923737] | 2.398954 | 0.0233500 |
| ENSMUSG00000044667 | 229791 | *Plppr4* | protein_coding | phospholipid phosphatase related 4 [Source:MGI Symbol;Acc:MGI:106530] | 2.418315 | 0.0137020 |
| ENSMUSG00000098383 | NA | *Gm27197* | antisense | predicted gene 27197 [Source:MGI Symbol;Acc:MGI:5521040] | 2.441261 | 0.00182277 |
| ENSMUSG00000055373 | 14348 | *Fut9* | protein_coding | fucosyltransferase 9 [Source:MGI Symbol;Acc:MGI:1330859] | 2.455653 | 0.0161981 |
| ENSMUSG00000062093 | NA | *Gm10110* | transcribed_processed_pseudogene | predicted gene 10110 [Source:MGI Symbol;Acc:MGI:3641718] | 2.524284 | 0.00187615 |
| ENSMUSG00000105442 | NA | *Gm42614* | TEC | predicted gene 42614 [Source:MGI Symbol;Acc:MGI:5662751] | 2.582344 | 0.0384395 |
| ENSMUSG00000069830 | 195046 | *Nlrp1a* | protein_coding | NLR family, pyrin domain containing 1A [Source:MGI Symbol;Acc:MGI:2684861] | 2.651208 | 0.00151537 |
| ENSMUSG00000105622 | NA | *Gm42615* | TEC | predicted gene 42615 [Source:MGI Symbol;Acc:MGI:5662752] | 2.6683 | 0.0414285 |
| ENSMUSG00000090955 | NA | *Gm17097* | processed_pseudogene | predicted gene 17097 [Source:MGI Symbol;Acc:MGI:4937924] | 2.713069 | 0.00729055 |
| ENSMUSG00000037071 | 20249 | *Scd1* | protein_coding | stearoyl-Coenzyme A desaturase 1 [Source:MGI Symbol;Acc:MGI:98239] | 2.733032 | 0.00032817 |
| ENSMUSG00000028341 | 18124 | *Nr4a3* | protein_coding | nuclear receptor subfamily 4, group A, member 3 [Source:MGI Symbol;Acc:MGI:1352457] | 3.009918 | 0.010881 |

| | | | | | | |
|---|---|---|---|---|---|---|
| ENSMUSG00000033882 | 633285 | *Rbm46* | protein_coding | RNA binding motif protein 46 [Source:MGI Symbol;Acc:MGI:3645057] | 3.026779 | 0.00061341 |
| ENSMUSG00000069584 | NA | *Gm10272* | protein_coding | predicted gene 10272 [Source:MGI Symbol;Acc:MGI:3642183] | 3.091171 | 0.00311695 |
| ENSMUSG00000104696 | NA | *Gm42946* | TEC | predicted gene 42946 [Source:MGI Symbol;Acc:MGI:5663083] | 3.125405 | 0.00721070 |
| ENSMUSG00000027513 | 18534 | *Pck1* | protein_coding | phosphoenolpyruvate carboxykinase 1, cytosolic [Source:MGI Symbol;Acc:MGI:97501] | 3.15991 | 0.0236407 |
| ENSMUSG00000115702 | NA | *Gm5206* | transcribed_processed_pseudogene | predicted pseudogene 5206 [Source:MGI Symbol;Acc:MGI:3645516] | 3.396874 | 0.00013707 |
| ENSMUSG00000032401 | 235435 | *Lctl* | protein_coding | lactase-like [Source:MGI Symbol;Acc:MGI:2183549] | 3.472537 | 0.0301608 |
| ENSMUSG00000097532 | NA | *Gm4349* | lincRNA | predicted gene 4349 [Source:MGI Symbol;Acc:MGI:3782533] | 3.506262 | 0.00022496 |
| ENSMUSG00000041698 | 28248 | *Slco1a1* | protein_coding | solute carrier organic anion transporter family, member 1a1 [Source:MGI Symbol;Acc:MGI:1351891] | 3.508825 | 0.0465098 |
| ENSMUSG00000108405 | NA | *Ywhaq-ps1* | processed_pseudogene | tyrosine 3-monooxygenase/tryptophan 5-monooxygenase activation protein theta, pseudogene 1 [Source:MGI Symbol;Acc:MGI:5010024] | 3.531612 | 0.00109884 |
| ENSMUSG00000087382 | NA | *Ctcflos* | antisense | CCCTC-binding factor (zinc finger protein)-like, opposite strand [Source:MGI Symbol;Acc:MGI:1921411] | 3.574997 | 0.00102563 |
| ENSMUSG00000081303 | NA | *Gm16011* | processed_pseudogene | predicted gene 16011 [Source:MGI Symbol;Acc:MGI:3801796] | 3.725768 | 0.00046571 |
| ENSMUSG00000081824 | NA | *BC002163* | processed_pseudogene | cDNA sequence BC002163 [Source:MGI Symbol;Acc:MGI:3612445] | 3.800623 | 0.0121500 |
| ENSMUSG00000033196 | 17882 | *Myh2* | protein_coding | myosin, heavy polypeptide 2, skeletal muscle, adult [Source:MGI Symbol;Acc:MGI:1339710] | 3.957639 | 1.25E-0[illegible] |
| ENSMUSG00000104232 | NA | *Gm37590* | TEC | predicted gene, 37590 [Source:MGI Symbol;Acc:MGI:5610818] | 4.269763 | 0.0007079 |
| ENSMUSG00000055197 | 260298 | *Fev* | protein_coding | FEV (ETS oncogene family) [Source:MGI Symbol;Acc:MGI:2449712] | 4.359803 | 0.00173289 |
| ENSMUSG00000067017 | NA | *Gm3608* | pseudogene | predicted gene 3608 [Source:MGI Symbol;Acc:MGI:3804932] | 4.368809 | 1.37E-0[illegible] |
| ENSMUSG00000117695 | NA | *AC161579.1* | lincRNA | novel transcript | 4.740878 | 0.00549716 |
| ENSMUSG00000043618 | NA | *Eif5al3-ps* | processed_pseudogene | eukaryotic translation initiation factor 5A-like 3, pseudogene [Source:MGI Symbol;Acc:MGI:3643585] | 4.923526 | 7.71E-0[illegible] |
| ENSMUSG00000083240 | NA | *Gm13453* | processed_pseudogene | predicted gene 13453 [Source:MGI Symbol;Acc:MGI:3651618] | 6.665199 | 7.43E-0[illegible] |
| ENSMUSG00000091383 | NA | *Hist1h2al* | processed_pseudogene | histone cluster 1, H2al [Source:MGI Symbol;Acc:MGI:3646032] | 7.007794 | 6.61E-1[illegible] |
| ENSMUSG00000099216 | NA | *Obox8* | protein_coding | oocyte specific homeobox 8 [Source:MGI Symbol;Acc:MGI:3645855] | 7.144301 | 1.66E-1[illegible] |
| ENSMUSG00000040264 | 14468 | *Gbp2b* | protein_coding | guanylate binding protein 2b [Source:MGI Symbol;Acc:MGI:95666] | 7.365655 | 5.01E-0[illegible] |
| ENSMUSG00000059751 | NA | *Rps3a3* | processed_pseudogene | ribosomal protein S3A3 [Source:MGI Symbol;Acc:MGI:3643406] | 7.967852 | 5.09E-1[illegible] |
