## SupplementalTable11 for "An enhancer:involucrin regulatory module impacts human skin barrier adaptation out-of-Africa and modifies atopic dermatitis risk"

**Table S11. Ranked list of differentially accessible regions between 923del/del and WT mice epidermis from ATAC-seq.**

| *Chromosome* | *start* | *end* | *Fold* | *FDR* |
|---|---|---|---|---|
| chrX | 169993996 | 169994243 | -4.64 | 0.0221 |
| chrX | 169996914 | 169998486 | -3.7 | 0.0266 |
| chr3 | 93176562 | 93176984 | -2.76 | 0.0462 |
| chr3 | 93814593 | 93815096 | -2.56 | 0.00859 |
| chr3 | 79242295 | 79242558 | 2.89 | 0.0462 |
| chrX | 50591524 | 50591728 | 3.78 | 0.0378 |
| chrX | 50611755 | 50611998 | 4.19 | 0.0378 |
| chr3 | 78966708 | 78967396 | 4.34 | 3.81E-07 |
| chr3 | 92978570 | 92978806 | 4.39 | 0.00104 |
| chr3 | 93530552 | 93530842 | 4.82 | 3.81E-07 |
