## SupplementalTable12 for "An enhancer:involucrin regulatory module impacts human skin barrier adaptation out-of-Africa and modifies atopic dermatitis risk"

**Table S12. Ranked list of differentially accessible regions between 923large/large and WT mice epidermis from ATAC-seq.**

| *Chromosome* | *start* | *end* | *Fold* | *FDR* |
|---|---|---|---|---|
| chrX | 169993996 | 169994243 | -5.41 | 9.91E-05 |
| chrX | 169996914 | 169998486 | -3.83 | 0.000487 |
| chr8 | 119234019 | 119234204 | -3.66 | 0.0297 |
| chr7 | 118642175 | 118642353 | -3.63 | 0.0464 |
| chr12 | 118918105 | 118918297 | -3.59 | 0.0441 |
| chr8 | 86904158 | 86904360 | -3.47 | 0.0243 |
| chr15 | 88995738 | 88995926 | -3.45 | 0.0159 |
| chr14 | 37306986 | 37307136 | -3.43 | 0.0431 |
| chr14 | 7972236 | 7972440 | -3.41 | 0.00792 |
| chr5 | 38901570 | 38901772 | -3.23 | 0.0323 |
| chr3 | 93780248 | 93780977 | -2.73 | 0.00728 |
| chr3 | 93814593 | 93815096 | -2.53 | 0.00728 |
| chr13 | 21172295 | 21172487 | 2.99 | 0.0431 |
| chr3 | 92978570 | 92978806 | 3.64 | 0.0431 |
| chr3 | 92586198 | 92586460 | 3.64 | 0.0471 |
| chrX | 52243896 | 52244124 | 3.72 | 0.0104 |
| chr3 | 78966708 | 78967396 | 3.99 | 0.000119 |
| chr5 | 123127103 | 123127309 | 4 | 0.0186 |
| chr3 | 92579546 | 92579913 | 4.54 | 0.000487 |
| chr3 | 92583166 | 92583453 | 4.61 | 9.91E-05 |
| chr3 | 93530552 | 93530842 | 4.84 | 8.89E-06 |
| chr3 | 92609764 | 92610128 | 5.5 | 5.05E-07 |
| chr1 | 24613142 | 24615948 | 5.69 | 0.0104 |
