## SupplementalTable13 for "An enhancer:involucrin regulatory module impacts human skin barrier adaptation out-of-Africa and modifies atopic dermatitis risk"

**Table S13. Variants in the 923 enhancer region found in Atopic Dermatitis Patients.**

| *rs number* | *GRCh37 (Chr1)* | *Ref* | *Alt* | *Global Frequency (GnomAD)* | *Global Frequency (1000G)* | *Frequency in AD subjects* |
|---|---|---|---|---|---|---|
| rs142064645 | 152877402 | C | A | 0.002102 | 0.002 | 0.009 |
| rs116579812 | 152877524 | T | C | 0.01679 | 0.025 | 0.027 |
| rs116579812 | 152877524 | TG | CA | | | 0.009 |
| rs75653984 | 152877795 | G | A | 0.02241 | 0.026 | 0.045 |
| rs191034924 | 152877827 | A | C | 0.0000957 | 0 | 0.009 |
| rs149241485 | 152877941 | A | T | 0.0167 | 0.025 | 0.018 |
| rs143834829 | 152877966 | G | A | 0.0168 | 0.025 | 0.045 |
| rs138479364 | 152877993 | G | A | 0.01663 | 0.025 | 0.045 |
| rs35991466 | 152878006 | G | A | 0.05352 | 0.037 | 0.045 |
| rs111360880 | 152878021 | T | C | 0.03736 | 0.045 | 0.036 |
| rs186688062 | 152878126 | G | A | 0.006533 | 0.005 | 0.009 |
| rs76506275 | 152878179 | G | A | 0.022 | 0.026 | 0.045 |
| rs79007216 | 152878189 | C | T | 0.01674 | 0.025 | 0.045 |
| rs1974141 | 152878222 | A | G | 0.03217 | 0.177 | 0.1802 |
| rs77106372 | 152878254 | G | A | 0.001596 | 0.002 | 0.018 |
| rs114630587 | 152878467 | G | T | 0.005319 | 0.006 | 0.009 |
| rs115251823 | 152878568 | G | T | 0.006021 | 0.007 | 0.018 |
| rs78868757 | 152878599 | G | A | 0.02249 | 0.026 | 0.045 |
| rs150846807 | 152878695 | C | T | 0.004777 | 0.004 | 0.009 |
| rs1854780 | 152878900 | G | A | 0.005574 | 0.008 | 0.027 |
| rs12036697 | 152878909 | A | G | 0.03206 | 0.177 | 0.1802 |
| rs1344252044 | 152879069 | G | A | | | 0.009 |
| rs12239648 | 152879091 | A | T | 0.08932 | 0.078 | 0.0991 |
| rs16834746 | 152879096 | T | C | 0.03179 | 0.177 | 0.1712 |
| rs12240158 | 152879103 | C | T | 0.1285 | 0.129 | 0.1802 |
| rs139775257 | 152879143 | G | A | 0.005989 | 0.012 | 0.018 |
| rs76513010 | 152879153 | T | C | 0.003696 | 0.004 | 0.009 |
| rs114911757 | 152879362 | G | A | 0.01742 | 0.022 | 0.036 |
| rs4845327 | 152879512 | G | T | 0.2484 | 0.407 | 0.2432 |
| rs147621599 | 152879576 | T | A | 0.003216 | 0.002 | 0.036 |
