## SupplementalTable14 for "An enhancer:involucrin regulatory module impacts human skin barrier adaptation out-of-Africa and modifies atopic dermatitis risk"

**Table S14. GTEX eQTLs for IVL in sun-exposed and no sun-exposed skins (GRCh37/hg37).**

| *Gencode Id* | *Gene Symbol* | *Variant Id* | *SNP Id* | *P-Value* | *Normalized Effect Size (NES)* | *Tissue* |
|---|---|---|---|---|---|---|
| ENSG00000163207.5 | *IVL* | .152688262_T_TGGG_b37 | rs397686224 | 0.000086 | -0.18 | Skin - Sun Exposed (Lower leg) |
| ENSG00000163207.5 | *IVL* | .152789507_A_G_b37 | rs12027736 | 0.000012 | -0.5 | Skin - Sun Exposed (Lower leg) |
| ENSG00000163207.5 | *IVL* | .152798560_G_A_b37 | rs4845325 | 4.00E-07 | -0.55 | Skin - Sun Exposed (Lower leg) |
| ENSG00000163207.5 | *IVL* | .152808977_G_A_b37 | rs61482767 | 5.70E-07 | -0.53 | Skin - Sun Exposed (Lower leg) |
| ENSG00000163207.5 | *IVL* | .152815406_G_A_b37 | rs7552416 | 5.70E-07 | -0.53 | Skin - Sun Exposed (Lower leg) |
| ENSG00000163207.5 | *IVL* | .152827652_G_A_b37 | rs10494280 | 3.30E-08 | -0.63 | Skin - Sun Exposed (Lower leg) |
| ENSG00000163207.5 | *IVL* | .152830714_A_G_b37 | rs58021079 | 4.80E-09 | -0.87 | Skin - Sun Exposed (Lower leg) |
| ENSG00000163207.5 | *IVL* | .152837434_C_T_b37 | rs11205126 | 4.00E-07 | -0.55 | Skin - Sun Exposed (Lower leg) |
| ENSG00000163207.5 | *IVL* | .152843377_T_C_b37 | rs4240863 | 4.00E-07 | -0.55 | Skin - Sun Exposed (Lower leg) |
| ENSG00000163207.5 | *IVL* | .152844142_G_A_b37 | rs11205128 | 0.000056 | -0.32 | Skin - Sun Exposed (Lower leg) |
| ENSG00000163207.5 | *IVL* | .152844585_C_T_b37 | rs16834715 | 0.000022 | -0.35 | Skin - Sun Exposed (Lower leg) |
| ENSG00000163207.5 | *IVL* | .152846219_TTAAA_T_b37 | rs59995047 | 0.0000097 | -0.35 | Skin - Sun Exposed (Lower leg) |
| ENSG00000163207.5 | *IVL* | .152846400_G_A_b37 | rs12022870 | 1.20E-08 | -0.62 | Skin - Sun Exposed (Lower leg) |
| ENSG00000163207.5 | *IVL* | .15284655.T_G_b37 | rs12048655 | 0.0000089 | -0.36 | Skin - Sun Exposed (Lower leg) |
| ENSG00000163207.5 | *IVL* | .15284662.A_G_b37 | rs12047888 | 5.40E-11 | -0.96 | Skin - Sun Exposed (Lower leg) |
| ENSG00000163207.5 | *IVL* | .152847483_TA_T_b37 | rs796268334 | 0.000014 | -0.33 | Skin - Sun Exposed (Lower leg) |
| ENSG00000163207.5 | *IVL* | .152848623_C_A_b37 | rs2339383 | 1.30E-08 | -0.62 | Skin - Sun Exposed (Lower leg) |
| ENSG00000163207.5 | *IVL* | .152848839_G_A_b37 | rs4845489 | 0.000013 | -0.33 | Skin - Sun Exposed (Lower leg) |
| ENSG00000163207.5 | *IVL* | .152850322_G_T_b37 | rs12021601 | 1.30E-08 | -0.62 | Skin - Sun Exposed (Lower leg) |
| ENSG00000163207.5 | *IVL* | .152850906_G_A_b37 | rs3737862 | 1.30E-08 | -0.62 | Skin - Sun Exposed (Lower leg) |
| ENSG00000163207.5 | *IVL* | .152850938_A_C_b37 | rs3737861 | 0.000016 | -0.16 | Skin - Sun Exposed (Lower leg) |
| ENSG00000163207.5 | *IVL* | .152851315_C_T_b37 | rs60566199 | 1.30E-08 | -0.62 | Skin - Sun Exposed (Lower leg) |
| ENSG00000163207.5 | *IVL* | .152851698_C_A_b37 | rs73006590 | 0.000024 | -0.36 | Skin - Sun Exposed (Lower leg) |
| ENSG00000163207.5 | *IVL* | .152851807_C_T_b37 | rs113126048 | 0.000026 | -0.36 | Skin - Sun Exposed (Lower leg) |
| ENSG00000163207.5 | *IVL* | .152853278_T_C_b37 | rs16834734 | 5.40E-11 | -0.96 | Skin - Sun Exposed (Lower leg) |
| ENSG00000163207.5 | *IVL* | .152853586_G_A_b37 | rs77107784 | 1.30E-08 | -0.62 | Skin - Sun Exposed (Lower leg) |
| ENSG00000163207.5 | *IVL* | .152854600_T_C_b37 | rs7556490 | 0.000026 | -0.36 | Skin - Sun Exposed (Lower leg) |
| ENSG00000163207.5 | *IVL* | .15285467.C_T_b37 | rs7542656 | 0.000026 | -0.36 | Skin - Sun Exposed (Lower leg) |
| ENSG00000163207.5 | *IVL* | .152854924_A_G_b37 | rs4845494 | 0.000026 | -0.36 | Skin - Sun Exposed (Lower leg) |
| ENSG00000163207.5 | *IVL* | .152855132_G_C_b37 | rs4363387 | 0.000025 | -0.36 | Skin - Sun Exposed (Lower leg) |
| ENSG00000163207.5 | *IVL* | .152856084_G_C_b37 | rs11801996 | 0.000026 | -0.36 | Skin - Sun Exposed (Lower leg) |
| ENSG00000163207.5 | *IVL* | .152856302_A_G_b37 | rs11205129 | 0.000026 | -0.36 | Skin - Sun Exposed (Lower leg) |
| ENSG00000163207.5 | *IVL* | .152860594_T_C_b37 | rs36033488 | 0.000001 | -0.29 | Skin - Sun Exposed (Lower leg) |
| ENSG00000163207.5 | *IVL* | .152861996_G_A_b37 | rs12028228 | 0.000013 | -0.36 | Skin - Sun Exposed (Lower leg) |
| ENSG00000163207.5 | *IVL* | .152869494_A_G_b37 | rs6668295 | 5.40E-11 | -0.96 | Skin - Sun Exposed (Lower leg) |
| ENSG00000163207.5 | *IVL* | .15286954.T_C_b37 | rs34198967 | 2.90E-07 | -0.29 | Skin - Sun Exposed (Lower leg) |
| ENSG00000163207.5 | *IVL* | .152871900_C_T_b37 | rs79834096 | 5.40E-11 | -0.96 | Skin - Sun Exposed (Lower leg) |
| ENSG00000163207.5 | *IVL* | .152872885_G_A_b37 | rs141317883 | 5.40E-11 | -0.96 | Skin - Sun Exposed (Lower leg) |
| ENSG00000163207.5 | *IVL* | .152875358_C_G_b37 | rs58290373 | 3.50E-09 | -0.81 | Skin - Sun Exposed (Lower leg) |
| ENSG00000163207.5 | *IVL* | .152878006_G_A_b37 | rs35991466 | 1.90E-07 | -0.31 | Skin - Sun Exposed (Lower leg) |
| ENSG00000163207.5 | *IVL* | .152878222_A_G_b37 | rs1974141 | 5.40E-11 | -0.96 | Skin - Sun Exposed (Lower leg) |
| ENSG00000163207.5 | *IVL* | .152878909_A_G_b37 | rs12036697 | 5.40E-11 | -0.96 | Skin - Sun Exposed (Lower leg) |
| ENSG00000163207.5 | *IVL* | .15287909.A_T_b37 | rs12239648 | 6.40E-08 | -0.3 | Skin - Sun Exposed (Lower leg) |
| ENSG00000163207.5 | *IVL* | .152879096_T_C_b37 | rs16834746 | 5.40E-11 | -0.96 | Skin - Sun Exposed (Lower leg) |
| ENSG00000163207.5 | *IVL* | .152879103_C_T_b37 | rs12240158 | 2.00E-08 | -0.3 | Skin - Sun Exposed (Lower leg) |
| ENSG00000163207.5 | *IVL* | .152879512_G_T_b37 | rs4845327 | 2.20E-14 | -0.36 | Skin - Sun Exposed (Lower leg) |
| ENSG00000163207.5 | *IVL* | .152880672_T_C_b37 | rs1854779 | 0.000012 | -0.25 | Skin - Not Sun Exposed (Suprapubic) |
| ENSG00000163207.5 | *IVL* | .152880672_T_C_b37 | rs1854779 | 7.00E-15 | -0.36 | Skin - Sun Exposed (Lower leg) |
| ENSG00000163207.5 | *IVL* | .15288121.A_C_b37 | rs16834751 | 4.70E-09 | -0.67 | Skin - Sun Exposed (Lower leg) |
| ENSG00000163207.5 | *IVL* | .152881350_C_T_b37 | rs34593101 | 2.90E-07 | -0.29 | Skin - Sun Exposed (Lower leg) |
| ENSG00000163207.5 | *IVL* | .152881430_T_C_b37 | rs4523473 | 0.000051 | -0.23 | Skin - Not Sun Exposed (Suprapubic) |
| ENSG00000163207.5 | *IVL* | .152881430_T_C_b37 | rs4523473 | 1.30E-14 | -0.36 | Skin - Sun Exposed (Lower leg) |
| ENSG00000163207.5 | *IVL* | .152881649_T_C_b37 | rs11205130 | 5.40E-11 | -0.96 | Skin - Sun Exposed (Lower leg) |
| ENSG00000163207.5 | *IVL* | .152881678_G_A_b37 | rs11205131 | 2.50E-11 | -0.85 | Skin - Sun Exposed (Lower leg) |
| ENSG00000163207.5 | *IVL* | .152881689_A_G_b37 | rs7528862 | 0.000012 | -0.25 | Skin - Not Sun Exposed (Suprapubic) |
| ENSG00000163207.5 | *IVL* | .152881689_A_G_b37 | rs7528862 | 7.00E-15 | -0.36 | Skin - Sun Exposed (Lower leg) |
| ENSG00000163207.5 | *IVL* | .152881746_C_G_b37 | rs7517189 | 0.000012 | -0.25 | Skin - Not Sun Exposed (Suprapubic) |
| ENSG00000163207.5 | *IVL* | .152881746_C_G_b37 | rs7517189 | 7.00E-15 | -0.36 | Skin - Sun Exposed (Lower leg) |
| ENSG00000163207.5 | *IVL* | .152881802_G_A_b37 | rs7539232 | 0.000012 | -0.25 | Skin - Not Sun Exposed (Suprapubic) |
| ENSG00000163207.5 | *IVL* | .152881802_G_A_b37 | rs7539232 | 7.00E-15 | -0.36 | Skin - Sun Exposed (Lower leg) |
| ENSG00000163207.5 | *IVL* | .152882135_A_G_b37 | rs11205132 | 0.000012 | -0.25 | Skin - Not Sun Exposed (Suprapubic) |
| ENSG00000163207.5 | *IVL* | .152882135_A_G_b37 | rs11205132 | 7.00E-15 | -0.36 | Skin - Sun Exposed (Lower leg) |
| ENSG00000163207.5 | *IVL* | .152882610_A_G_b37 | rs2229496 | 0.000012 | -0.25 | Skin - Not Sun Exposed (Suprapubic) |
| ENSG00000163207.5 | *IVL* | .152882610_A_G_b37 | rs2229496 | 7.00E-15 | -0.36 | Skin - Sun Exposed (Lower leg) |
| ENSG00000163207.5 | *IVL* | .152883608_G_A_b37 | rs7545413 | 0.000012 | -0.25 | Skin - Not Sun Exposed (Suprapubic) |
| ENSG00000163207.5 | *IVL* | .152883608_G_A_b37 | rs7545413 | 7.00E-15 | -0.36 | Skin - Sun Exposed (Lower leg) |
| ENSG00000163207.5 | *IVL* | .152883680_A_G_b37 | rs7535306 | 0.000014 | -0.25 | Skin - Not Sun Exposed (Suprapubic) |
| ENSG00000163207.5 | *IVL* | .152883680_A_G_b37 | rs7535306 | 4.20E-15 | -0.37 | Skin - Sun Exposed (Lower leg) |
| ENSG00000163207.5 | *IVL* | .15288371.G_C_b37 | rs7545520 | 0.000012 | -0.25 | Skin - Not Sun Exposed (Suprapubic) |
| ENSG00000163207.5 | *IVL* | .15288371.G_C_b37 | rs7545520 | 7.00E-15 | -0.36 | Skin - Sun Exposed (Lower leg) |
| ENSG00000163207.5 | *IVL* | .15288411.G_A_b37 | rs913996 | 0.000012 | -0.25 | Skin - Not Sun Exposed (Suprapubic) |
| ENSG00000163207.5 | *IVL* | .15288411.G_A_b37 | rs913996 | 7.00E-15 | -0.36 | Skin - Sun Exposed (Lower leg) |
| ENSG00000163207.5 | *IVL* | .152884246_G_T_b37 | rs12407682 | 0.000012 | -0.25 | Skin - Not Sun Exposed (Suprapubic) |
| ENSG00000163207.5 | *IVL* | .152884246_G_T_b37 | rs12407682 | 7.00E-15 | -0.36 | Skin - Sun Exposed (Lower leg) |
| ENSG00000163207.5 | *IVL* | .152884247_A_G_b37 | rs12406539 | 0.000012 | -0.25 | Skin - Not Sun Exposed (Suprapubic) |
| ENSG00000163207.5 | *IVL* | .152884247_A_G_b37 | rs12406539 | 7.00E-15 | -0.36 | Skin - Sun Exposed (Lower leg) |
| ENSG00000163207.5 | *IVL* | .152884438_C_T_b37 | rs3820136 | 0.000037 | -0.24 | Skin - Not Sun Exposed (Suprapubic) |
| ENSG00000163207.5 | *IVL* | .152884438_C_T_b37 | rs3820136 | 2.20E-13 | -0.35 | Skin - Sun Exposed (Lower leg) |
| ENSG00000163207.5 | *IVL* | .152884526_A_AC_b37 | rs71582272 | 0.000004 | -0.25 | Skin - Sun Exposed (Lower leg) |
| ENSG00000163207.5 | *IVL* | .152885500_C_T_b37 | rs2879484 | 2.50E-11 | -0.85 | Skin - Sun Exposed (Lower leg) |
| ENSG00000163207.5 | *IVL* | .152885913_G_A_b37 | rs2879485 | 2.50E-11 | -0.85 | Skin - Sun Exposed (Lower leg) |
| ENSG00000163207.5 | *IVL* | .152886874_G_C_b37 | rs34599045 | 4.40E-07 | -0.27 | Skin - Sun Exposed (Lower leg) |
| ENSG00000163207.5 | *IVL* | .152887237_T_C_b37 | rs6587711 | 0.0000011 | -0.28 | Skin - Not Sun Exposed (Suprapubic) |
| ENSG00000163207.5 | *IVL* | .152887237_T_C_b37 | rs6587711 | 3.50E-12 | -0.33 | Skin - Sun Exposed (Lower leg) |
| ENSG00000163207.5 | *IVL* | .152888178_A_G_b37 | rs4845496 | 0.000038 | -0.14 | Skin - Not Sun Exposed (Suprapubic) |
| ENSG00000163207.5 | *IVL* | .152889327_T_C_b37 | rs7539610 | 9.40E-09 | -0.62 | Skin - Sun Exposed (Lower leg) |

| | | | | | | |
|---|---|---|---|---|---|---|
| ENSG00000163207.5 | *IVL* | .15288966.C_T_b37 | rs3845340 | 0.0000016 | -0.28 | Skin - Not Sun Exposed (Suprapubic) |
| ENSG00000163207.5 | *IVL* | .15288966.C_T_b37 | rs3845340 | 1.50E-12 | -0.33 | Skin - Sun Exposed (Lower leg) |
| ENSG00000163207.5 | *IVL* | .152889944_G_A_b37 | rs12239808 | 1.00E-09 | -0.69 | Skin - Sun Exposed (Lower leg) |
| ENSG00000163207.5 | *IVL* | .152890078_G_A_b37 | rs12239812 | 5.50E-11 | -0.96 | Skin - Sun Exposed (Lower leg) |
| ENSG00000163207.5 | *IVL* | .152890138_G_A_b37 | rs12239814 | 1.10E-09 | -0.69 | Skin - Sun Exposed (Lower leg) |
| ENSG00000163207.5 | *IVL* | .152890470_G_A_b37 | rs11586313 | 0.000011 | -0.15 | Skin - Not Sun Exposed (Suprapubic) |
| ENSG00000163207.5 | *IVL* | .152890568_G_A_b37 | rs11205138 | 0.0000036 | -0.16 | Skin - Not Sun Exposed (Suprapubic) |
| ENSG00000163207.5 | *IVL* | .152890604_A_G_b37 | rs11806470 | 9.10E-09 | -0.71 | Skin - Sun Exposed (Lower leg) |
| ENSG00000163207.5 | *IVL* | .152894355_C_T_b37 | rs4845328 | 2.10E-08 | -0.79 | Skin - Sun Exposed (Lower leg) |
| ENSG00000163207.5 | *IVL* | .152895586_G_A_b37 | rs79520860 | 5.20E-07 | -0.31 | Skin - Sun Exposed (Lower leg) |
| ENSG00000163207.5 | *IVL* | .152897923_C_T_b37 | rs12027303 | 0.000011 | -0.26 | Skin - Not Sun Exposed (Suprapubic) |
| ENSG00000163207.5 | *IVL* | .152897923_C_T_b37 | rs12027303 | 1.20E-11 | -0.33 | Skin - Sun Exposed (Lower leg) |
| ENSG00000163207.5 | *IVL* | .152898182_T_A_b37 | rs35686780 | 2.30E-07 | -0.32 | Skin - Sun Exposed (Lower leg) |
| ENSG00000163207.5 | *IVL* | .152898726_C_T_b37 | rs61491150 | 1.90E-09 | -0.85 | Skin - Sun Exposed (Lower leg) |
| ENSG00000163207.5 | *IVL* | .152900457_C_T_b37 | rs76915750 | 1.60E-07 | -0.32 | Skin - Sun Exposed (Lower leg) |
| ENSG00000163207.5 | *IVL* | .152900538_G_T_b37 | rs80207490 | 1.60E-07 | -0.32 | Skin - Sun Exposed (Lower leg) |
| ENSG00000163207.5 | *IVL* | .152901128_T_A_b37 | rs71517761 | 5.20E-07 | -0.31 | Skin - Sun Exposed (Lower leg) |
| ENSG00000163207.5 | *IVL* | .152901477_G_A_b37 | rs4240864 | 1.90E-09 | -0.85 | Skin - Sun Exposed (Lower leg) |
| ENSG00000163207.5 | *IVL* | .152901858_A_T_b37 | rs10888519 | 0.000011 | -0.27 | Skin - Not Sun Exposed (Suprapubic) |
| ENSG00000163207.5 | *IVL* | .152901858_A_T_b37 | rs10888519 | 4.70E-12 | -0.34 | Skin - Sun Exposed (Lower leg) |
| ENSG00000163207.5 | *IVL* | .152901930_A_G_b37 | rs71626737 | 5.20E-07 | -0.31 | Skin - Sun Exposed (Lower leg) |
| ENSG00000163207.5 | *IVL* | .152901982_G_C_b37 | rs78073526 | 5.20E-07 | -0.31 | Skin - Sun Exposed (Lower leg) |
| ENSG00000163207.5 | *IVL* | .152906086_G_A_b37 | rs71626738 | 5.20E-07 | -0.31 | Skin - Sun Exposed (Lower leg) |
| ENSG00000163207.5 | *IVL* | .152906133_T_C_b37 | rs71626739 | 5.20E-07 | -0.31 | Skin - Sun Exposed (Lower leg) |
| ENSG00000163207.5 | *IVL* | .152906716_A_C_b37 | rs12755214 | 1.60E-07 | -0.32 | Skin - Sun Exposed (Lower leg) |
| ENSG00000163207.5 | *IVL* | .15290972.T_G_b37 | rs35738590 | 4.90E-07 | -0.31 | Skin - Sun Exposed (Lower leg) |
| ENSG00000163207.5 | *IVL* | .15291050.G_A_b37 | rs112224666 | 4.00E-07 | -0.32 | Skin - Sun Exposed (Lower leg) |
| ENSG00000163207.5 | *IVL* | .152911018_A_G_b37 | rs150688308 | 5.20E-07 | -0.31 | Skin - Sun Exposed (Lower leg) |
| ENSG00000163207.5 | *IVL* | .152913068_G_A_b37 | rs12750261 | 5.80E-07 | -0.31 | Skin - Sun Exposed (Lower leg) |
| ENSG00000163207.5 | *IVL* | .152914338_T_A_b37 | rs36090881 | 0.000047 | -0.5 | Skin - Sun Exposed (Lower leg) |
| ENSG00000163207.5 | *IVL* | .152914389_G_A_b37 | rs12756264 | 0.0000017 | -0.3 | Skin - Sun Exposed (Lower leg) |
| ENSG00000163207.5 | *IVL* | .15291563.C_T_b37 | rs12724858 | 5.20E-07 | -0.31 | Skin - Sun Exposed (Lower leg) |
| ENSG00000163207.5 | *IVL* | .152915878_A_G_b37 | rs4845498 | 0.000072 | -0.46 | Skin - Sun Exposed (Lower leg) |
| ENSG00000163207.5 | *IVL* | .152917419_C_A_b37 | rs60500053 | 0.000047 | -0.5 | Skin - Sun Exposed (Lower leg) |
| ENSG00000163207.5 | *IVL* | .15291749.G_T_b37 | rs79934428 | 5.20E-07 | -0.31 | Skin - Sun Exposed (Lower leg) |
| ENSG00000163207.5 | *IVL* | .152920629_A_T_b37 | rs79402685 | 5.20E-07 | -0.31 | Skin - Sun Exposed (Lower leg) |
| ENSG00000163207.5 | *IVL* | .152920663_A_G_b37 | rs71626740 | 5.20E-07 | -0.31 | Skin - Sun Exposed (Lower leg) |
| ENSG00000163207.5 | *IVL* | .152920864_A_G_b37 | rs2879486 | 7.60E-10 | -0.92 | Skin - Sun Exposed (Lower leg) |
| ENSG00000163207.5 | *IVL* | .152920953_T_C_b37 | rs189320510 | 5.20E-07 | -0.31 | Skin - Sun Exposed (Lower leg) |
| ENSG00000163207.5 | *IVL* | .152920954_G_T_b37 | rs548020237 | 5.20E-07 | -0.31 | Skin - Sun Exposed (Lower leg) |
| ENSG00000163207.5 | *IVL* | .152923046_G_A_b37 | rs12737395 | 5.20E-07 | -0.31 | Skin - Sun Exposed (Lower leg) |
| ENSG00000163207.5 | *IVL* | .152923124_A_G_b37 | rs34889136 | 5.20E-07 | -0.31 | Skin - Sun Exposed (Lower leg) |
| ENSG00000163207.5 | *IVL* | .152937926_T_G_b37 | rs3885994 | 0.0000094 | -0.27 | Skin - Sun Exposed (Lower leg) |
| ENSG00000163207.5 | *IVL* | .152941289_A_G_b37 | rs71626743 | 0.0000077 | -0.27 | Skin - Sun Exposed (Lower leg) |
| ENSG00000163207.5 | *IVL* | .152945228_A_G_b37 | rs12032522 | 6.70E-07 | -0.71 | Skin - Sun Exposed (Lower leg) |
| ENSG00000163207.5 | *IVL* | .152946240_C_T_b37 | rs10494288 | 6.70E-07 | -0.71 | Skin - Sun Exposed (Lower leg) |
| ENSG00000163207.5 | *IVL* | .15295119.CT_C_b37 | rs34374141 | 0.0000077 | -0.27 | Skin - Sun Exposed (Lower leg) |
| ENSG00000163207.5 | *IVL* | .152952920_G_A_b37 | rs16834838 | 6.90E-07 | -0.71 | Skin - Sun Exposed (Lower leg) |
| ENSG00000163207.5 | *IVL* | .152955598_C_T_b37 | rs12047630 | 6.70E-07 | -0.71 | Skin - Sun Exposed (Lower leg) |
| ENSG00000163207.5 | *IVL* | .152960916_GC_G_b37 | rs201133598 | 0.00002 | -0.26 | Skin - Sun Exposed (Lower leg) |
| ENSG00000163207.5 | *IVL* | .152961342_C_T_b37 | rs10888521 | 0.0000048 | -0.71 | Skin - Sun Exposed (Lower leg) |
| ENSG00000163207.5 | *IVL* | .152966077_A_T_b37 | rs1415973 | 6.70E-07 | -0.71 | Skin - Sun Exposed (Lower leg) |
| ENSG00000163207.5 | *IVL* | .152968025_C_G_b37 | rs77702899 | 0.000021 | -0.26 | Skin - Sun Exposed (Lower leg) |
| ENSG00000163207.5 | *IVL* | .152971103_GGGCAGCCATGCGCT_G_b37 | rs71093258 | 0.000023 | -0.26 | Skin - Sun Exposed (Lower leg) |
| ENSG00000163207.5 | *IVL* | .152972016_G_A_b37 | rs71626744 | 0.000023 | -0.26 | Skin - Sun Exposed (Lower leg) |
| ENSG00000163207.5 | *IVL* | .152972747_G_A_b37 | rs17680493 | 0.000023 | -0.26 | Skin - Sun Exposed (Lower leg) |
| ENSG00000163207.5 | *IVL* | .152974920_T_A_b37 | rs28924725 | 0.000023 | -0.26 | Skin - Sun Exposed (Lower leg) |
| ENSG00000163207.5 | *IVL* | .152978307_G_T_b37 | rs78798271 | 0.00002 | -0.26 | Skin - Sun Exposed (Lower leg) |
| ENSG00000163207.5 | *IVL* | .152981048_A_G_b37 | rs12047099 | 3.10E-09 | -1 | Skin - Sun Exposed (Lower leg) |
| ENSG00000163207.5 | *IVL* | .152983644_A_G_b37 | rs35036485 | 0.00002 | -0.27 | Skin - Sun Exposed (Lower leg) |
| ENSG00000163207.5 | *IVL* | .152986260_T_G_b37 | rs12024694 | 3.30E-09 | -1 | Skin - Sun Exposed (Lower leg) |
| ENSG00000163207.5 | *IVL* | .15299616.G_C_b37 | rs16834871 | 5.50E-08 | -0.88 | Skin - Sun Exposed (Lower leg) |
| ENSG00000163207.5 | *IVL* | .152999973_T_G_b37 | rs12758478 | 0.000023 | -0.26 | Skin - Sun Exposed (Lower leg) |
| ENSG00000163207.5 | *IVL* | .153002303_G_A_b37 | rs12744709 | 0.000023 | -0.26 | Skin - Sun Exposed (Lower leg) |
| ENSG00000163207.5 | *IVL* | .153008252_A_T_b37 | rs34686286 | 0.000023 | -0.26 | Skin - Sun Exposed (Lower leg) |
| ENSG00000163207.5 | *IVL* | .153012765_T_C_b37 | rs1846857 | 0.000011 | -0.49 | Skin - Sun Exposed (Lower leg) |
| ENSG00000163207.5 | *IVL* | .153013948_T_C_b37 | rs2339501 | 3.50E-07 | -0.77 | Skin - Sun Exposed (Lower leg) |
| ENSG00000163207.5 | *IVL* | .15301431.G_A_b37 | rs1995307 | 3.10E-09 | -1 | Skin - Sun Exposed (Lower leg) |
| ENSG00000163207.5 | *IVL* | .153014845_G_A_b37 | rs28647391 | 3.10E-09 | -1 | Skin - Sun Exposed (Lower leg) |
| ENSG00000163207.5 | *IVL* | .153017026_A_T_b37 | rs34974823 | 0.000023 | -0.26 | Skin - Sun Exposed (Lower leg) |
| ENSG00000163207.5 | *IVL* | .153018913_A_T_b37 | rs4041337 | 3.60E-07 | -0.73 | Skin - Sun Exposed (Lower leg) |
| ENSG00000163207.5 | *IVL* | .153019087_C_T_b37 | rs3856020 | 3.60E-07 | -0.73 | Skin - Sun Exposed (Lower leg) |
| ENSG00000163207.5 | *IVL* | .153019217_A_T_b37 | rs71626747 | 0.000023 | -0.26 | Skin - Sun Exposed (Lower leg) |
| ENSG00000163207.5 | *IVL* | .153025233_C_G_b37 | rs12047225 | 3.20E-09 | -1 | Skin - Sun Exposed (Lower leg) |
| ENSG00000163207.5 | *IVL* | .153025499_A_G_b37 | rs34417380 | 0.000024 | -0.26 | Skin - Sun Exposed (Lower leg) |
| ENSG00000163207.5 | *IVL* | .15302572.A_T_b37 | rs12037754 | 3.10E-09 | -1 | Skin - Sun Exposed (Lower leg) |
| ENSG00000163207.5 | *IVL* | .153034230_T_C_b37 | rs12753609 | 0.000023 | -0.26 | Skin - Sun Exposed (Lower leg) |
| ENSG00000163207.5 | *IVL* | .153042009_A_ACAATACAAATTTCAAAATACAAT_b37 | rs559423296 | 0.0000014 | -0.84 | Skin - Sun Exposed (Lower leg) |
| ENSG00000163207.5 | *IVL* | .153042010_G_T_b37 | rs200671030 | 0.0000014 | -0.84 | Skin - Sun Exposed (Lower leg) |
| ENSG00000163207.5 | *IVL* | .153048524_T_C_b37 | rs11205183 | 0.0000015 | -0.83 | Skin - Sun Exposed (Lower leg) |
| ENSG00000163207.5 | *IVL* | .153050149_G_A_b37 | rs35000490 | 0.000081 | -0.25 | Skin - Sun Exposed (Lower leg) |
| ENSG00000163207.5 | *IVL* | .153058866_A_G_b37 | rs12750312 | 0.000023 | -0.26 | Skin - Sun Exposed (Lower leg) |
| ENSG00000163207.5 | *IVL* | .153085029_G_T_b37 | rs74133286 | 0.000032 | -0.53 | Skin - Sun Exposed (Lower leg) |
| ENSG00000163207.5 | *IVL* | .153101566_T_C_b37 | rs12408875 | 0.000032 | -0.53 | Skin - Sun Exposed (Lower leg) |
| ENSG00000163207.5 | *IVL* | .153103772_G_A_b37 | rs904949 | 0.000032 | -0.53 | Skin - Sun Exposed (Lower leg) |
