## SupplementalTable15 for "An enhancer:involucrin regulatory module impacts human skin barrier adaptation out-of-Africa and modifies atopic dermatitis risk"

**Table S15. Genes in the Epidermal Differentiation Complex (EDC) with annotated GTEX eQTLs in sun-exposed and non sunexposed skins (GTEX V7; GRCh37/hg37).**

| *Gene stable ID* | *Gene description/Gene type* | *Gene start (bp)* | *Gene end (bp)* | *Gene name* | *Sun-exposed Skin* | *Non sun-exposed skin* | *Lowest Effect size for alt allele* | *Highest effect size for alt allele* |
|---|---|---|---|---|---|---|---|---|
| ENSG00000237975 | FLG antisense RNA 1 [Source:HGNC Symbol;Acc:HGNC:27913] | 152168125 | 152445456 | *FLG-AS1* | Yes | Yes | -0.83 | 0.86 |
| ENSG00000197915 | hornerin [Source:HGNC Symbol;Acc:HGNC:20846] | 152212082 | 152224193 | *HRNR* | Yes | Yes | -0.6 | 0.56 |
| ENSG00000143536 | cornulin [Source:HGNC Symbol;Acc:HGNC:1230] | 152409243 | 152414263 | *CRNN* | Yes | Yes | -0.55 | 0.56 |
| ENSG00000185966 | late cornified envelope 3E [Source:HGNC Symbol;Acc:HGNC:29463] | 152565654 | 152566772 | *LCE3E* | Yes | No | -0.2 | -0.48 |
| ENSG00000163202 | late cornified envelope 3D [Source:HGNC Symbol;Acc:HGNC:16615] | 152579381 | 152580504 | *LCE3D* | Yes | No | -0.39 | 0.25 |
| ENSG00000244057 | late cornified envelope 3C [Source:HGNC Symbol;Acc:HGNC:16612] | 152600662 | 152601086 | *LCE3C* | Yes | Yes | -0.86 | 1.1 |
| ENSG00000185962 | late cornified envelope 3A [Source:HGNC Symbol;Acc:HGNC:29461] | 152622834 | 152623103 | *LCE3A* | Yes | Yes | 0.21 | 0.56 |
| ENSG00000187170 | late cornified envelope 4A [Source:HGNC Symbol;Acc:HGNC:16613] | 152708160 | 152709491 | *LCE4A* | Yes | Yes | -0.16 | -0.2 |
| ENSG00000198854 | chromosome 1 open reading frame 68 [Source:HGNC Symbol;Acc:HGNC:29468] | 152719522 | 152720470 | *C1orf68* | Yes | Yes | -0.55 | -0.098 |
| ENSG00000186226 | late cornified envelope 1E [Source:HGNC Symbol;Acc:HGNC:29466] | 152786214 | 152788426 | *LCE1E* | Yes | Yes | -0.68 | 0.63 |
| ENSG00000172155 | late cornified envelope 1D [Source:HGNC Symbol;Acc:HGNC:29465] | 152796751 | 152798181 | *LCE1D* | Yes | Yes | -0.91 | 0.44 |
| ENSG00000163207 | involucrin [Source:HGNC Symbol;Acc:HGNC:6187] | 152908545 | 152911886 | *IVL* | Yes | Yes | -1 | -0.16 |
| ENSG00000169469 | small proline rich protein 1B [Source:HGNC Symbol;Acc:HGNC:11260] | 153031202 | 153032900 | *SPRR1B* | Yes | Yes | -0.37 | 0.25 |
| ENSG00000163216 | small proline rich protein 2D [Source:HGNC Symbol;Acc:HGNC:11264] | 153039725 | 153041931 | *SPRR2D* | Yes | No | -1.1 | 0.58 |
| ENSG00000196805 | small proline rich protein 2B [Source:HGNC Symbol;Acc:HGNC:11262] | 153070224 | 153070840 | *SPRR2B* | Yes | Yes | -0.37 | -0.19 |
| ENSG00000203785 | small proline rich protein 2E [Source:HGNC Symbol;Acc:HGNC:11265] | 153093135 | 153106184 | *SPRR2E* | Yes | Yes | -0.81 | -0.39 |
| ENSG00000244094 | small proline rich protein 2F [Source:HGNC Symbol;Acc:HGNC:11266] | 153112114 | 153113515 | *SPRR2F* | Yes | Yes | 1.5 | 2.1 |
| ENSG00000229035 | small proline rich protein 2C (pseudogene) [Source:HGNC Symbol;Acc:HGNC:11263] | 153140491 | 153140709 | *SPRR2C* | Yes | Yes | -0.29 | 1.5 |
| ENSG00000229699 | lncRNA | 153174518 | 153191676 | *AL161636.1* | Yes | No | -0.24 | 0.35 |
| ENSG00000203782 | loricrin [Source:HGNC Symbol;Acc:HGNC:6663] | 153259700 | 153262122 | *LOR* | Yes | No | -0.091 | -0.12 |
| ENSG00000159527 | peptidoglycan recognition protein 3 [Source:HGNC Symbol;Acc:HGNC:30014] | 153297862 | 153310718 | *PGLYRP3* | Yes | Yes | -0.47 | 0.65 |
| ENSG00000163218 | peptidoglycan recognition protein 4 [Source:HGNC Symbol;Acc:HGNC:30015] | 153330120 | 153348840 | *PGLYRP4* | Yes | Yes | -0.82 | 1.2 |
| ENSG00000143556 | S100 calcium binding protein A7 [Source:HGNC Symbol;Acc:HGNC:10497] | 153457744 | 153460701 | *S100A7* | Yes | No | -0.35 | 0.3 |
| ENSG00000188643 | S100 calcium binding protein A16 [Source:HGNC Symbol;Acc:HGNC:20441] | 153606886 | 153613145 | *S100A16* | Yes | Yes | -0.33 | 0.43 |
| ENSG00000189171 | S100 calcium binding protein A13 [Source:HGNC Symbol;Acc:HGNC:10490] | 153618787 | 153631360 | *S100A13* | Yes | Yes | -0.55 | 0.2 |
| ENSG00000160678 | S100 calcium binding protein A1 [Source:HGNC Symbol;Acc:HGNC:10486] | 153627926 | 153632039 | *S100A1* | Yes | Yes | -0.75 | 0.69 |
