## SupplementalTable16 for "An enhancer:involucrin regulatory module impacts human skin barrier adaptation out-of-Africa and modifies atopic dermatitis risk"

**Table S16. Allele-Specific Amplicons.**

| *Target* | *F Primer* | *R Primer* | *Length* |
|---|---|---|---|
| *Ivl* | TGGGTCAGTCACTTAAGCAAGA | TTCTGCTGCTGCTTCTCTGT | 260 |
| *m2210017l01Rik* | GGTCCCCAGGTTCCTACTTC | TCAAAGCTTATCCTGGGCCA | 267 |
| *Lce6a* | TCCAGAACACTGTCAGCCAT | GCACCATGATCAATTTTATTGTTG | 263 |

| *Amplicon* | *Sequence* | *rs ID* | *C57B6* | *BALB* |
|---|---|---|---|---|
| *Ivl* | TGGGTCAGTCACTTAAGCAAGAGA<br>AAGCTTCAAGG**A**AACAGCAGCTAG<br>ATTACTCACATCTAGAACAGGAGAA<br>GGAGCTCTCAGACCAGCCACTGGA<br>TCAAGCACTAGTAAAGAAGGGTAA<br>ACAACTGGAAAGGAAGAAACACGA<br>ATTGGAGAACCGGACACAGCAGGA<br>GAAGTAGatagagcaattagtaccaagcactg<br>actaagccagtccaaccagtgaaaggagacg**t**g<br>ctcactacagagaagcagcagcagaa | rs32990753 | A | G |
| | | rs32990750 | T | C |
| *2210017l01Rik* | GGTCCCCAGGTTCCTACTTCATGT<br>CTCCCTCTCAGTGCCCCACCAGCT<br>CCTGCTTGCTGTGTTTCTACCTGCT<br>ATATTTCTGG**T**TTGGGAAGCAGCT<br>GCTCTTTAATATCTCACCGATTTCC<br>TCGGTTCTACCTCCGCCAGCCTCA<br>G**A**GTTCTGAGTGTCCTGAGAAAGA<br>GGCTGCAGAATGTTCGAGCTGTTG<br>CCACAACCCTGGAAACTGTAGCTA<br>Aactgcatccctcgagaaaacaaagaacatggc<br>ccaggataagctttga | rs32991827 | T | G |
| | | rs32991824 | A | G |
| *Lce6a* | TCCAGAACACTGTCAGCCATAAGG<br>AAATTCATCACCCACAACCTCGCTG<br>TCTTAGGGGTAGTACCACCTACCA<br>CTGCAAAGAAGAAGAGTGCTAAgaa<br>actgggcacaaacgagggtaaatagctacaaca<br>acctttccagataaactcatgaatt**t**caccaggaag<br>gccaggccctccacctctcttggctagaaaaacatt<br>tcttgtttctttctttagc**c**taccctttcagttcaacaac<br>aataaaattgatcatggtgc | rs31222976 | T | C |
| | | rs31417097 | C | T |
