## SupplementalTable17 for "An enhancer:involucrin regulatory module impacts human skin barrier adaptation out-of-Africa and modifies atopic dermatitis risk"

**Table S17. ATAC-seq library statistics.**

| *Sample* | *Sex* | *Total Reads pairs* | *Post filtering read pairs* | *Alignment Rate* | *Library complexity* | | | *NFR?* | *mono-nucleosome peak?* | *FRiP* | *TSS enrichment* |
|---|---|---|---|---|---|---|---|---|---|---|---|
| | | | | | *NFR* | *PBC1* | *PBC2* | | | | |
| WT_1 | M | 44,104,640 | 39,024,826 | 79.13 | 0.829 | 0.835 | 6.167 | ✔ | ✔ | 0.036 | 8.214 |
| WT_3 | F | 29,518,179 | 32,720,098 | 95.79 | 0.856 | 0.864 | 7.539 | ✔ | ✔ | 0.052 | 9.760 |
| WT_4 | F | 33,796,509 | 36,502,990 | 95.37 | 0.831 | 0.831 | 6.434 | ✔ | ✔ | 0.059 | 11.093 |
| Del_1 | M | 58,966,233 | 28,521,337 | 92.66 | 0.760 | 0.765 | 4.254 | ✔ | ✔ | 0.088 | 14.844 |
| Del_2 | M | 43,934,578 | 21,294,165 | 96.73 | 0.779 | 0.786 | 4.774 | ✔ | ✔ | 0.026 | 6.913 |
| Del_3 | M | 26,577,482 | 14,366,531 | 96.92 | 0.842 | 0.859 | 7.341 | ✔ | ✔ | 0.106 | 14.729 |
| Del_4 | M | 26,880,711 | 14,383,001 | 97 | 0.820 | 0.837 | 6.303 | ✔ | ✔ | 0.092 | 13.897 |
| Large_1 | M | 74,101,870 | 39,840,266 | 99.46 | 0.837 | 0.841 | 6.105 | ✔ | ✔ | 0.033 | 7.750 |
| Large_2 | M | 60,641,676 | 39,482,578 | 99.43 | 0.962 | 0.967 | 31.580 | ✔ | ✔ | 0.042 | 9.260 |
| Large_4 | M | 47,393,679 | 39,766,656 | 82 | 0.799 | 0.809 | 5.320 | ✔ | ✔ | 0.027 | 7.650 |

*Reproducibility*

| | *IDR Values* | | *naïve overlap peaks* | *IDR peaks* |
|---|---|---|---|---|
| **Per Genotype** | **Rescue** | **self consistency** | | |
| WT | 1.1506 | 1.3738 | 169158 | 78032 |
| 923del/del | 1.3075 | 1.5896 | 126285 | 51601 |
| 923 large/large | 1.2698 | 2.0893 | 190190 | 98605 |
